# Molecular and circuit determinants in the globus pallidus mediating control of cocaine-induced behavioral plasticity

**DOI:** 10.1101/2024.05.29.596557

**Authors:** Guilian Tian, Katrina Bartas, May Hui, Lingxuan Chen, Jose J. Vasquez, Ghalia Azouz, Pieter Derdeyn, Rían W. Manville, Erick L. Ho, Amanda S. Fang, Yuan Li, Isabella Tyler, Vincent Setola, Jason Aoto, Geoffrey W. Abbott, Kevin T. Beier

## Abstract

The globus pallidus externus (GPe) is a central component of the basal ganglia circuit, receiving strong input from the indirect pathway and regulating a variety of functions, including locomotor output and habit formation. We recently showed that it also acts as a gatekeeper of cocaine-induced behavioral plasticity, as inhibition of parvalbumin-positive cells in the GPe (GPe^PV^) prevents the development of cocaine-induced reward and sensitization. However, the molecular and circuit mechanisms underlying this function are unknown. Here we show that GPe^PV^ cells control cocaine reward and sensitization by inhibiting GABAergic neurons in the substantia nigra pars reticulata (SNr^GABA^), and ultimately, selectively modulating the activity of ventral tegmental area dopamine (VTA^DA^) cells projecting to the lateral shell of the nucleus accumbens (NAcLat). A major input to GPe^PV^ cells is the indirect pathway of the dorsomedial striatum (DMS^D^^2^), which receives DAergic innervation from collaterals of VTA^DA^→NAcLat cells, making this a closed-loop circuit. Cocaine likely facilitates reward and sensitization not directly through actions in the GPe, but rather in the upstream DMS, where the cocaine-induced elevation of DA triggers a depression in DMS^D^^2^ cell activity. This cocaine-induced elevation in DA levels can be blocked by inhibition of GPe^PV^ cells, closing the loop. Interestingly, the level of GPe^PV^ cell activity prior to cocaine administration is correlated with the extent of reward and sensitization that animals experience in response to future administration of cocaine, indicating that GPe^PV^ cell activity is a key predictor of future behavioral responses to cocaine. Single nucleus RNA-sequencing of GPe cells indicated that genes encoding voltage-gated potassium channels KCNQ3 and KCNQ5 that control intrinsic cellular excitability are downregulated in GPe^PV^ cells following a single cocaine exposure, contributing to the elevation in GPe^PV^ cell excitability. Acutely activating channels containing KCNQ3 and/or KCNQ5 using the small molecule carnosic acid, a key psychoactive component of *Salvia rosmarinus* (rosemary) extract, reduced GPe^PV^ cell excitability and also impaired cocaine reward, sensitization, and volitional cocaine intake, indicating its potential as a therapeutic to counteract psychostimulant use disorder. Our findings illuminate the molecular and circuit mechanisms by which the GPe orchestrates brain-wide changes in response to cocaine that are required for reward, sensitization, and self-administration behaviors.

## INTRODUCTION

VTA^DA^ cells serve a central role in motivation, being critical for a variety of reward- and aversion-related behaviors^1–3^. They also play a critical role in pathological behaviors, including substance abuse. The ventral midbrain DA system has been implicated in all phases of drug abuse, from the initial rewarding effect to withdrawal and ultimately compulsive drug seeking^4–8^. However, it is precisely because the DA system is central to so many functions that it has proved to be a poor target to combat substance abuse; drugs that modulate DA levels or DA receptor function have been largely ineffective in mitigating the impact of addiction. Rather than modulating the DA system as a whole, modulating signaling in particular DA subcircuits may provide therapeutic efficacy while minimizing detrimental off-target effects.

While distinct subpopulations of VTA^DA^ cells play different roles in motivated behaviors, the lack of unique genetic markers has made targeting these subtypes difficult^9–13^. In the absence of a reliable way to target these DA cell subpopulations, another approach would be to target the circuits in which these subpopulations are embedded. We recently successfully applied this approach to leverage circuits controlling cocaine withdrawal. In that study, we first identified a key role of VTA^DA^ cells projecting to the amygdala (VTA^DA^→amygdala) in mediating anxiety-like behaviors induced by withdrawal from repeated cocaine administration^14^. We then performed rabies virus (RABV)-based circuit mapping in saline- and cocaine-treated mice to map which inputs onto VTA^DA^→amygdala cells may be modified by cocaine, finding an elevation in RABV-labeled inputs from GABAergic neurons in the BNST (BNST^GABA^). This elevation in input labeling, which corresponds to an increase in spontaneous activity, in turn contributes to the development of cocaine withdrawal-induced anxiety-like behavior and reinstatement through disinhibition of VTA^DA^→amygdala cells^14^. That work demonstrated the role of a key subcircuit centered on VTA^DA^ cells that contributes to some of the later stages of substance abuse, namely drug withdrawal and reinstatement. Here we aim to map circuits that control the earliest stages of substance abuse through facilitating drug reward and volitional drug intake.

Many brain regions that contribute to the processing of natural and drug rewards have been intensively studied, including the nucleus accumbens (NAc), medial prefrontal cortex (mPFC), and amygdala^5,15–17^. However, there are likely additional key circuits that have not been identified or are poorly understood. For example, we recently showed that the GPe functions as a critical gate for drug-induced behavioral plasticity, including conditioned place preference (CPP) and locomotor sensitization^18^, a relatively surprising discovery given that the GPe is more often implicated in habit formation and motor behavior than substance abuse. Using RABV retrograde circuit mapping, we found that the number of RABV-labeled GPe neurons that innervate VTA^DA^ neurons significantly increased after cocaine exposure. A long-lasting increase in the activity of GPe parvalbumin-positive neurons (GPe^PV^) was observed one day following cocaine administration, as evidenced both by an elevation in intrinsic neuronal excitability as well as an increase in the spontaneous activity *in vivo*. Importantly, we provided evidence that GPe^PV^ cells play a critical role in the development of cocaine-induced behavioral changes as inhibition of GPe^PV^ cells prevented cocaine reward and sensitization. Although we observed a striking increase in the number of VTA^DA^-projecting GPe^PV^ neurons after cocaine exposure, the strength of the synapses made by GPe^PV^ cells onto GABA cells in the substantia nigra pars reticulata (SNr^GABA^) was much stronger than those made directly onto VTA^DA^ cells. Thus, we hypothesized that cocaine-induced increases in GPe^PV^ cell activity leads to a net disinhibition of VTA^DA^ cells, which promotes cocaine-induced behavioral changes such as reward and sensitization.

One limitation of that study is that we did not comprehensively explore how the GPe engages broader brain circuitry to mediate these behaviors. For example, we did not explore which VTA^DA^ cells were principally controlled by GPe cells, or how inputs to GPe^PV^ cells may be modified by cocaine. It also is unknown if the change in GPe^PV^ cell activity following cocaine is due to a change in inputs, a change in gene expression in GPe^PV^ cells, or both. Lastly, it remains unknown whether targeting the GPe and its input/output connections can impact volitional drug taking. One major factor limiting the translatability of our previous findings is that there are no known methods to selectively inhibit GPe cells non-invasively, limiting therapeutic potential in humans. Small molecules that dampen the activity of GPe^PV^ neurons with minimal side-effects may have the potential to reduce cocaine reward and potentially self-administration. In this manuscript, we explore these questions in detail, and report the identification of a natural compound, derived from *Salvia rosmarinus* (rosemary), to combat cocaine abuse.

## RESULTS

### GPe^PV^ cells gate CPP and locomotor sensitization through disinhibition of VTA^DA^→NAcLat cells

As VTA^DA^ neurons can be classified based on their functionally-relevant projection specificity to distinct cortical and subcortical brain regions, our first goal was to define which projection-specific populations of VTA^DA^ cells mediate the GPe’s effect on cocaine-induced behavioral changes. To accomplish this, we systematically chemogenetically inhibited the activity of ventral midbrain-projecting GPe^PV^ cells (GPe^PV^→ventral midbrain) as well as populations of projection-specific VTA^DA^ neurons. To validate our previous findings that chemogenetic inhibition of GPe^PV^→ventral midbrain cells blocked the development of cocaine CPP and sensitization, we injected an adeno-associated virus (AAV) expressing a Cre-dependent chemogenetic inhibitor, hM4Di (AAV-FLEx^loxP^-hM4Di), or YFP as a control into the GPe of PV-Cre mice. One month later, we injected microspheres in the ventral midbrain that release CNO over the course of a week^14,18,19^. Consistent with our previous study, inhibition of GPe^PV^→ventral midbrain cells blocked CPP and sensitization following injections of 15mg/kg cocaine (Figure S1A-F). Although not tested in our original study, we also measured the time spent in the open arms of the elevated plus maze and center of the open field during withdrawal following repeated cocaine administration. Unlike CPP and sensitization, inhibition of GPe^PV^→ventral midbrain cells did not impact these behaviors (Figure S1G-I). To ensure that the microspheres were working as expected, we validated their inhibition of hM4Di-expressing cells using both slice electrophysiology and cFos labeling (Figure S1J-R). Interestingly, optogenetic activation of GPe^PV^ projections to the ventral midbrain was neither reinforcing nor locomotor activating (Figure S1S-Y), indicating that GPe^PV^→ventral midbrain cells serve as a gate of cocaine GPe^PV^-induced plasticity rather than promoting reward or locomotion *per se*.

We next assessed whether discrete populations of projection-specific VTA^DA^ cells are controlled by GPe^PV^ cells. Given that GPe^PV^→ventral midbrain cell activation leads to a net activation of DA cells^18^, if DA cells are being controlled by GPe^PV^ cells, inhibition of either population should have a similar effect. We assessed this by chemogenetically inhibiting different ventral midbrain DA cell populations and testing which had similar effects to inhibition of the GPe^PV^→ventral midbrain projection. To inhibit select subpopulations of ventral midbrain DA cells in mice expressing Cre in DA neurons (DAT-Cre), we injected a retrograde canine adenovirus expressing a Cre-dependent Flp recombinase (CAV-FLEx^loxP^-Flp) into the terminal field of a subset of ventral midbrain DA neurons^18,20,21^. We chose five forebrain sites: NAcLat, NAcMed, dorsolateral striatum (DLS), central amygdala, and mPFC. In the same surgery, we injected an AAV expressing a Flp-dependent hM4Di (AAV-FLEx^FRT^-hM4Di) or AAV-FLEx^FRT^-YFP as a control, into the VTA. This strategy enables specific gene expression in projection-defined DA cells^21^. Notably, there is minimal collateralization between these five subpopulations^21,22^. Clozapine-N-oxide (CNO; 5 mg/kg) was administered intraperitoneally (i.p.) 30 minutes before each administration of cocaine as well as the saline pairing in the CPP task to inhibit the output-defined subpopulation of DA cells (Figure 1A, representative images shown in Figure 1B). Importantly, of the five brain regions targeted, only inhibition of VTA^DA^→NAcLat cells during cocaine administration prevented both reward and sensitization (Figure 1C-V) with no effect on the time spent in the open arms of the elevated plus maze, or center of the open field (Figure S2A-C). These results indicate that GPe^PV^→ventral midbrain cells and VTA^DA^→NAcLat cells effect the same cocaine-induced behavioral changes.

**Figure 1:**
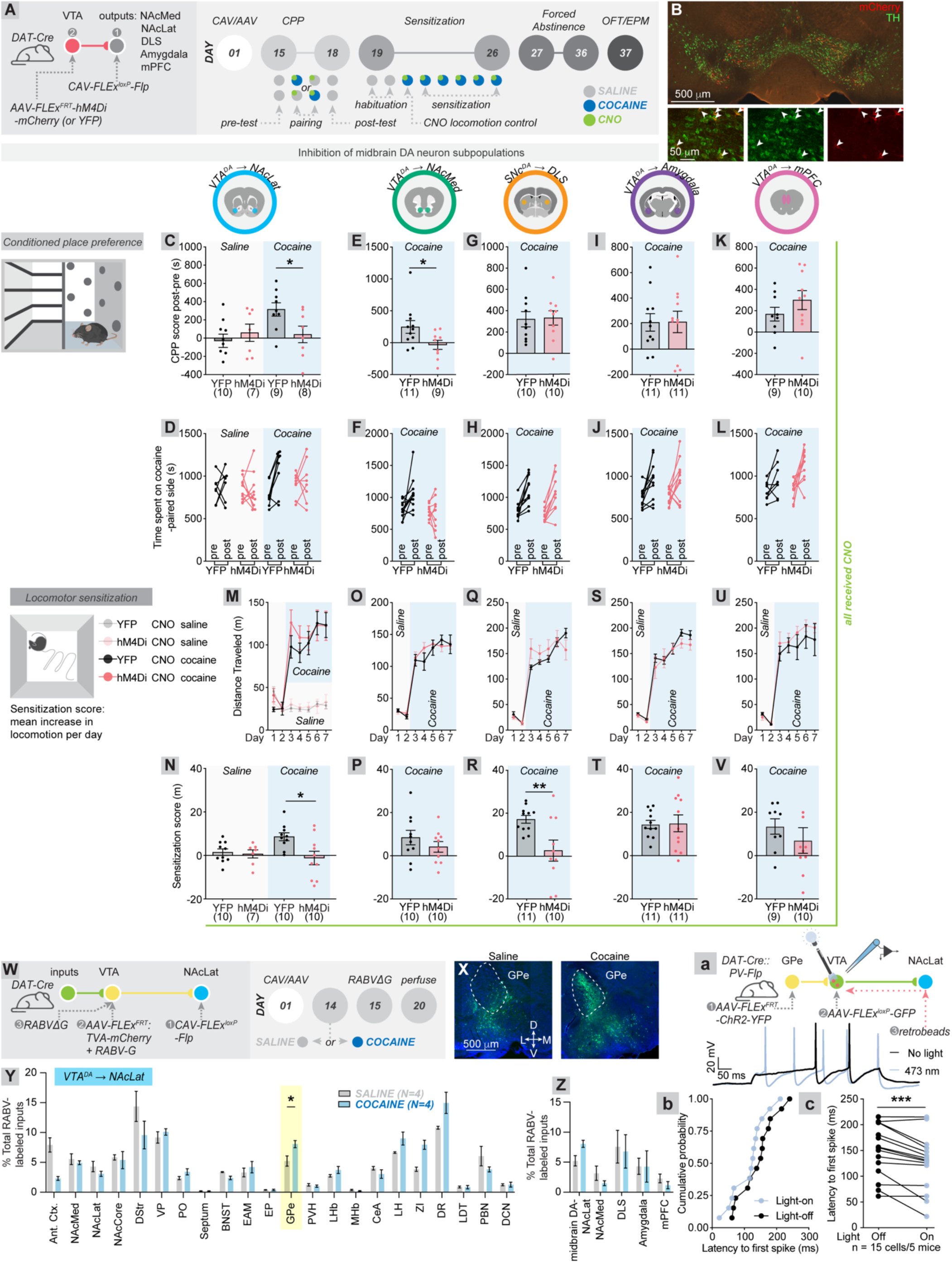
Linking the GPe and ventral midbrain DA cells. (A) Schematic of DREADD inhibition experiments of five projection-defined ventral midbrain DA neuron subpopulations. 1) CAV-FLEx^loxP^-Flp was injected into one of five DA neuron target sites. Then, 2) AAV-FLEx^FRT^-hM4Di-mCherry, or AAV-FLEx^FRT^-YFP as a control was injected into the VTA/SNc. Two weeks later, mice were tested for cocaine CPP, sensitization, and withdrawal anxiety-related behavior (15 mg/kg cocaine). CNO 5 mg/kg was given i.p. 30 minutes before each cocaine administration. (B) Sample image of the ventral midbrain of a mouse where CAV-FLEx^loxP^-Flp was injected into the NAcLat and AAV-FLEx^FRT^-hM4Di-mCherry into the VTA. Green = tyrosine hydroxylase (TH), a marker for DA neurons. Red = mCherry. Bottom, high magnification images of overlap between mCherry and TH. (C-L) CPP scores and time spent in each chamber for each group of animals with saline or cocaine, and hM4Di or YFP treatments. No CPP was observed in saline-treated mice (C), whether or not VTA^DA^→NAcLat cells were inhibited during both side pairings. In cocaine-treated mice, hM4Di-mediated inhibition of VTA^DA^→NAcLat (C-D) and VTA^DA^→NAcMed (E-F) cells prevented CPP, whereas inhibition of SNc^DA^→DLS (G-H), VTA^DA^→amygdala (I-J) or VTA^DA^→mPFC (K-L) cells had no effect (C: One-way ANOVA p = 0.0065; pairwise t-tests with multiple comparisons corrections, YFP saline vs. YFP cocaine, -29.7s vs. 365.5s, p = 0.0054; YFP saline vs. hM4Di saline, -29.7s vs. 59.4s, p = 0.88; YFP saline vs. hM4Di cocaine, -29.7s vs 67.7, p = 0.81; YFP cocaine vs. hM4Di cocaine, 365.5s vs. 67.7s, p = 0.049. Note that one data point (YFP saline) is below the y-axis cutoff. E: YFP vs hM4Di, 266.4 vs. -42.2, p = 0.02; G: YFP vs hM4Di, 320.0s vs. 332.7s, p = 0.90; I: YFP vs hM4Di, 209.5s vs. 214.0s, p = 0.97; K: YFP vs hM4Di, 167.8s vs. 299.2s, p = 0.26). Numbers of mice for each group are reported below the x-axis. (M-V) Sensitization scores for each group of animals with saline or cocaine, and hM4Di or YFP treatments. No sensitization was observed in saline-treated mice (M), whether or not VTA^DA^→NAcLat cells were inhibited during testing in the open field. hM4Di-mediated inhibition of VTA^DA^→NAcLat (M-N) or SNc^DA^→DLS (Q-R) cells prevented locomotor sensitization, whereas inhibition of VTA^DA^→NAcMed (O-P), VTA^DA^→amygdala (S-T) or VTA^DA^→mPFC (U-V) cells had no effect (N: One-way ANOVA p = 0.016; pairwise t-tests with multiple comparisons corrections, YFP saline vs. YFP cocaine, 1.5 m vs. 8.6 m, p = 0.11; YFP saline vs. hM4Di saline, 1.5 m vs. 0.7 m, p = 1.0; YFP saline vs. hM4Di cocaine, 1.5 m vs -1.2 m, p = 0.82; YFP cocaine vs. hM4Di cocaine, 8.6 m vs. -1.2 m, p = 0.014. P: YFP vs hM4Di, 8.5 m vs. 4.1 m, p = 0.31; R: YFP vs hM4Di, 17.3 m vs. 2.7 m, p = 0.009; T: YFP vs hM4Di, 14.5 m vs. 15.0 m, p = 0.91; V: YFP vs hM4Di, 13.4 m vs. 7.0 m, p = 0.37. Note that in panel V, 1 data point (hM4Di) is located above the y-axis cutoff). (W) Schematic of cTRIO experiments from VTA^DA^→NAcLat cells. CAV-FLEx^loxP^-Flp was injected into NAcLat, and a mixture of AAV-FLEx^FRT^-TVA-mCherry (TC) and AAV-FLEx^FRT^-RABV-G was injected into the VTA of DAT-Cre mice. Thirteen days later, mice received a single injection of 15 mg/kg cocaine, or saline. EnvA-pseudotyped RABVΔG was injected into the VTA on day 15. Experiments were terminated on day 20. (X) Representative images of the GPe for cTRIO experiments onto VTA^DA^→NAcLat cells in saline- and cocaine-treated mice. The total number of labeled input cells for these two brains were approximately equivalent. Blue = DAPI, green = RABV-labeled input cells. (Y) Percentage of total RABV-labeled inputs in 22 different input sites. Two-way ANOVA drug effect p > 0.99, interaction effect p < 0.0001. Focusing on GPe inputs specifically, labeled inputs from GPe in saline- vs. cocaine-treated mice, 5.2% of total inputs labeled in saline-treated vs. 8.1% of inputs in cocaine-treated mice, p = 0.03 (no multiple comparisons corrections), n = 4 each. (Z) Percentage of RABV-labeled inputs in the GPe for cTRIO experiments from VTA^DA^→NAcLat, VTA^DA^→NAcMed, SNc^DA^→DLS, VTA^DA^→amygdala, and VTA^DA^→mPFC cells, in mice treated with either saline or cocaine. n = 4 each. Labeled GPe inputs were only elevated onto VTA^DA^→NAcLat cells (p = 0.03), and not other DA cell populations. (a) Schematic of slice electrophysiology experiments testing the effect of GPe^PV^ cell stimulation on VTA^DA^→NAcLat cell activity. AAV-FLEx^FRT^-ChR2-YFP was injected into the GPe, AAV-FLEx^loxP^-GFP was injected into the VTA, and red retrobeads were injected into the NAcLat of DAT-Cre::PV-Flp mice. Acute slice preparations were prepared one month later, and recordings were performed in current clamp from GFP+, retrobead+ cells in the VTA, with or without 20 Hz stimulation of 473 nm light. Shown below are representative traces from the same VTA^DA^→NAcLat cell with (blue) and without (black) 20 Hz optogenetic stimulation of GPe^PV^ fibers in slice. (b) Cumulative probability plot for the latency to first spike. (c) Latency to first spike from the same cells, with light off and light on. Latency with light off, 149 ms vs. latency with light on, 126 ms, p = 0.0007.

A single cocaine exposure triggers a long-lasting elevation in spontaneous activity in GPe^PV^ cells^18^. We hypothesize that since GPe^PV^→ventral midbrain cells and VTA^DA^→NAcLat cells mediate the same behaviors in response to cocaine, the elevation in GPe^PV^ cell activity will impact VTA^DA^→NAcLat cells, but not other VTA^DA^ cells. To test this hypothesis, we performed RABV mapping experiments from each of the five subtypes of ventral midbrain DA neurons using cell type-specific Tracing the Relationships between Inputs and Output (cTRIO^20,21^) and assessed whether the proportion of labeled GPe inputs was modified or not by a single administration of cocaine. To do this, we injected a canine adenovirus expressing the Flp recombinase in cells expressing Cre (CAV-FLEx^loxP^-Flp) into the NAcLat, NAcMed, DLS, amygdala, or mPFC. During the same surgery, we injected a combination of AAVs expressing TVA (AAV-FLEx^FRT^-TVA-mCherry), which mediates infection by EnvA-pseudotyped viruses, and the RABV glycoprotein (AAV-FLEx^FRT^-RABV-G), which facilitates RABV spread to connected input cells, into the VTA or SNc. Both AAVs were expressed specifically in cells expressing the Flp recombinase. Thirteen days later, a single dose of saline or cocaine was given to each mouse. The following day, a glycoprotein-deleted, EnvA-pseudotyped, GFP-expressing rabies virus (RABVΔG) was injected into the VTA or SNc, and animals were sacrificed five days later (Figure 1W, representative images in Figure 1X). Of the 22 distinct quantified brain regions identified by cTRIO that innervate VTA^DA^→NAcLat cells, only inputs from four brain regions were elevated following a single cocaine exposure, including from the GPe, which is consistent with our hypothesis (Figure 1Y). Furthermore, the percentage of labeled GPe inputs onto other VTA^DA^ cell populations were not significantly affected by cocaine (Figure 1Z). In addition, this elevation was specific to rewarding experiences such as cocaine administration, as aversive foot shocks elicited during an auditory fear conditioning protocol did not trigger an elevation in GPe input labeling onto VTA^DA^→NAcLat cells (Figure S2F-G). These experiments together link GPe^PV^ cells and VTA^DA^→NAcLat cells and suggest that VTA^DA^→NAcLat cells likely effect behavioral changes induced by the GPe.

Notably, given that the mechanisms of RABV cell-cell transmission are not well understood, we treat our results as a screen to identify input populations that were likely influenced by cocaine. We therefore do not heavily weight statistical significance in this assay, but only use it to select input sites for further study. Importantly, one known caveat to this RABV mapping approach is that the extent of input labeling onto a defined cell population cannot be used as the sole method to define functional properties of connections; labeled inputs can be elevated onto multiple cell populations in a given brain region, and the effect of these changes on the firing of each cell type may not be uniform. For example, even though a single cocaine administration elevates the proportion of labeled GPe inputs onto VTA^DA^ cells, it also elevates the proportion of labeled GPe inputs onto nearby ventral midbrain GABA cells^18^. Due to this limitation in interpreting the functional consequences of RABV input labeling on the functional output of downstream cell populations, it is critical to use orthogonal approaches to define the functional connectivity relationships between connected cells. To examine if activation of GPe^PV^ cells inhibits or disinhibits VTA^DA^→NAcLat cells, we injected AAV-FLEx^FRT^-ChR2-YFP into the GPe, AAV-FLEx^loxP^-GFP into the VTA, and red retrobeads into the NAcLat of PV-Flp::DAT-Cre mice and performed whole-cell recordings from acute brain slices one month later. We patched onto GFP+, retrobead-containing cells (VTA^DA^→NAcLat) in current clamp and assessed the effects of 20 Hz optical stimulation on the latency to the first action potential (Figure 1a). We observed that optogenetic activation of GPe^PV^ terminals reduced the latency to the first action potential, indicating that the effect of GPe^PV^ stimulation on VTA^DA^→NAcLat cell activity was indeed disinhibitory, as expected (Figure 1b-c).

### Elevation in GPe^PV^ cell activity occurs through cocaine-triggered DA release in VTA^DA^→NAcLat collaterals in the dorsomedial striatum (DMS)

The GPe is an *input* to VTA^DA^ cells; inhibition of the GPe^PV^→ventral midbrain projection blocks cocaine CPP and sensitization through control of VTA^DA^ cells^18^. However, cocaine acts by increasing DA levels in the synapse; thus, its effects are likely mediated by a brain region receiving significant innervation from DA cells. There is no appreciable expression of the D_1_ DA receptor-encoding mRNA (*Drd1*) in the GPe, and only a small number of GPe cells express the D_2_ DA receptor-encoding mRNA (*Drd2*) (Figure S3A-B). However, the GPe receives innervation from both *Drd1*-expressing and *Drd2-*expressing cells, with the input from *Drd2*-expressing cells being much stronger (Figure S3C). Given that few GPe cells express DA receptors but the GPe receives strong innervation from cells that do, a key question is: where in the brain does DA act to facilitate the persistent elevation in GPe^PV^ cell activity? The NAc, including the NAcLat, does not project to the GPe. To identify the relevant connection(s), we first asked the question: which DA cells send projections to brain sites expressing DA receptors that in turn project to the GPe? To answer this question, we injected CAV-FLEx^loxP^-Flp into the NAcLat, NAcMed, DLS, amygdala, or mPFC, AAV-FLEx^FRT^-mGFP into the VTA or SNc and waited 2 months for viral expression. This resulted in mGFP being expressed in subpopulations of ventral midbrain DA neurons, labeling their entire axonal tree (Figure 2A-B). Of the DA neuron populations targeted, only VTA^DA^→NAcLat and SNc^DA^→DLS cells showed any appreciable projection to the GPe, with the projection from VTA^DA^→NAcLat cells being quantitatively larger (Figure 2C-D). However, the projections to the GPe were minor; a much more substantial collateral exists to the dorsomedial striatum (DMS), in particular from VTA^DA^→NAcLat cells (Figure 2B-D).

**Figure 2:**
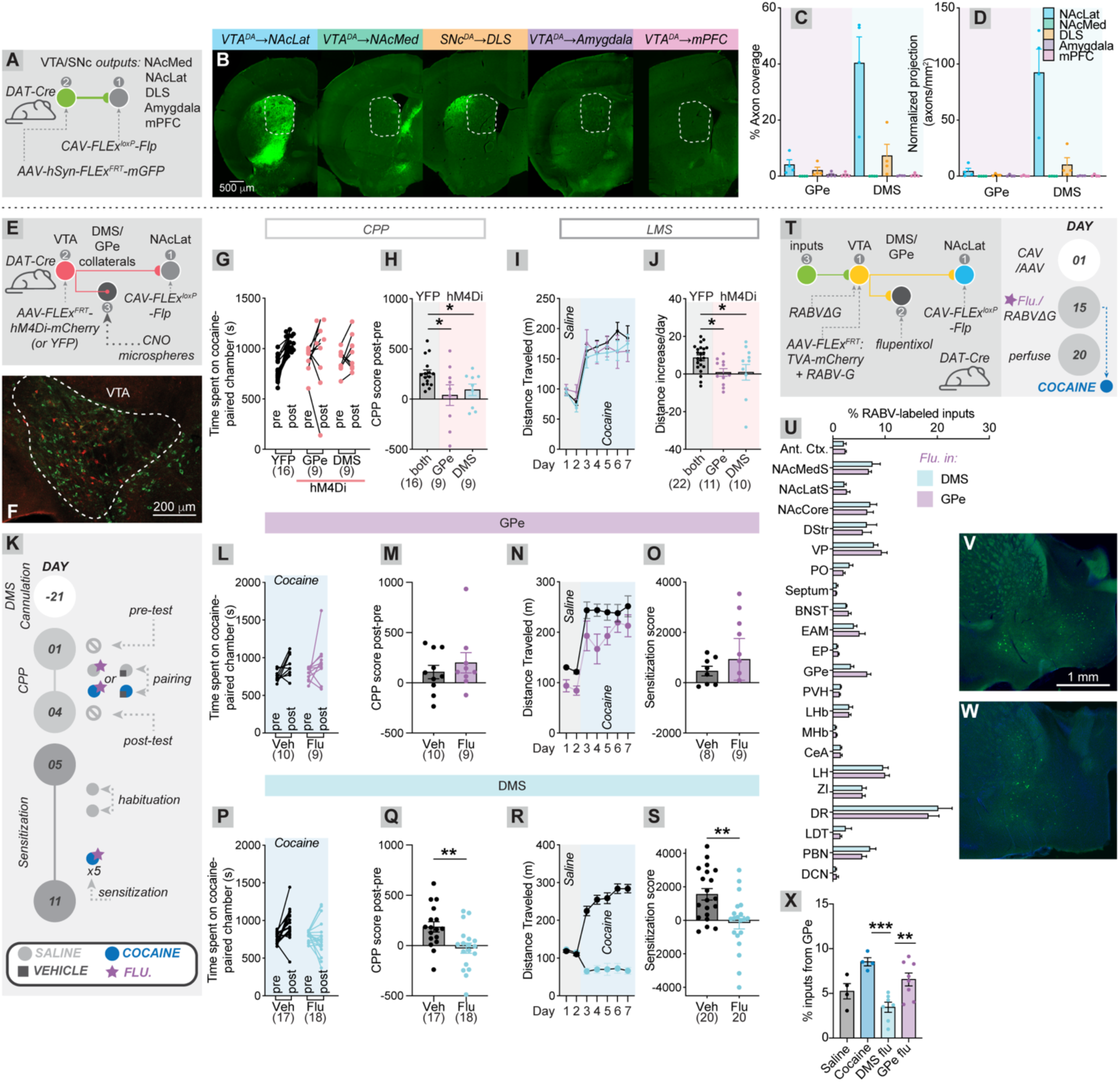
DA release into the DMS, and not GPe, contributes to cocaine-evoked changes in GPe inputs to the ventral midbrain. (A) Schematic of experiments of output mapping of five projection-defined ventral midbrain DA neuron subpopulations. CAV-FLEx^loxP^-Flp was injected into one of five ventral midbrain DA neuron target sites, and AAV-FLEx^FRT^-mGFP was injected into the VTA/SNc. Experiments were terminated two months later. (B) Sample images of the DMS for each target injection, indicating that injections targeting the NAcLat resulted in the most GFP-expressing axons terminating in the DMS, consistent with the highly unique projections of each DA cell population and demonstrating specificity of injections. (C) The percentage of each brain region covered by axons from the corresponding ventral midbrain DA neuron subtype. GPe: NAcLat, 4.1%; NAcMed, 0.02%; DLS, 2.4%; amygdala, 0.52%; mPFC, 0.41%. DMS: NAcLat, 40.4%; NAcMed, 0.02%; DLS, 7.9%; amygdala, 0.15%; mPFC, 0.33%. n=4 for each condition. (D) The normalized projections from each subpopulation, normalizing for the numbers of GFP-expressing cells. GPe: NAcLat, 4.4 axons/mm^2^; NAcMed, 0.01 axons/mm^2^; DLS, 1.23 axons/mm^2^; amygdala, 0.53 axons/mm^2^; mPFC, 0.29 axons/mm^2^. DMS: NAcLat, 92.36 axons/mm^2^; NAcMed, 0.04 axons/mm^2^; DLS, 11.9 axons/mm^2^; amygdala, 0.35 axons/mm^2^; mPFC, 0.51 axons/mm^2^. n=4 for each condition. (E) Experimental schematic of chemogenetic inhibition of VTA^DA^→NAcLat collaterals in the GPe or DMS. CAV-FLEx^loxP^-Flp was injected into the NAcLat, and AAV-FLEx^FRT^-hM4Di-mCherry or AAV-FLEx^FRT^- hM4Di-YFP was injected into the VTA. One month later, experiments were initiated, including an injection of CNO microspheres into either the DMS or GPe, following by cocaine injections testing the effects on CPP and locomotor sensitization. The shortened behavioral protocol was used as in Figure S1B. (F) Sample image of the VTA indicating hM4Di expression from VTA^DA^→NAcLat cells. (G-H) Inhibition of VTA^DA^→NAcLat collaterals in the GPe or DMS prevented cocaine CPP. YFP-expressing controls were combined as both control groups showed similar CPP (One-way ANOVA p = 0.033; pairwise t-tests, YFP vs. hM4Di, GPe CNO 256.7 s vs. 39.9 s, p = 0.024; YFP vs. hM4Di, DMS CNO 256.7 s vs. 93.4 s, p = 0.018). (I-J) Inhibition of VTA^DA^→NAcLat collaterals in the GPe or DMS prevented cocaine-induced locomotor sensitization. YFP-expressing controls were combined as both control groups showed similar sensitization (One-way ANOVA, p = 0.023; unpaired t-tests, YFP vs. hM4Di, GPe CNO 8.62 m vs. 0.87 m, p = 0.022; YFP vs. hM4Di, DMS CNO 8.62 m vs. 0.93 m, p = 0.027). (K) Schematic of experiments to test whether infusion of flupentixol into the GPe (L-O) or DMS (P-S) prior to cocaine injection interferes with cocaine-induced behavioral changes. (L-M) Flupentixol administration into the GPe has no effect on cocaine-induced place preference (vehicle vs. flupentixol 107.6 s vs. 199.7 s, p = 0.45). (N-O) Flupentixol administration into the GPe has no effect on cocaine-induced locomotor sensitization (vehicle vs. flupentixol 4.6 m vs. 9.3 m, p = 0.61). One point in the flupentixol-treated group is below the y-axis. (P-Q) Flupentixol administration into the DMS blocks cocaine-induced place preference (vehicle vs. flupentixol 185.6 s vs. -25.2 s, p = 0.0063). (R-S) Flupentixol administration into the DMS blocks cocaine-induced locomotor sensitization (vehicle vs. flupentixol 15.6 m vs. -1.4 m, p = 0.0019). (T) Schematic of experiments to block DA signaling in either the GPe or DMS during cocaine administration and assess effects on GPe input labeling in cTRIO experiments from the VTA^DA^→NAcLat cells. CAV-FLEx^loxP^-Flp was injected into NAcLat, and a mixture of AAV-FLEx^FRT^-TC and AAV-FLEx^FRT^-RABV-G was injected into the VTA of DAT-Cre mice. On Day 15, EnvA-pseudotyped RABV was injected into the VTA, and flupentixol (0.25 μL, 40 mg/mL) was infused into either the DMS or GPe bilaterally. Five minutes after awakening from isoflurane anesthesia, mice received a single injection of 15 mg/kg cocaine. Experiments were terminated on Day 20. (U) Comparison of whole-brain input labeling from VTA^DA^→NAcLat cells when flupentixol was administered into the DMS or GPe. The GPe showed the most significant difference between conditions (Two-way ANOVA p > 0.99; subsequent unpaired t-tests, adjusted p-value GPe p = 0.29; next closest region, DR, p = 0.96). n = 7 DMS flupentixol; n = 8 GPe flupentixol. (V-W) Sample images from the GPe of mice treated with flupentixol in the (V) DMS, (W) GPe. Images were taken from brains with approximately the same number of total RABV-labeled inputs. (V) Percent of total quantified RABV-labeled input cells arising from the GPe in mice not receiving flupentixol infusions (receiving i.p. saline or cocaine), and those receiving flupentixol infusions (receiving i.p. cocaine). One way ANOVA p = 0.0007; pairwise t-tests with Tukey multiple comparisons corrections, DMS flupentixol vs. GPe flupentixol 3.4% vs. 6.6%, p = 0.0099; DMS flupentixol vs. cocaine 3.4% vs. 8.5%, p = 0.0006; GPe flupentixol vs. cocaine 6.6% vs. 8.5%, p = 0.25.

Given the dense expression of both *Drd1* and *Drd2* in the striatum, we hypothesized that perhaps the dorsal striatum, in particular the DMS, may be influenced by cocaine, and DMS cells in turn could affect activity in GPe^PV^ cells. To assess this question, we first wanted to test whether the GPe^PV^ cells that project to the midbrain are innervated more strongly by the DMS or the adjacent DLS. To explore the topology of the dorsal striatum to GPe connection, we first quantified the density of RABV-labeled input cells in the GPe onto VTA^DA^→NAcLat cells, defined using cTRIO, to identify where in the GPe the input cells were located. Most GPe cells providing inputs to VTA^DA^→NAcLat cells are located in the medial GPe, and cocaine preferentially enhanced input labeling from these medially-located GPe cells (Figure S3D). We next assessed the topology of inputs from the indirect pathway that comprise the majority of total GPe inputs^23^ arising from the DMS and DLS using publicly available data from the Allen Mouse Brain Connectivity Atlas^24^. These data showed that DMS inputs have a projection bias onto the medial GPe, whereas DLS inputs have a bias onto the lateral GPe (Figure S3E). These experiments together indicate that the DMS provides stronger input onto the GPe cells that innervate VTA^DA^→NAcLat cells than the DLS.

Given these data, we hypothesized that VTA^DA^→NAcLat cells effect changes in GPe^PV^ cell activity through their collaterals either to DMS^D2^ cells themselves, or, less likely, through their inputs directly to the GPe. To test the function of VTA^DA^→NAcLat collaterals in the DMS or GPe on cocaine reward and sensitization, we expressed hM4Di in VTA^DA^→NAcLat cells as in Figure 1. Instead of administering CNO systemically, we targeted collaterals in the DMS or GPe through local administration of CNO microspheres (Figure 2E-F). The same abridged behavioral testing protocol was used as in Figure S1B. We found that CNO microsphere administration into either the GPe or DMS blocked the development of CPP and sensitization (Figure 2G-J). These results indicate that inhibition of VTA^DA^→NAcLat projections to, or through, these brain sites blocked cocaine CPP and sensitization. Notably, the collaterals of VTA^DA^→NAcLat cells to the DMS pass through the GPe; thus, if the DMS is the critical target, we would anticipate that CNO inhibition of either axons passing through the GPe or terminals in the DMS would have a similar effect. Therefore, while these experiments indicate that activity in these collaterals is necessary for development of CPP and sensitization, they do not unambiguously show the critical site for DA release in mediating CPP and sensitization.

To definitively test this question, we implanted mice with cannula into the DMS or GPe and administered either 250 μL of 40 mg/mL flupentixol, a non-selective DA receptor antagonist, or vehicle as a control into the DMS or GPe. Flupentixol or vehicle administration was given before both saline and cocaine pairings in the CPP task, and before cocaine each day in the locomotor sensitization tests (Figure 2K). Thirty minutes following infusion, mice were injected with cocaine or saline. While flupentixol administration into the GPe only had a moderate effect on overall locomotion but no significant effect on CPP or locomotor sensitization, flupentixol infusion into the DMS completely blocked CPP and locomotor sensitization (Figure 2L-S). These results support the conclusion that cocaine-evoked DA release in the DMS, but not GPe, is critical for CPP and locomotor sensitization. If DA release in the DMS is critical for cocaine-induced CPP and sensitization, and cocaine also increases RABV labeling of GPe^PV^ input cells onto VTA^DA^→NAcLat cells which is important for cocaine CPP and sensitization, we would expect that flupentixol administration into the DMS, but not GPe, would prevent the cocaine-induced elevation in GPe^PV^ cell input labeling onto VTA^DA^→NAcLat cells. To test this hypothesis, we administered flupentixol into either the DMS or GPe fifteen minutes prior to a single i.p. cocaine injection. One day later, we performed cTRIO from VTA^DA^→NAcLat cells and examined if either perturbation prevented the cocaine-induced elevation in GPe input labeling (Figure 2T). Infusion of flupentixol into the DMS completely blocked the cocaine-induced elevation in GPe input labeling, while flupentixol infusion into the GPe had no effect (Figure 2U-X). These results together indicate that VTA^DA^→NAcLat cells trigger the elevation in GPe^PV^ cell activity, and CPP and sensitization, via DA release from their collaterals in the DMS.

### A single cocaine exposure reduces RABV-labeled DMS inputs onto GPe^PV^ cells

An elevation in GPe^PV^ cell activity could be induced by a change in the ratio of excitatory and inhibitory inputs these cells receive, and/or a change in intrinsic cellular excitability. To assess how inputs to GPe^PV^ cells may be impacted by a single cocaine exposure, we performed RABV input mapping onto GPe^PV^ cells following a single exposure to cocaine or saline (Figure 3A). We observed that cocaine reduced the relative labeling of inputs from inhibitory populations, most notably the DMS, and elevated the relative labeling of excitatory inputs (Figure 3B-D). This relative change in input labeling is consistent with an elevation in GPe^PV^ cell activity following a single cocaine exposure. It is important to consider that changes in labeled inputs induced by cocaine may not reflect a change in synapse number but may reflect a change in cellular activity^14,18^, synapse strength, or a combination of these three factors. Given the importance of the indirect pathway control of GPe^PV^ cells, we hypothesized that activation of the DMS indirect pathway (DMS^D2^ cells) should have the same effects on CPP and sensitization as inhibition of GPe^PV^ cells. To test this, we first used a non-pathway-specific method to activate indirect pathway neurons through systemic administration of 0.25 mg/kg CGS 21680, an agonist of A_2A_ receptors that are predominately expressed on indirect pathway neurons in the striatum. This perturbation prevented cocaine CPP and sensitization (Figure S3F-I), consistent with our hypothesis that activation of the indirect pathway should suppress CPP and sensitization. To selectively activate only DMS^D2^ cells, we injected AAV-FLEx^loxP^-hM3Dq, or AAV-DIO-YFP as a control, into the DMS of A2a-Cre mice. Two weeks later, we performed tests of CPP and locomotor sensitization as done previously, with i.p. CNO being administered 30 minutes before each cocaine injection (Figure 3E-F). Activation of DMS^D2^ cells prevented both CPP and sensitization, as expected (Figure 3G-J), indicating that activation of DMS^D2^ cells and inhibition of downstream GPe^PV^ cells have the same behavioral consequences.

**Figure 3:**
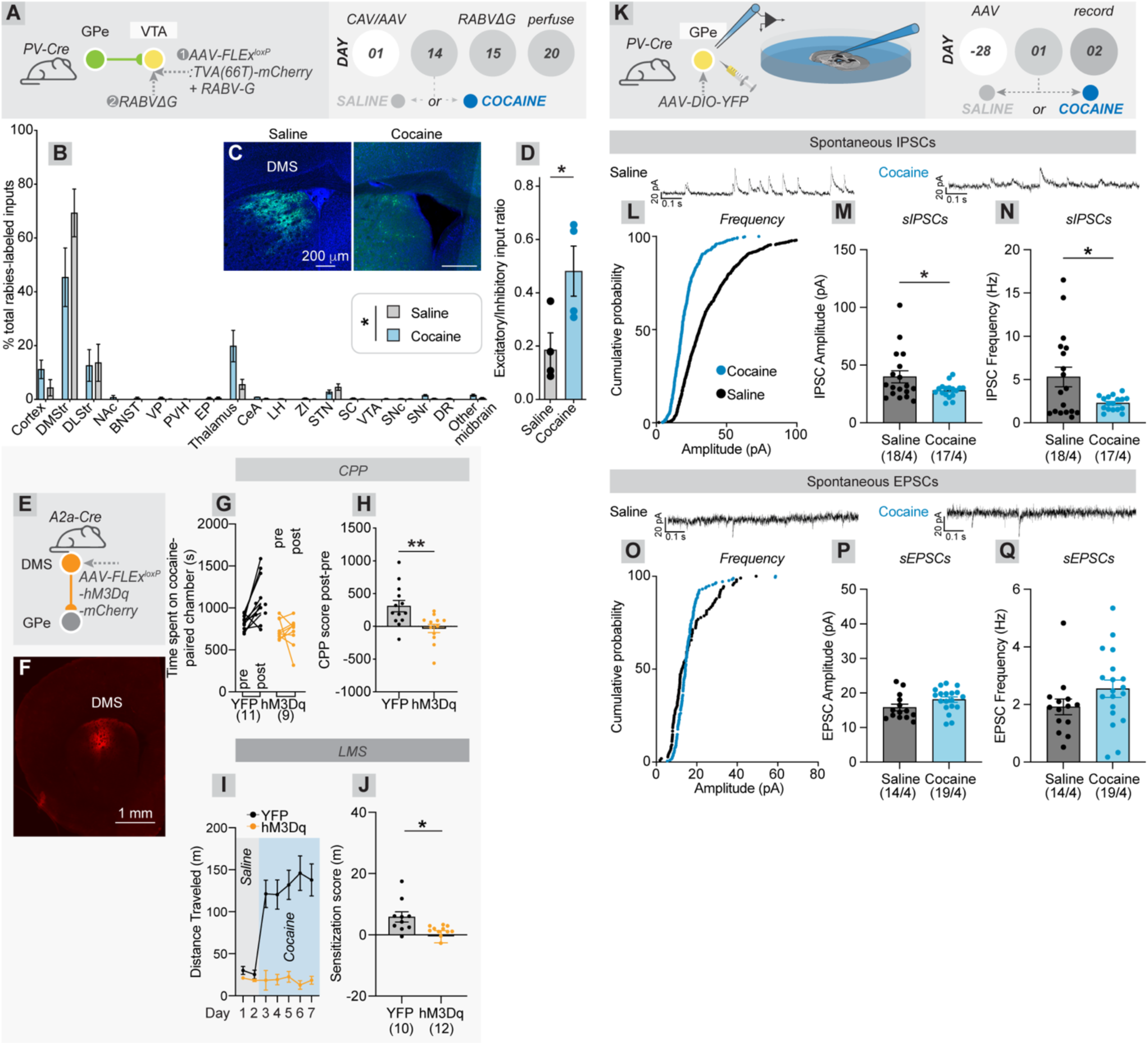
Cocaine alters RABV-mediated input labeling onto GPe^PV^ cells and reduces spontaneous inhibitory input onto GPe^PV^ cells. (A) Experimental schematic of RABV input mapping from GPe^PV^ cells. A mixture of AAV-FLEx^loxP^-TC66T and AAV-FLEx^loxP^-RABV-G was injected into the GPe of PV-Cre mice. Thirteen days later, mice received a single injection of 15 mg/kg cocaine, or saline. EnvA-pseudotyped rabies virus (RABV) was injected into the GPe on Day 15. Experiments were terminated on Day 20. (B) Percentage of total RABV-labeled inputs from 19 different input sites to GPe^PV^ cells. Two-way ANOVA drug effect p >0.99, interaction effect p = 0.006. n = 4 saline, 6 cocaine. (C) Sample images from the DMS from saline- and cocaine-treated mice. (D) The ratio of RABV-labeled inputs from putative excitatory cells (cortex/thalamus/subthalamic nucleus (STN)) to putative inhibitory inputs (DMS/DLS) was higher in cocaine-treated mice. Saline vs. cocaine, 0.19 vs. 0.48, p = 0.04. Two points were excluded from the cocaine group for this analysis as outliers (more than 2 standard deviations outside of the mean, here one above and one below the mean). (E) Experimental schematic of chemogenetic activation of DMS^D2^ cells. AAV-FLEx^loxP^-hM3Dq-mCherry, or AAV-DIO-YFP as a control, was injected into the DMS. Two weeks later, cocaine CPP and sensitization were tested, with CNO being injected i.p. 30 minutes before cocaine. (F) Sample image of hM3Dq-mCherry expression in the DMS of A2a-Cre mice. (G-H) Chemogenetic activation of DMS^D2^ cells prevented cocaine CPP. YFP vs. hM3Dq, 309.4 s vs. - 31.4 s, p = 0.0049. (I-J) Chemogenetic activation of DMS^D2^ cells prevented cocaine-induced locomotor sensitization. YFP vs. hM3Dq, 5.84 m vs. -0.63 m, p = 0.025. (K) Schematic of experiment. AAV-DIO-YFP was injected into the GPe of PV-Cre mice. Four weeks later, mice received a single administration of either saline or 15 mg/kg cocaine. The following day, acute slices were cut, and whole-cell patch clamp recordings were conducted from YFP+ (GPe^PV^) cells, in voltage clamp mode. (L) Cumulative probability plot of the amplitude of individual spontaneous IPSCs in saline- vs. cocaine- treated mice. (M) Comparison of IPSC amplitude in saline- vs. cocaine-treated mice. Saline vs. cocaine, 39.9 pA vs. 28.1 pA, p = 0.040. (N) Comparison of IPSC frequency in saline- vs. cocaine-treated mice. Saline vs. cocaine, 5.31 Hz vs. 2.28 Hz, p = 0.016. (O) Cumulative probability plot of the amplitude of individual spontaneous EPSCs in saline- vs. cocaine- treated mice. (P) Comparison of EPSC amplitude in saline- vs. cocaine-treated mice. Saline vs. cocaine, 15.8 pA vs. 18.1 pA, p = 0.08. (Q) Comparison of EPSC frequency in saline- vs. cocaine-treated mice. Saline vs. cocaine, 1.9 Hz vs. 2.55 Hz, p = 0.14.

We next wanted to assess the effects of cocaine on the input control of GPe^PV^ cells. Given that cocaine increases the activity of GPe^PV^ cells, we hypothesized that this may be due to either an increase in excitatory input, or reduction in inhibitory input. To test these possibilities, we injected AAV-DIO-YFP into the GPe of PV-Cre mice, administered either a single dose of saline or cocaine one month later, and performed voltage-clamp recordings in acute slice preparations the following day from GPe^PV^ cells. In these cells we measured both spontaneous excitatory and inhibitory currents (sEPSCs and sIPSCs, respectively) (Figure 3K). We observed a significant cocaine-evoked reduction in both the frequency as well as the amplitude of spontaneous sIPSCs (Figure 3L-N), with no significant change in the frequency or amplitude of spontaneous sEPSCs (Figure 3O-Q). These results are consistent with a reduction in inhibitory drive onto GPe^PV^ cells, which would contribute to an elevation in GPe^PV^ cell activity.

### DMS^D2^, GPe^PV^, SNr^GABA^, and VTA^DA^→NAcLat cell activity is functionally integrated and carries a common <u>signal</u>

Based on our results, we propose the following circuit mechanism: a single cocaine administration causes DA release from collaterals of VTA^DA^→NAcLat cells in the DMS, which then triggers, likely via G_αi_-coupled DRD2 DA receptor signaling, a reduction in inhibition from DMS^D2^ cells onto GPe^PV^ cells that disinhibits GPe^PV^ cells. Activation of GPe^PV^ cells in turn disinhibits VTA^DA^ cells through suppression of activity in SNr^GABA^ cells^18^ (Figure 4A). If this circuit acts as a closed loop, we hypothesize the following: 1) Activity changes induced by cocaine should influence most or all nodes in the circuit, 2) Activity in each circuit node should similarly reflect aspects of both cocaine CPP and sensitization, and 3) Targeted modulation of activity of each circuit node should impact activity in the others. To assess hypothesis 1, we performed fiber photometry from each of the four nodes (DMS^D2^, GPe^PV^→ventral midbrain, SNr^GABA^, VTA^DA^→NAcLat) while performing the cocaine administration protocol shown in Figure 4B. The GFP-based calcium indicator GCaMP was expressed in each of the four targeted cell populations by injecting AAV-FLEx^loxP^-GCaMP7f into either the DMS in A2a-Cre mice or the VTA in DAT-Cre mice, by injecting AAV-FLEx^FRT^-GCaMP6f into the SNr of vGAT-Flp mice, or by injecting AAV_rg_-DIO-FLPo in the ventral midbrain and AAV-FLEx^FRT^-GCaMP6f in the GPe of PV-Cre mice. The optical fibers were placed over the cell bodies at the target site, except for in the DAT-Cre mice, when the fiber was implanted over the NAcLat to image terminals of VTA^DA^→NAcLat cells. We found that spontaneous cellular activity, as measured one day following cocaine administration, was depressed in DMS^D2^ cells, elevated in GPe^PV^→ventral midbrain cells, depressed in SNr^GABA^ cells, and elevated in VTA^DA^→NAcLat cells relative to measurements performed on the day prior to cocaine injection (Figure 4C-N). Given that the first three of these cell populations are inhibitory, the increase of activity in GPe^PV^ and VTA^DA^→NAcLat cells, and decrease in SNr^GABA^ cells, are as predicted by our circuit model. Notably, we observed a similar elevation in spontaneous activity of GPe^PV^→ventral midbrain cells as in GPe^PV^ cells generally (Figure 4H, S4A-D), suggesting either targeting method captures essentially the same cell population, and thus we combined results from these populations here and in Figure 5.

**Figure 4:**
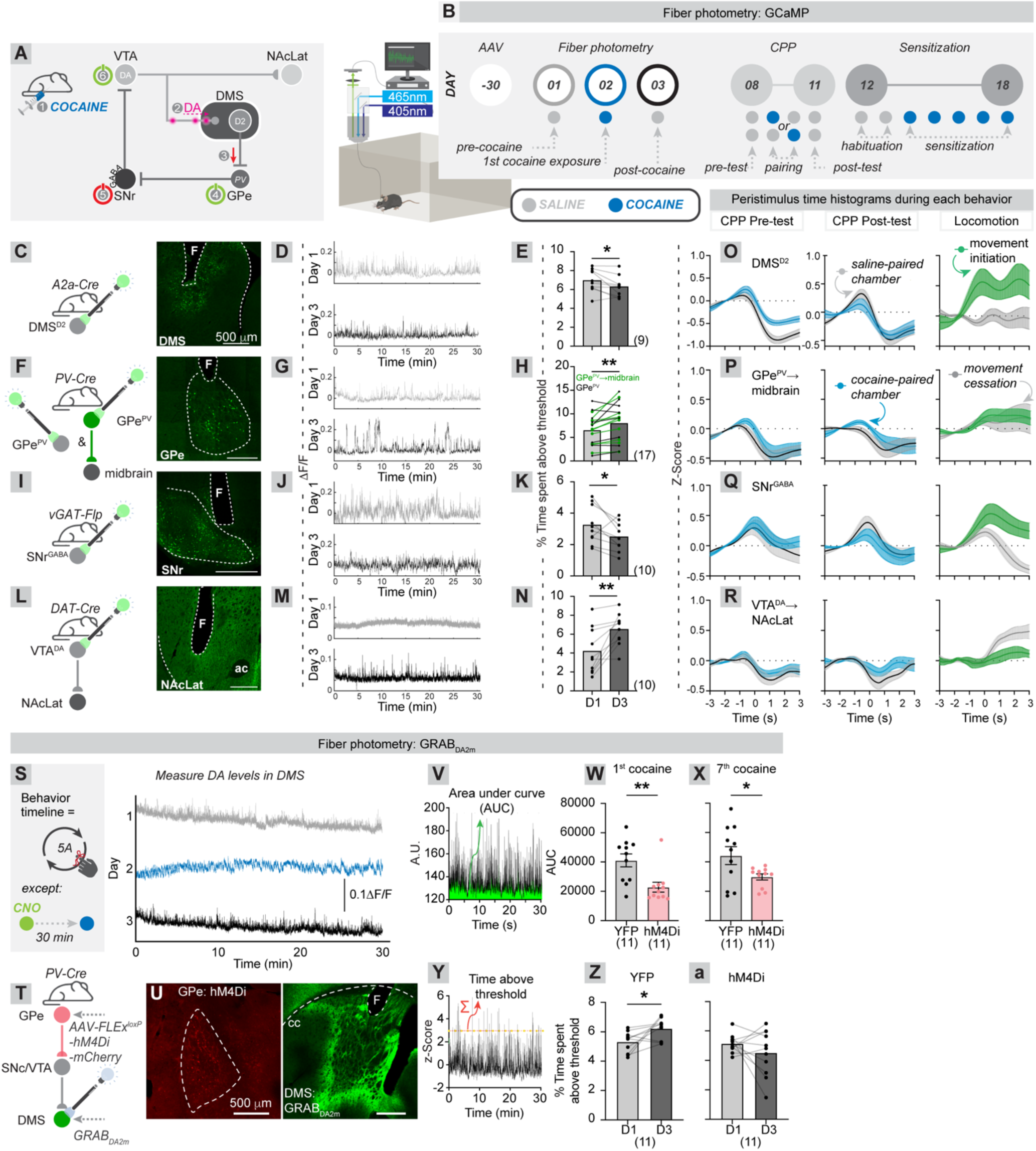
Recording activity of each node of the DMS^D2^→GPe^PV^→SNr^GABA^→VTA^DA^→NAcLat pathway. (A) Proposed pathway of basal ganglia circuit being studied. Cocaine evokes DA release from VTA^DA^→NAcLat collaterals in the DMS. This reduces activity in DMS^D2^ cells that project to GPe^PV^ cells, increasing their activity. This increase then elevates inhibition onto SNr^GABA^ cells, reducing their inhibition. This in turn disinhibits VTA^DA^→NAcLat cells. (B) Schematic of experiment. GCaMP was expressed in each of the four targeted cell populations by injecting AAV-FLEx^loxP^-GCaMP7f into DMS (A2a-Cre), or VTA (DAT-Cre), or AAV-FLEx^FRT^-GCaMP6f into SNr (vGAT-Flp) or GPe^PV^→ventral midbrain cells (PV-Cre, AAV_rg_-DIO-FLPo in the VTA, AAV-FLEx^FRT^- GCaMP6f in the GPe). A chronic fiber was implanted approximately 0.2 mm above where GCaMP was injected, except in the case of in DAT-Cre mice, where the fiber was implanted into NAcLat. Experiments began one month later. GCaMP activity was recorded for 30 minutes in the open field on Day 1 following a saline injection, Day 2 following an injection of 15 mg/kg cocaine, and on day 3 following a saline injection. Five days later, mice were run through CPP and sensitization protocols, as performed previously. (C-N) Schematic, GCaMP injection/fiber implantation sites for each of the four conditions, sample traces from Days 1-3, and percentage of time spent above the activity threshold for each population on Days 1 and 3. Shown are data from DMS^D2^ cells (C-E; p = 0.025, paired t-test), GPe^PV^ cells (combined GPe^PV^→ventral midbrain and GPe^PV^, F-H; p = 0.005, paired t-test), SNr^GABA^ cells (I-K; p = 0.033, paired t-test), and VTA^DA^→NAcLat terminals (L-N; p = 0.004, paired t-test). Individual comparison for GPe^PV^ and GPe^PV^→ventral midbrain cells are shown in Figure S4D. (O-R) Peri-stimulus time histograms (PSTH) for DMS^D2^ cells (O, n = 9), GPe^PV^→ventral midbrain cells (P, n = 9), SNr^GABA^ cells (Q, n = 10), and VTA^DA^→NAcLat terminals (R, n = 10) during chamber crossings in the CPP pre-test and post-test, as well as locomotor initiation and cessation in the open field. Locomotion recordings were taken from the second day of habituation (Day 13, panel B). A comparison of GPe^PV^ and GPe^PV^→ventral midbrain cells is shown in S4N-O. (S) Experimental schematic. AAV-FLEx^loxP^-hM4Di-mCherry, or AAV-DIO-YFP was injected bilaterally into the GPe of PV-Cre mice, AAV-GRAB_DA2m_ was injected unilaterally into the right DMS, and a chronic fiber was implanted above the DMS. Four weeks later, experiments were initiated. The experimental protocol was the same as for GCaMP imaging experiments shown in panel B, except that 5 mg/kg CNO was given i.p. 30 minutes prior to each cocaine exposure for all mice. Sample traces from the DMS are shown for Days 1-3. (T) Schematic of the pathway modulation and recording procedure. (U) Sample images of hM4Di expression in the GPe, and GRAB_DA2m_ expression in the DMS of PV-Cre mice. (V) Sample area under the curve (AUC) analysis of GRAB_DA2m_ signal. (W-X) Significantly reduced DA signaling was observed in mice expressing hM4Di in GPe^PV^ cells than those expressing YFP during the 1^st^ cocaine dose on day 2 of the protocol (W; YFP vs. hM4Di, 41125 vs. 22830, p = 0.004), as well as the final cocaine dose in the sensitization test (X; YFP vs. hM4Di, 44328 vs. 29803, p = 0.036). (Y) Sample measure of spontaneous activity, as performed for GCaMP imaging experiments. (Z) Spontaneous DA signaling one day prior to, and one day following, cocaine administration in mice expressing YFP in GPe^PV^ cells (paired t-test, p = 0.040, n = 11). (a) Spontaneous DA signaling one day prior to, and one day following, cocaine administration in mice expressing hM4Di in GPe^PV^ cells (paired t-test, p = 0.096, n = 11).

**Figure 5:**
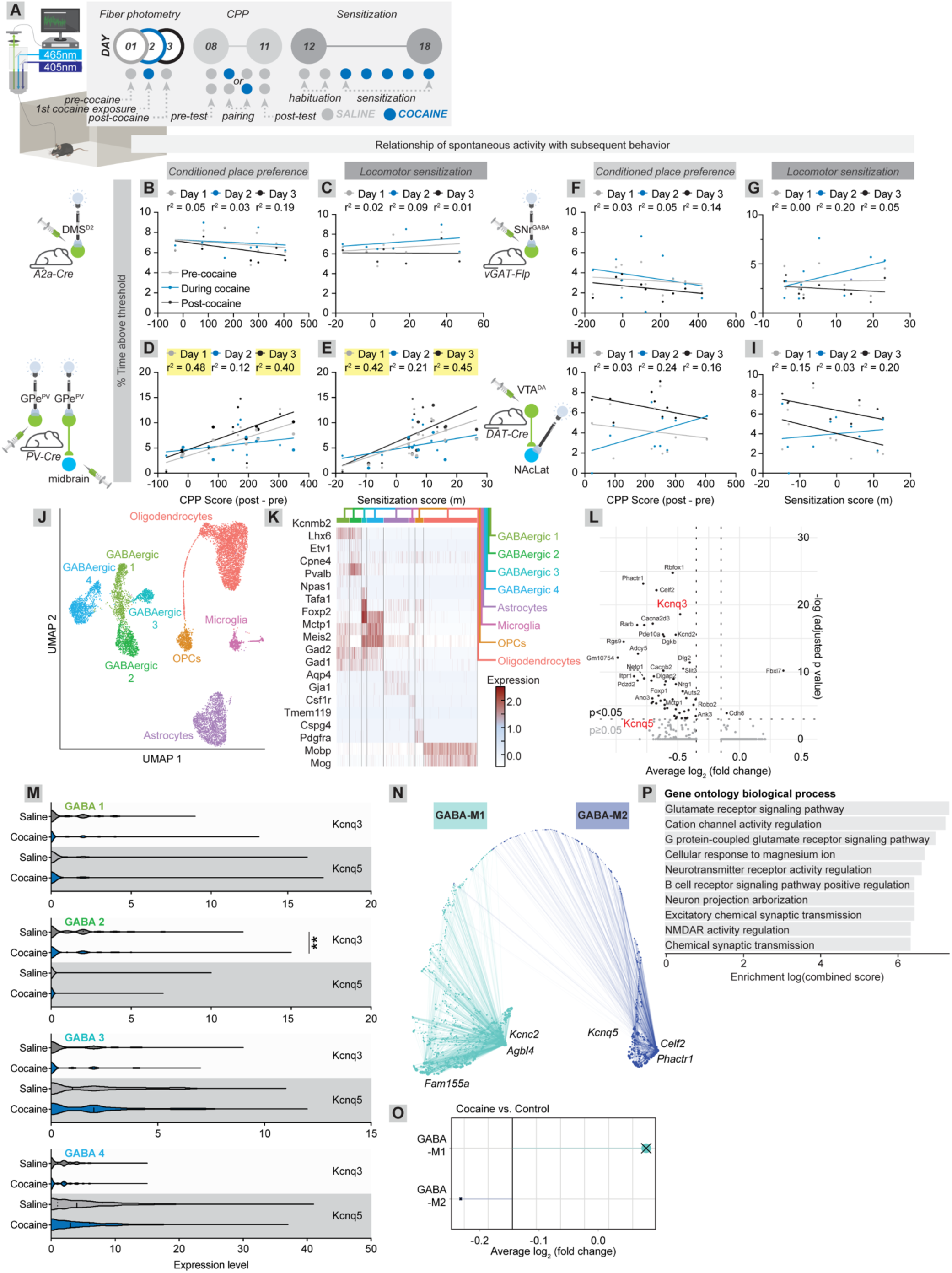
Relationship of GPe^PV^ cell activity relates to subsequent CPP and sensitization, and cocaine-induced downregulation of *Kcnq3* and *Kcnq5* in GPe^PV^ cells. (A) Schematic of fiber photometry experiments. Mice were the same as those used in Figure 4B-N. (B-C) The spontaneous activity of DMS^D2^ cells one day preceding and one day following cocaine administration was not correlated with the extent of CPP formed to a subsequent cocaine dose/chamber pairing (B; D1 r^2^ = 0.05, p = 0.57; D3 r^2^ = 0.19, p = 0.24, n = 8), nor was it correlated with sensitization following repeated cocaine injections (C; D1 r^2^ = 0.02, p = 0.75; D3 r^2^ = 0.01, p = 0.78, n = 8). (D-E) The spontaneous activity of GPe^PV^ cells one day preceding and one day following cocaine administration was highly correlated with subsequent CPP (D; D1 r^2^ = 0.48, p = 0.006; D3 r^2^ = 0.40, p = 0.016, n = 14) and sensitization (E; D1 r^2^ = 0.42, p = 0.004 D3 r^2^ = 0.45, p = 0.002, n = 18). Data from GPe^PV^→ventral midbrain and GPe^PV^ cells generally were combined in panels D and E as all data (e.g., Figure 4, S4, and our previous work^18^) indicated these were largely the same cell populations. Data from GPe^PV^→ventral midbrain cells are shown as large dots, and GPe^PV^ cells as small dots. (F-G) The spontaneous activity of SNr^GABA^ cells one day preceding and one day following cocaine administration was not correlated with CPP (F; D1 r^2^ = 0.03, p = 0.62; D3 r^2^ = 0.14, p = 0.29, n = 9) or sensitization (G; D1 r^2^ = 0.00, p = 0.92; D3 r^2^ = 0.05, p = 0.54, n = 9). (H-I) The spontaneous activity of VTA^DA^→NAcLat cells one day preceding and one day following cocaine administration was not correlated with CPP (H; D1 r^2^ = 0.03, p = 0.64; D3 r^2^ = 0.16, p = 0.25, n = 10) or sensitization (I; D1 r^2^ = 0.15, p = 0.26; D3 r^2^ = 0.20, p = 0.19, n = 10). (J) UMAP plot of cells identified in snRNA-seq experiments from the GPe. Four distinct clusters of GABAergic neurons were identified along with astrocytes, oligodendrocytes, microglia, and oligodendrocyte precursor cells (OPCs). (K) Heatmap of gene expression for known marker genes that define cell types. (L) Volcano plot of differentially expressed genes in cocaine- vs. saline-treated mice. Many more genes were downregulated after cocaine administration than were upregulated. This includes the voltage-gated potassium channel *Kcnq3*. Both *Kcnq3* and *Kcnq5* are shown. (M) Violin plots of *Kcnq3* and *Kcnq5* expression in each GABAergic cluster identified in panel J. Expression of *Kcnq3* was significantly downregulated in GABAergic cluster 2 following cocaine (*log_2_FC = -0.22, p-val adj = 0.0013*); all other comparisons were not statistically significant. (N) UMAP plot of gene modules calculated using hdWGCNA applied to snRNA-seq data from GABAergic cells. Two gene expression modules were identified, the teal (M1) and blue (M2) modules. Both *Kcnq3* and *Kcnq5* were part of the M2 cluster, and *Kcnq5* was identified as a hub gene of this cluster. (O) Average fold change of genes in each expression module when comparing cocaine-treated to saline-treated mice. Expression of genes in module 1 were upregulated while expression of genes in module 2 were downregulated, though neither of these differences were statistically significant. (P) Gene ontology analysis of genes in module 2, focusing on biological processes.

To test hypothesis 2, we plotted the peri-stimulus time histogram (PSTH) for each cell population when the animals were crossing into the saline-paired or cocaine-paired chambers in the CPP pre-test and post-test, or initiated or terminated locomotion in the open field, assessed on the second habituation day for the sensitization task (Day 13), a day on which cocaine was not given. Each cell population had a similar response to cocaine- and saline-paired chamber entries in the CPP task, and a clear response was observed in all but the DA cells for locomotor initiation (Figure 4O-R). Furthermore, the magnitude of activity in each cell population in the CPP and locomotor tasks scaled with the behavioral responses observed, consistent with the observation that these cells causally contribute to both CPP and sensitization (Figure S4E-L). Notably, the PSTH analysis indicated a similarity in the task-linked activity profiles of GPe^PV^→ventral midbrain and GPe^PV^ cells generally (Figure S4M-O), a second indication that either targeting method captures the same cell population. The DMS^D2^ and SNr^GABA^ cells were activated prior to chamber entries while downstream GPe^PV^→ventral midbrain and VTA^DA^→NAcLat cells were inhibited; in general, the DMS^D2^ and SNr^GABA^ cell pairs, and GPe^PV^→ventral midbrain and VTA^DA^→NAcLat cell pairs show closely matching time courses and magnitudes of activation across the examined time window (Figure 4O-R; S5A-F). To test if this was the case, we took data from the time window between -0.5 s and 1.5 s around the event of interest for each of the three tests for each cell population and performed a dimensionality reduction analysis to assess the overall relationship between the activity of each cell population across tests. These tests reinforced our conclusion that DMS^D2^ and SNr^GABA^ cells form one pair with similar activity profiles across tests, and GPe^PV^→ventral midbrain and VTA^DA^→NAcLat cells form another pair (Figure S5G-H). These patterns are consistent with a circuit defined by inhibitory connections, where the 1^st^ and 3^rd^ nodes of the circuit have a common activity pattern, while the 2^nd^ and 4^th^ have paired patterns as well.

To test hypothesis 3 that targeted modulation of activity of each circuit node should impact the others, we measured the effect of inhibiting GPe^PV^ neurons on DA levels in the DMS, which could arise at least in part through collaterals of VTA^DA^→NAcLat cells. According to our circuit model, inhibition of GPe^PV^ cells should prevent cocaine-induced elevations in their own activity through blunting cocaine-evoked DA release in the DMS (Figure 4A). Inhibition of GPe^PV^ cells during cocaine administration should also reduce the elevated levels of spontaneous DA release in the DMS long-term following cocaine administration, given that it should prevent the elevation in activity of VTA^DA^→NAcLat cells. To test this hypothesis, we injected AAV-FLEx^loxP^-hM4Di or AAV-DIO-YFP as a control bilaterally into the GPe, the DA sensor GRAB_DA2m_ was injected unilaterally into the DMS of PV-Cre mice, and a chronic fiber was implanted over the DMS (Figure 4S-U). Two weeks later when experiments were initiated, we administered 5 mg/kg CNO i.p. 30 minutes prior to each cocaine administration, using the same behavioral protocol as used for VTA^DA^ cell inhibition in Figure 1. We found that CNO administration into mice expressing hM4Di, but not YFP, blunted the cocaine-induced elevation of DA levels in the DMS, measured either during the very first cocaine exposure, or the last (Figure 4V-X). Furthermore, the long-lasting elevation in spontaneous DA release at baseline following cocaine administration was also reduced in hM4Di-expressing mice (Figure 4Y-a), as was cocaine-induced changes in activity in the CPP task during saline-paired and cocaine-paired chamber crossings (Figure S5P-Q). These results suggest that GPe^PV^ cell inhibition during cocaine administration prevents cocaine-induced behavioral changes by blunting acute DA increases in DMS driven by cocaine (Figure 4V-X). Notably, the pattern of DA activity in the DMS (Figure S5J-O) is similar to the dynamic pattern of GCaMP-based activity for VTA^DA^→NAcLat cells, assessed in the NAcLat (Figure 4R), providing further evidence that these results reflect activity defined in two different ways from the same cells.

### Levels of spontaneous activity in GPe^PV^ cells relate to the development of cocaine CPP and sensitization arising after subsequent cocaine exposures

Our data support the existence of a DMS^D2^→GPe^PV^→SNr^GABA^→VTA^DA^→NAcLat closed-loop circuit that controls the development of cocaine CPP and sensitization. However, it is not clear how inter-animal variations in this circuit may relate to individual variability in response to drugs of abuse. A central question in substance abuse is why only some individuals who take drugs – approximately 20% for cocaine^25^ – develop a substance use disorder, whereas others do not. One hypothesis is that differential function of the DMS^D2^→GPe^PV^→SNr^GABA^→VTA^DA^→NAcLat circuit may impact how animals respond to cocaine and influence the later development of addiction-related behaviors. If this is true, identifying how activity throughout the circuit tracks with subsequent development of drug reward and sensitization could provide predictive biomarkers of future responses to cocaine, as well as allow for identification of key brain nodes that control individual susceptibility to substance abuse-related behaviors. To test this hypothesis, in the same GCaMP-expressing animals as used in Figure 4, we assessed how spontaneous activity in each node one day before cocaine administration, during cocaine administration, and one day following cocaine administration related to the development of cocaine CPP and sensitization (Figure 5A). When assessing DMS^D2^, SNr^GABA^, and VTA^DA^→NAcLat cell activity, we did not find a strong relationship between activity in these nodes at any of the three timepoints tested and the subsequent development of CPP or sensitization (Figure 5B-C, F-I). However, we noted a strong relationship between activity in GPe^PV^ cells and the development of both CPP and sensitization in both the days before and after cocaine administration (Figure 5D-E). These results indicate that activity of these cells, even before cocaine administration, is predictive of behavioral outcomes in response to subsequent cocaine exposures.

### Single nucleus RNA sequencing reveals cocaine-induced changes in ion channel expression in GPe^PV^ <u>cells</u>

The *in vivo* elevation of GPe^PV^ cell spontaneous activity could be caused by changes in inputs, intrinsic changes such as in GPe^PV^ cell gene expression, or both. We showed that inhibitory inputs onto GPe^PV^ cells were reduced following a single cocaine exposure (Figure 3), suggesting that a change in input control may contribute to the elevation of *in vivo* spontaneous activity. Meanwhile, the changes in intrinsic excitability we previously observed^18^ are most likely due to changes in ion channel expression within GPe^PV^ cells themselves. To assess potential alterations in ion channel expression, we performed single nucleus RNA sequencing (snRNA-seq) on nuclei from the GPe of mice following a single saline or cocaine exposure. After filtering out nuclei based on QC metrics (Figure S6A), 5,485 nuclei from the saline-treated group and 4,718 nuclei from the cocaine-treated group were retained, for a total of 10,203 nuclei. Following data integration and dimensionality reduction, we applied Uniform Manifold Approximation and Projection, UMAP^26^, to visualize nuclei and confirmed that each group, saline-treated or cocaine-treated, had a similar cellular composition (Figure S6B). Annotation of cell types was then performed manually using known marker genes. Based on these annotations, we identified microglia, astrocytes, oligodendrocytes, oligodendrocyte precursor cells (OPCs), and 4 distinct clusters of GABAergic neurons (Figure 5J). The neuronal composition in our dataset aligns with previous characterization of neurons in the GPe as almost completely GABAergic, with subpopulations expressing a combination of markers such as *Pvalb* and *Lhx6*, or *Foxp2* and *Meis2* (Figure 5K)^27^. Differential gene expression analysis was then performed for each major cell type to explore changes induced by a single cocaine exposure. While there were no significantly differentially expressed genes in any of the glial cell clusters (p-adj < 0.05, and log_2_FC > 0.1 or log_2_FC < - 0.1), there were 2 significantly upregulated genes and 53 significantly downregulated genes in GABAergic nuclei in saline-treated versus cocaine-treated mice. Of particular interest were differentially expressed ion channel genes that may influence cellular excitability. We found that *Kcnq3* and *Kcnq5*, encoding voltage gated potassium channel pore-forming α subunits KCNQ3 and KCNQ5 that form homomeric or heteromeric channels that generate M-type potassium currents in the brain^28^, were both downregulated after cocaine exposure (Figure 5L; Supplementary Table 1). There are five *Kcnq* genes in mice (and humans), but in the GABAergic clusters, only *Kcnq3* was significantly downregulated following cocaine exposure (log_2_FC = -0.230, p-adj = 8.35x10^-09^). *Kcnq5* also appeared to be downregulated in GABAergic neurons, though this effect did not reach statistical significance (log_2_FC = -0.346, p-adj = 0.335). Notably, when considering all nuclei within the dataset, *Kcnq5* was significantly downregulated after cocaine exposure (log_2_FC = -0.484, p-adj = 7.33x10^-07^), consistent with a mild reduction of *Kcnq5* expression in the GPe generally. When examining GABAergic clusters individually, *Kcnq3* was downregulated in all clusters following cocaine, though this effect was only statistically significant in GABAergic cluster 2, which expressed the gene *Pvalb* (Figure 5K, M). Furthermore, the other *Kcnq* genes — *Kcnq1*, *Kcnq2*, and *Kcnq4* — were expressed at much lower levels than *Kcnq3* and *Kcnq5* in GABAergic neurons (Figure S6C) and were not significantly differentially expressed in any cell type. These findings suggest that KCNQ3/5 homomers and/or heteromers are present in GPe^PV^ neurons and are depleted after a single cocaine exposure.

### Identification of gene co-expression networks in GPe GABAergic neurons

To go beyond consideration of individual genes and further probe for differences in broad patterns of gene expression in GABAergic neurons, we utilized high-dimensional weighted gene co-expression network analysis, hdWGCNA^29^. WGCNA builds a correlation network of genes based on their expression across samples. This correlation network can then be split into network modules or sets of genes that are highly interconnected^30^. hdWGCNA adapts WGCNA for high dimensional RNA-seq data such as snRNA-seq and can be used to identify cell-type specific modules differentially regulated in disease^31^. We applied hdWGCNA to the GABAergic neuron clusters in our snRNA-seq data. Following network construction and hierarchical clustering, we found 2 gene modules in GABAergic neurons: M1 (teal) containing 911 genes, and M2 (blue) containing 504 genes (Figure 5N, Supplementary Table 2). Notably, *Kcnq3* and *Kcnq5* were both in the M2 module, with *Kcnq5* being one of the top 3 hub genes, or genes most central to the module network. These results further support the hypothesis that changes in *Kcnq3* and *Kcnq5* are among the most significant gene expression changes induced by cocaine and may play a role in cocaine-induced activity changes in GPe cells. The module eigengenes (MEs) for M1 and M2 were calculated to summarize gene expression for each module. Differential module eigengene (DME) analysis indicated that M2 was downregulated in cocaine-treated mice, while M1 was upregulated, though neither module was statistically significantly different when considering all 4 GABAergic subclusters combined (Figure 5O).

We then used DME analysis to assess how these two modules changed or not in each GABAergic subcluster in saline- vs. cocaine-treated mice. We found that the blue M2 module was the most significantly downregulated in cocaine-treated mice in GABAergic clusters 1 and 2, and slightly downregulated in cluster 4. There was no significant difference in the M2 ME for GABAergic cluster 3. The teal M1 module was not significantly different in any of the 4 GABAergic subclusters (Figure S6F). Notably, the 2 GABAergic subclusters with large and significant changes in module M2 (GABAergic clusters 1 and 2) express parvalbumin, while the other two GABAergic subclusters (GABAergic clusters 3 and 4) do not, indicating that the most significant cocaine-induced gene expression changes occur largely in the GPe^PV^ cells (Figure 5K, S6D). Lastly, we explored the biological functions of the genes comprising module M2. Gene ontology (GO) analysis indicated that M2 is highly associated with biological processes regulating glutamate receptor signaling and cation channel activity (Figure 5P), indicating that the most significant gene expression changes in GPe^PV^ cells following a single cocaine exposure likely relate to the control of cellular activity following cocaine.

### Discovery of a novel KCNQ3/5 activator that blunts cocaine reward, sensitization, and self-administration

Our data indicate that GPe^PV^ cells play a critical role in gating early drug-induced behavioral adaptations. However, without a clear way to selectively modulate the activity of GPe^PV^ cells in a non-invasive manner, the therapeutic potential of this approach is limited. A previous study noted that the GPe to SNr pathway has the densest expression of the combination of KCNQ3 and KCNQ5 protein in the mouse brain^32^. Notably, while we also found that *Kcnq3* and *Kcnq5* transcripts were expressed at relatively high levels in GPe cells, *Kcnq2*, the other member of the *Kcnq* family expressed in the brain, is expressed at lower levels in GPe cells (Figure S6C). This raises the intriguing possibility that GPe cells may preferentially express KCNQ3/5 and not KCNQ2/3 heteromeric channels, the latter of which are thought to be responsible for the majority of the M-current in the nervous system as a whole. Thus, we may be able to effectively inhibit the GPe→SNr pathway relatively selectively if we could specifically open KCNQ3/5 homomeric and/or heteromeric channels without opening KCNQ2/3 heteromeric channels. Several existing drugs can open KCNQ channels broadly, such as retigabine^33^, or KCNQ2/3 selectively, such as the retigabine derivative SF0034^34^. KCNQ2/3 heteromers are best known for their link to brain diseases associated with cellular hyperexcitability, including epilepsy and tinnitus^35–38^. As low expression or activity of KCNQ2 and KCNQ3 can contribute to these diseases, drugs that open KCNQ2/3 channels have anti-epileptic effects^33,34^. Given the enhanced expression of KCNQ3/5 in the GPe→SNr pathway, we hypothesized that selectively opening KCNQ3/5 channels would prevent early cellular and circuit changes associated with cocaine use. Until recently, no known compounds could selectively open KCNQ3/5 heteromeric channels. Fortunately, we recently found that carnosic acid, derived from rosemary, acts as a relatively selective KCNQ3/5 opener^39^. To test its selectivity, we conducted voltage clamp experiments in *Xenopus laevis* oocytes in which we expressed either KCNQ2 or a pore mutant of KCNQ3 that enables KCNQ3 homomers to pass current, as homomeric wild-type KCNQ3 cannot (Figure 6A). We found that 10 μM carnosic acid had negligible effects on KCNQ2 channels but very strongly opened KCNQ3 channels (Figure 6B-E). When exploring its effects on functionally relevant heteromeric channels, we found that carnosic acid had no significant effect on KCNQ2/3 channels, but strongly opened KCNQ3/5 channels (Figure S7A-Y). Therefore, carnosic acid has the ideal selectivity profile for inhibiting the GPe→SNr pathway.

**Figure 6:**
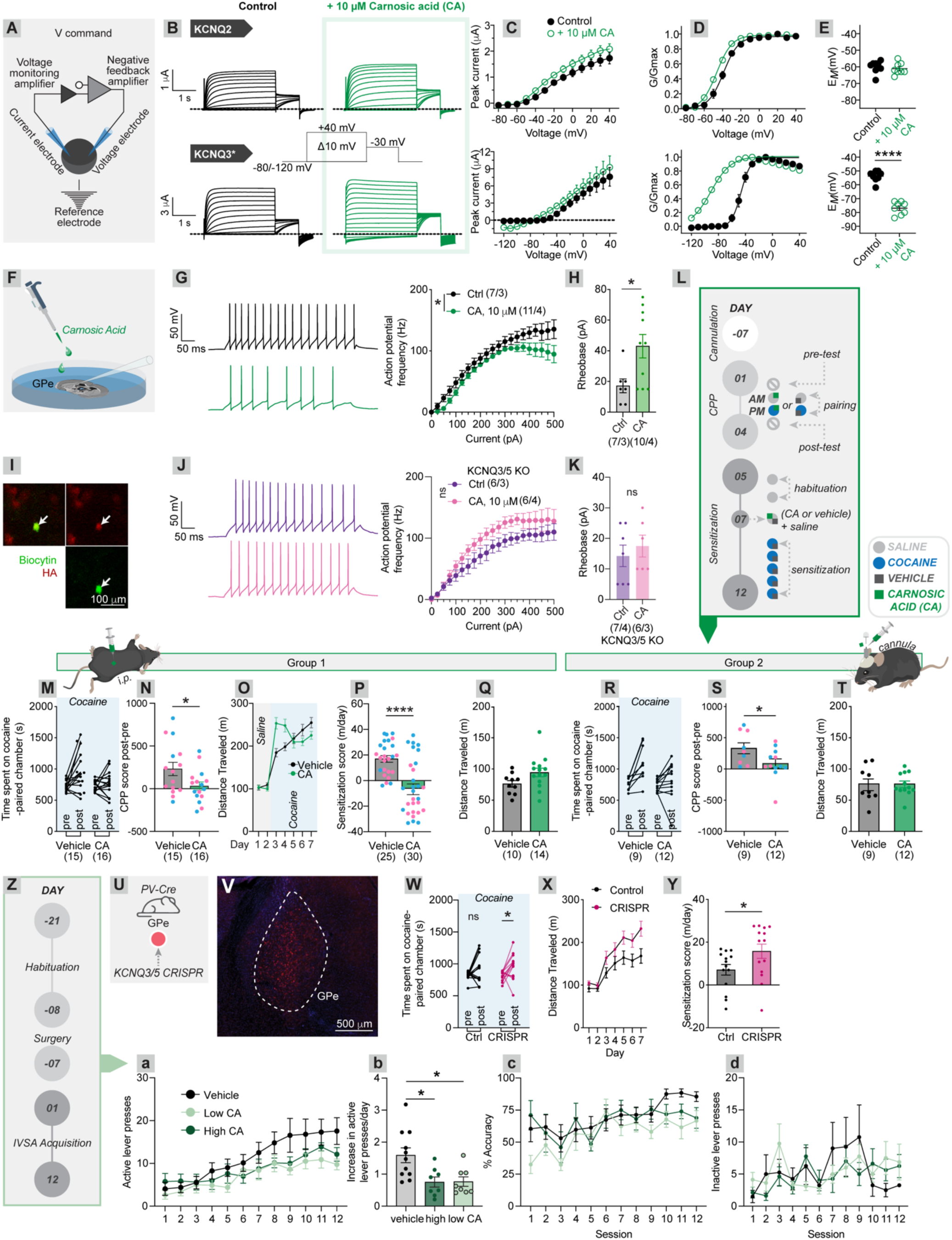
The KCNQ3/5 opener carnosic acid reduces GPe^PV^ cell excitability, impairs cocaine reward and sensitization, and reduces cocaine self-administration. (A) Schematic of dual voltage clamp setup in Xenopus oocytes to test carnosic acid effects on select members of the *Kcnq* gene family. (B) Mean traces showing KCNQ currents as indicated, +/- carnosic acid (voltage protocol inset), expressed in *Xenopus* oocytes and measured by two-electrode voltage clamp; *n* = 8 per group for all panels. KCNQ3* = KCNQ3-A315T, a pore mutant that facilitates robust current expression from homomeric KCNQ3. (C) Peak current for KCNQ2 and KCNQ3*. (D) Conductance (G/Gmax) for KCNQ2 and KCNQ3*. (E) Membrane voltage for each condition with and without carnosic acid application. KCNQ2, p-value = 0.55; KCNQ3*, p < 0.0001. (F) Schematic of experiments to test the effects of carnosic acid on GPe^PV^ cell excitability. (G) Carnosic acid caused GPe^PV^ cells to fire fewer action potentials across a range of current injections (Two-way repeat measures ANOVA condition effect, vehicle vs. carnosic acid p = 0.033). n = 7 cells for vehicle, 10 for carnosic acid. Shown are sample traces from each condition following a 50 pA current injection. (H) The rheobase, defined as the minimum current required to elicit a single action potential, was higher in carnosic acid-treated cells (17.1 pA vs. 43.0 pA, p = 0.020). (I) Sample image of a recorded, biocytin-filled cell co-staining with HA, which was tagged to Cas9. (J) Carnosic acid application to slices where *Kcnq3* and *Kcnq5* were deleted from GPe cells using CRISPR had no significant effect on GPe^PV^ cell excitability, and in fact, slightly increased it (Two-way repeat measures ANOVA condition effect, vehicle vs. carnosic acid p = 0.12). n = 7 cells for vehicle, 10 for carnosic acid. Shown are sample traces from each condition following a 50 pA current injection. (K) Carnosic acid application did not alter the rheobase of cells lacking *Kcnq3* and *Kcnq5* (14.3 pA vs. 17.5 pA, p = 0.54). n = 6 cells each. (L) Schematic for behavioral experiments where the behavioral effects of carnosic acid were tested. These included both i.p. administration of 2.5 mg/kg carnosic acid, as well as direct infusion into the GPe of 10 μM carnosic acid. (M-N) i.p. administration of 2.5 mg/kg carnosic acid 30 minutes prior to cocaine administration blocked cocaine CPP. Saline vs. carnosic acid, 231.5 s vs. 31.5 s, p = 0.034. Males (blue) and females (magenta) are denoted by the colors of the individual data points. (O-P) i.p. administration of 2.5 mg/kg carnosic acid 30 minutes prior to cocaine administration blocked cocaine-induced locomotor sensitization. Saline vs. carnosic acid, 18.0 m vs. -6.4 m, p < 0.0001. One point for the carnosic acid group is located below the axis. (Q) i.p. administration of 2.5 mg/kg carnosic acid had no significant effect on locomotion as assessed in a 30-minute locomotor test with carnosic acid only (no cocaine). Saline vs. carnosic acid, 76.3 m vs. 94.5 m, p = 0.06. (R-S) Local GPe infusion of 10 μM carnosic acid prior to cocaine administration blocked cocaine CPP. Saline vs. carnosic acid, 328.6 s vs. 86.5 s, p = 0.045. (T) Local GPe infusion of 10 μM carnosic acid prior to cocaine administration had no significant effect on locomotion as assessed in a 30-minute locomotor test with carnosic acid only (no cocaine). Saline vs. carnosic acid, 75.5 m vs. 75.4 m, p = 0.99. (U) Schematic of experiments testing the effects CRISPR-based knockout of *Kcnq3* and *Kcnq5* on cocaine-induced behaviors. (V) Verification of Cas9 expression in the GPe, as assessed by immunostaining for HA. (W) Control animals receiving carnosic acid did not form a significant CPP, whereas animals in which *Kcnq3* and *5* were knocked out in GPe cells did form a significant CPP (control pretest 817.9 s, posttest 897.1 s, p = 0.23; CRISPR pretest 820.2 s, 952.9 s posttest, p = 0.044, n = 13 and 14, respectively). (X-Y) Animals in which *Kcnq3* and *5* were knocked out in GPe cells showed an elevated locomotor response to cocaine, as well as enhanced sensitization relative to control mice (709.4 s vs. 1,575 s, p = 0.046, n = 14 for each). (Z) Timeline of cocaine IVSA experiments. (a) Number of active lever presses over the twelve-day acquisition period for animals treated with vehicle, or one of two carnosic acid concentrations, termed low (830 μg/kg) and high (7.5 mg/kg). Carnosic-acid treated mice administered significantly less cocaine over this period (Two-way ANOVA, group effect p < 0.0001; vehicle vs. low CA p < 0.0001; vehicle vs. high CA p = 0.015). n = 11 for vehicle group, 8 for CA-treated groups. (b) Carnosic acid-treated mice showed a significantly lower escalation in cocaine intake over the course of the twelve-day self-administration session (One-way ANOVA p = 0.0059; vehicle vs. low CA 1.59 vs. 0.77, p = 0.012; vehicle vs. high CA 1.59 vs. 0.76, p = 0.011). (c) Low concentrations of carnosic acid significantly reduced the preference for pressing the active lever, while higher concentrations of carnosic acid had no effect (Two-way ANOVA group effect p = 0.0004; vehicle vs. low CA p = 0.0002; vehicle vs. high CA p = 0.44). (d) No significant effects of carnosic acid on inactive lever pressing were observed (Two-way ANOVA group effect p = 0.70).

Our snRNA-seq data indicated that both *Kcnq3* and *Kcnq5* were downregulated in GPe cells (Figure 5L-M). By this same logic, we hypothesized that opening KCNQ3/5 channels during cocaine administration should have a similar effect as hM4Di-mediated inhibition of these cells, which prevented cocaine CPP and sensitization (Figure S1C-F, and previous work^18^). To test this, we first measured whether carnosic acid application reduced the intrinsic neuronal excitability of GPe^PV^ cells. We injected AAV-FLEx^loxP^-GFP into the GPe of PV-Cre mice, and two weeks later, cut acute slices and performed whole-cell patch clamp recordings in current clamp mode, measuring the number of action potentials fired across a range of current injections, and the rheobase, defined as the lowest current injection needed to elicit action potentials (Figure 6F-H). Carnosic acid-treated cells exhibited a higher rheobase and fired fewer action potentials than vehicle-treated cells, two key indicators of a reduced excitability (Figure 6G-H). To test whether this effect was mediated by KCNQ3/5 channels, we used a CRISPR-based gene knockout strategy, designing guide RNAs to *Kcnq3* and *Kcnq5* along with a Cre-dependent GFP and virally delivering them along with Cas9 to the GPe of PV-Cre mice (Figure S7Z-a). Mice in which Cas9 but not guide RNAs were delivered were used as controls. We again tested whether carnosic acid reduced the excitability of GPe^PV^ cells, this time in GPe^PV^ cells lacking KCNQ3/5. Carnosic acid application had no significant effect on the action potential rate nor the rheobase of cells in which *Kcnq3* and *Kcnq5* were knocked out, indicating that carnosic acid’s effects in GPe^PV^ cells are indeed mediated by KCNQ3/5 (Figure 6I-K).

We next tested whether carnosic acid could reduce cocaine-mediated reward and sensitization. To test this, we first injected 2.5 mg/kg carnosic acid i.p. 30 minutes prior to cocaine injection and tested the effects on cocaine CPP and sensitization (Figure 6L). Systemic administration of carnosic acid completely prevented cocaine CPP and sensitization, with no effects on locomotion (Figure 6M-Q). To test if these effects were mediated by the GPe, we locally infused 10 μM carnosic acid into the GPe 30 minutes prior to cocaine injection in the CPP test, and again found that cocaine CPP was blocked, again with no effects on locomotion (Figure 6R-T). Importantly, similar results for both systemic and local infusions were observed in males and females, suggesting that carnosic acid works approximately equally well in both sexes (Figure 6M-T). To test whether this *in vivo* effect was mediated by KCNQ3/5, we used CRISPR to again delete the *Kcnq3* and *Kcnq5* genes the GPe, and tested cocaine reward and sensitization when animals were given 2.5 mg/kg carnosic acid 30 minutes prior to cocaine (Figure 6U-V). In control mice infected only with Cas9-expressing AAVs, carnosic acid impaired cocaine CPP, whereas GPe-specific CRISPR-mediated *Kcnq3* and *Kcnq5* knockout mice developed significant CPP even following carnosic acid injection (Figure 6W). Similarly, knockout mice showed an elevated locomotion in response to cocaine as well as a higher level of sensitization than carnosic acid-treated mice (Figure 6X-Y). These results support the conclusion that carnosic acid administration *in vivo* can reduce cocaine reward and sensitization through reducing the excitability of GPe^PV^ cells via increased activity of KCNQ3/5.

Lastly, we tested whether carnosic acid could impair volitional drug-taking in an intravenous self-administration (IVSA) model (Figure 6Z). Mice were implanted with indwelling jugular vein catheters, and after seven days of recovery, mice passing catheter patency testing were placed in operant chambers and trained to self-administer cocaine under a fixed ratio 1 (FR1) schedule using a previously detailed protocol^40^. One-hour acquisition trials were conducted daily for ten days. On the first day, mice were primed with a cocaine infusion (0.5 mg/kg/infusion). Subsequently, the pressing of the active (cocaine-paired) lever actuated a cue light and a micropump-driven cocaine infusion (0.5 mg/kg/infusion) followed by a 40-second timeout period, on an FR1 schedule. We quantified the number of active and inactive lever presses excluding the timeout period, assessed the accuracy of active lever presses, as well as the rate of escalation of cocaine intake over the twelve-day session. Two different concentrations of carnosic acid were tested (low: 830 μg/kg and high: 7.5 mg/kg) to assess whether i.p. delivered carnosic acid may impact cocaine intake across a range of concentrations. We observed that mice treated with either the low or the high carnosic acid concentrations showed a reduced number of active lever presses over the twelve-day session, and a reduced rate of escalation of cocaine consumption (Figure 6a-b). This effect is unlikely to be due to an inability to perform or learn the task. Although mice receiving the low concentration of carnosic acid showed a slightly lower rate of accuracy of active lever presses over the twelve-day session, they reached the same level as vehicle-treated mice by the end of the trial; furthermore, mice treated with the high concentration of carnosic acid showed a nearly identical accuracy for lever pressing as vehicle-treated controls (Figure 6c). In addition, the number of inactive lever presses was not significantly different among the three groups (Figure 6d). The most parsimonious explanation for these results is that carnosic acid administration reduced cocaine intake in a cocaine IVSA task, likely through reducing the rewarding value of cocaine infusions via reduced excitability of GPe^PV^ cells.

## DISCUSSION

Here we comprehensively map the key brain circuits that mediate cocaine reward and sensitization, implicate the GPe as the central mediator of these effects, and demonstrate that dampening GPe^PV^ cell activity through activation of KCNQ3/5 can reduce volitional cocaine consumption. We first link the behavioral functions of the GPe specifically to VTA^DA^ subpopulations projecting to the NAcLat and show that these regions are linked through a disinhibitory mechanism involving the SNr. We next show that VTA^DA^→NAcLat cells send collaterals to the DMS, and that DA release from these DMS collaterals is critical for triggering elevations in activity of GPe^PV^ cells. This effect occurs in part through a reduction in inhibitory drive from DMS^D2^ cells onto GPe^PV^ cells that is likely triggered by cocaine’s actions within the DMS. Within this closed-loop circuit, all the cell populations appear to carry similar signals during the CPP reward task as well as in locomotion, and the magnitude of activity in each population scales with the extent to which cocaine elicited CPP and sensitization. Interestingly, activity in the GPe^PV^ cells, but not the three other cell populations in the circuit, before and after a single dose of cocaine tracked with subsequent development of CPP and sensitization, indicating that 1) GPe^PV^ cell activity differs between animals, and 2) this variation is linked to the extent to which animals develop CPP and sensitization to cocaine doses that are administered later on. We then performed snRNA-seq on GPe cells and found that these cells express lower levels of the voltage-gated potassium ion channel genes *Kcnq3* and *Kcnq5* following cocaine administration, which would be predicted to contribute to an elevation in spontaneous activity. Given that the GPe→SNr pathway expresses high levels of KCNQ3/5 channels, this pathway is a uniquely attractive target for carnosic acid, which we recently identified opens KCNQ3/5 but not KCNQ2/3 channels^39^. Indeed, carnosic acid administration blocked cocaine reward and sensitization via KCNQ3/5 in GPe^PV^ cells, likely by reducing the intrinsic excitability of these cells, and impaired volitional administration of cocaine, making carnosic acid a promising potential therapeutic against psychostimulant abuse.

### Updated circuit framework for cocaine reward and sensitization

This study reinforces our prior work indicating that the GPe acts as a gateway for drug reward and sensitization^18^. Here we extend these observations in several important ways. First, we comprehensively map the entire circuit in which the GPe is embedded and provide a clear mechanistic framework by which cocaine induces long-lasting pathological changes in this circuit (Figure 7). While we originally identified the GPe as being an altered *input* to the ventral midbrain whose elevated activity disinhibited VTA^DA^ cells, this left unanswered the question of how this change occurs. As cocaine acts in large part by blocking the dopamine transporter (DAT) and leads to transient increases in extracellular DA levels, the elevation in GPe^PV^ cell activity is likely triggered either directly or indirectly through DA’s actions in the brain. Here we demonstrate that VTA^DA^→NAcLat cells, through collaterals to the DMS, likely indirectly trigger an elevation in GPe^PV^ cell activity. GPe^PV^ cells then control the activity of VTA^DA^→NAcLat cells; this provides a mechanism by which the dorsal striatum can feed back onto the VTA. Therefore, DA levels in the NAc are likely affected by changes in GPe^PV^ cell activity, albeit in NAcLat, a structure that has historically not been included in studies of the NAc, which have typically included the NAcCore and NAc shell (referred to here as the NAcMed). While the majority of work in this area has centered on the roles of the VTA and NAc in drug reward and sensitization^8,41–45^, our work indicates that DA action in the DMS critically contributes, at least in part through its projection to the GPe. Notably, activation of the GPe^PV^→ ventral midbrain projection directly is neither rewarding nor locomotor activating (Figure S1S-Y), consistent with its role as a gate for cocaine-induced plasticity. Therefore, our data do not conflict with a large body of literature implicating the NAc in mediating the reinforcing effects of stimulants such as cocaine^4^ but rather, extends these observations to include additional brain regions and cell types not previously thought to be central to cocaine-induced behavioral changes. As a note, retrobead-mediated labeling of input cells, used in the experiments to show GPe^PV^-mediated disinhibition of VTA^DA^→NAcLat cells (Figure 1a-c), has the potential to cause cellular toxicity. While we attempted to limit toxicity by diluting the beads prior to injection and all recordings of bead-containing cells were controlled by recording the latency to first action potential with and without optical stimulation in the same cells, this is an important potential caveat to consider.

**Figure 7:**
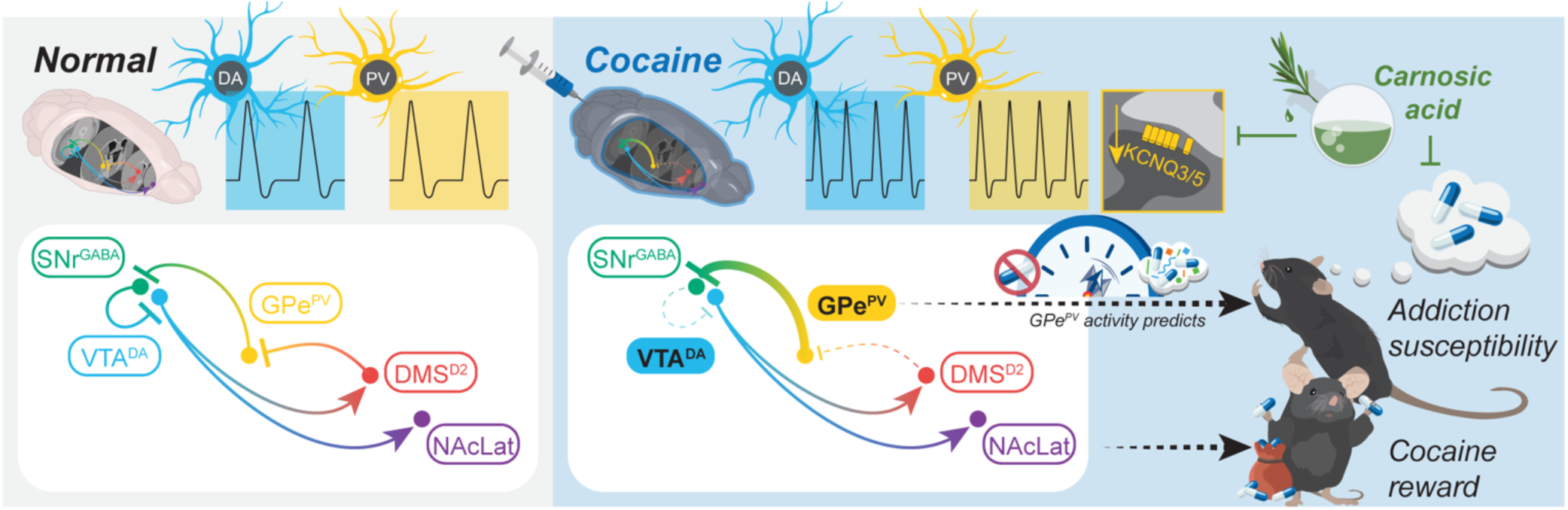
Schematic of key findings from this study. We implicated a four-node closed loop circuit consisting of DMS^D2^→GPe^PV^→SNr^GABA^→VTA^DA^→NAcLat (DMS) cells in cocaine-induced behavioral plasticity. The DMS, GPe, and SNr connections are all GABAergic, and control DAergic output of VTA^DA^→NAcLat cells, which mediate cocaine reward. Following cocaine, spontaneous activity in the GPe^PV^ and VTA^DA^→NAcLat (DMS) cells increases, leading to elevations in DA signaling in the striatum. This elevation in GPe^PV^ cell activity is mediated both by decreases in inhibitory drive from DMS^D2^ cells as well as a downregulation in the expression of *Kcnq3* and *Kcnq5* in GPe^PV^ cells. As baseline GPe^PV^ cell activity is linked to behavioral responses to subsequent cocaine administrations, GPe^PV^ cell activity can be viewed as a barometer of the likelihood of developing addiction-related behaviors later in life in response to cocaine. Carnosic acid, a KCNQ3/5 channel opener, can prevent cocaine reward when administered systemically or within the GPe, and can interfere with cocaine self-administration behaviors. The hypothesized mechanism of action is through inhibition of GPe^PV^ cells, which prevents cocaine-induced behavioral plasticity.

Interestingly, while DA depletion from the NAc impairs cocaine self-administration behavior, lesioning either the NAcCore or NAcMed does not^46,47^. Notably, the strategy to deplete DA using 6-OHDA administration in the above study spares most DA cells located medially within the VTA, which includes mostly VTA^DA^→NAcMed cells (and VTA^DA^→mPFC cells), and preferentially kills those projecting to the dorsal striatum and in the lateral VTA, which mostly include VTA^DA^→NAcLat cells^48^. Thus, while DA in the NAcCore and NAcMed likely influence cocaine reward, a hypothesis supported by our own results (Figure 1E-F), the DMS/NAcLat is likely the critical site for cocaine reward. It is also important to consider that the NAcMed is heterogeneous, with the dorsal NAcMed being implicated in reward and ventral NAcMed in aversion^49–51^. Our studies do not distinguish these subregions, providing an explanation for why broad chemogenetic inhibition of VTA^DA^→NAcMed cells in this study prevented reward despite evidence that at least subsets of these cells may not encode reward^51,52^. The assertion of the importance of the NAcLat in cocaine reward is consistent with recent studies showing that DA terminals in NAcMed are excited by aversive outcomes but DA terminals in other NAc subregions are depressed, while excitation to reward-related cues occur in VTA^DA^→NAcLat but not VTA^DA^→NAcMed cells^51^. Another recent study confirmed these observations and extended them by showing that activation patterns in the NAc subregions recapitulate those of their VTA^DA^ inputs^52^. Furthermore, stimulation of NAcLat terminals in the VTA is rewarding, while stimulation of NAcMed terminals induces a state of behavioral suppression but not reward^53^. Activation of NAcLat terminals in the ventral midbrain leads to disinhibition of VTA^DA^→NAcLat cells, while activation of the NAcMed inhibits VTA^DA^→NAcMed cells^53^, thus providing a circuit mechanism by which the NAcLat may play a central role in cocaine reward.

### GPe^PV^ cell activity as a barometer for future cocaine-induced behaviors

While we implicated each of the nodes studied here (DMS^D2^→GPe^PV^→SNr^GABA^→VTA^DA^→NAcLat) in cocaine reward and sensitization, activity in the GPe is the most closely linked to inter-animal variation in cocaine reward and sensitization (Figure 5). Importantly, this link can be observed even before cocaine is administered, indicating that the basal level of GPe^PV^ cell activity in cocaine-naïve mice can be linked to subsequent responses to cocaine. These data indicate that GPe^PV^ cell activity may be used, even in cocaine-naïve mice, as a “barometer” of how animals will respond to cocaine in the future. Our report linking the native, un-altered *in vivo* activity of a defined cell population using fiber photometry to the magnitude of cocaine-induced place preference and sensitization in response to subsequent cocaine injections is a significant advance in understanding the biological sources of behavioral vulnerability. As these experiments were performed in genetically identical mice, individual variations in cellular activity between mice are presumably driven by epigenetic differences. Therefore, differences in cellular activity could be one mechanism by which epigenetic mechanisms may influence behavioral differences induced by cocaine or other drugs^15,54–57^. This information could be crucial towards identifying vulnerable populations before developing substance use disorders or designing targeted interventions to prevent the development of addiction. The question is, then: what factors control this inter-animal variability in the behavioral response to cocaine? One interesting hypothesis is that genes linked to susceptibility may be differentially expressed in GPe^PV^ cells, and the basal levels or dynamic regulation of these genes by drugs of abuse may relate to individual susceptibility to substance abuse. The principal method for identifying genes and gene variants that may contribute to addiction susceptibility has been through Genome-Wide Association Studies (GWAS). Interestingly, a number of GWAS hits^58^ and genes previously found to be differentially expressed in transcriptomic studies of cocaine^59,60^ are also significantly differentially expressed in our snRNA-seq data of the GPe, such as *Cacnb2*, *Nrg1*, *Rasgef1b*, and *Rbfox1* (Supplementary Table 1). These results suggest that the expression levels of these genes in the GPe may play an important role in influencing susceptibility to developing cocaine abuse by differentially regulating cellular activity of GPe cells critical for early drug-induced behavioral adaptations, a topic for future study.

Here, we found that basal levels of GPe^PV^ cell activity positively correlated with the extent of cocaine CPP and sensitization. This is consistent with our hypothesis that GPe^PV^ cell activity promotes cocaine-induced plasticity: according to this hypothesis, higher basal GPe^PV^ cell activity would enhance the long-lasting behavioral effects of cocaine. We observed evidence for two separate mechanisms by which cocaine may enhance the spontaneous activity of GPe^PV^ cells, including a reduction in inhibitory drive as well as a downregulation of voltage-gated potassium ion channels that would hyperpolarize these cells. It is not clear which mechanism, or combination of the two, may be responsible for inter-animal variability in susceptibility. Notably, it appears that GPe^PV^ cells downregulated genes related to glutamatergic synaptic function (Figure 5O-P), potentially as a compensatory mechanism to reduce excitatory input drive. However, we also note that no significant changes in excitatory transmission were observed one day following cocaine administration (Figure 3O-Q). As GPe^PV^ cells undergo both a reduction in inhibitory input and elevation in intrinsic excitability, which both would serve to increase persistent cellular activity above baseline, these changes likely are a pathological consequence of cocaine administration rather than an adaptive compensatory response. Therefore, we hypothesized that counteracting the elevation in excitability should effectively combat cocaine-induced behavioral changes. Indeed, effectively countering this hyperactivity through opening KCNQ3/5 channels via administration of carnosic acid, including directly in the GPe, was sufficient to blunt cocaine’s behavioral effects. Furthermore, we found that carnosic acid also reduced cocaine intake in the IVSA model, indicating that systemic activation of KCNQ3/5 channels is sufficient to prevent key reward-related behavioral adaptations that contribute to substance abuse.

### Carnosic acid and KCNQ modulators as potential addiction therapeutics

Despite the magnitude of the national addiction epidemic, there remains an absence of effective therapeutics to treat psychostimulant addiction. This contrasts with opioid abuse, which has garnered significantly more national attention, and for which numerous effective pharmacological interventions exist, including methadone, naltrexone, and suboxone. Unfortunately, these therapeutics are mainly for opioid use disorder and are not effective for psychostimulant use. The effectiveness of GPe^PV^ cell inhibition in preventing drug reward^18^ combined with the high KCNQ3/5 expression in the GPe→SNr pathway^32^ led us to pursue the use of carnosic acid, shown by our recent publication to be the first known KCNQ3/5 opener that activates neither KCNQ2 nor KCNQ2/3^39^, as an intervention to prevent psychostimulant intake. We demonstrate that carnosic acid injection prior to cocaine administration can prevent cocaine reward and sensitization and can reduce volitional cocaine intake (Figure 6L-d). Furthermore, carnosic acid has been reported to have wide-ranging health benefits, including exhibiting anti-inflammatory, anti-viral, anti-obesity, anti-carcinogenic, and anti-depressive properties, and generally shows promise as a neuroprotective agent, including against Alzheimer’s and Parkinson disease^61–69^. However, to our knowledge, this is the first report of its potential as an anti-addictive agent. As such, we should note that much remains unknown about carnosic acid’s effects on the brain, both acutely and long-term. Both KCNQ3 and KCNQ5 are expressed widely in the brain, and though they are highly expressed in the GPe, expression is not specific to the GPe. Thus, while our data directly support our hypothesis that carnosic acid exerts its effects through KCNQ3/5 channels in GPe^PV^ cells (Figure 6F-K, U-Y), it is possible that carnosic acid works at least in part through other mechanisms. Notably, however, areas of KCNQ3/KCNQ5 expression overlap are not particularly common, with the GPe being an exception. KCNQ3 is thought to most commonly heteromerize with KCNQ2 in the brain, and recently KCNQ5 was also found to heteromerize with KCNQ2 in mouse brain; tripartite complexes of KCNQ2, KCNQ3, and KCNQ5 were also detected^70^. While not perfectly selective for KCNQ3/5 heteromers, carnosic acid is qualitatively more effective at opening this versus other KCNQ heteromers (Figure S7A-Y) and this likely imparts a regional specificity to the effects of carnosic acid that is unique among neuronal KCNQ channel openers. Furthermore, carnosic acid is inactive against the epithelial and cardiac-expressed KCNQ1^39^. Lastly, while carnosic acid has been used in other studies, generally without substantial side effects noted at the concentrations used here, more work needs to be done to assess its effects on a wide variety of cognitive and motivational processes that may mitigate its potential as an anti-addiction therapeutic.

---

**Figure S1:**
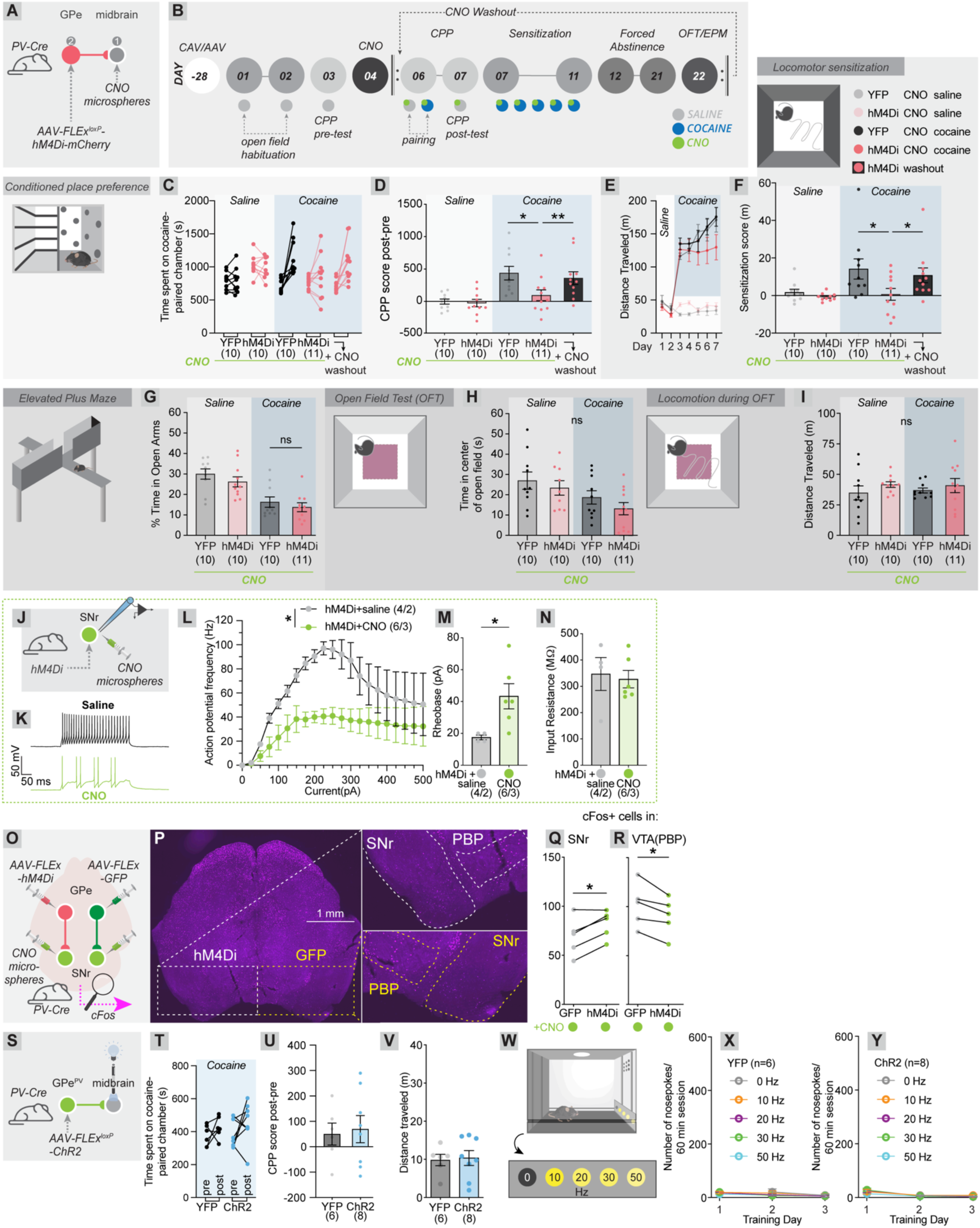
Effects of GPe^PV^→ventral midbrain cell inhibition and activation, and validation of CNO microspheres. (A) Schematic of experiment. AAV-FLEx^loxP^-hM4Di-mCherry, or AAV-DIO-YFP as a control was injected bilaterally into the GPe, and CNO-releasing microspheres were injected into the ventral midbrain to inhibit outputs of GPe^PV^ cells in the ventral midbrain. (B) Abridged timeline for testing cocaine place preference and locomotor sensitization after CNO microspheres were injected, as performed previously^14^. This protocol was used so that all cocaine administrations would occur within 7 days of microsphere injection, the timeframe in which the microspheres are predicted to release CNO. (C-D) Cocaine pairing, but not saline pairing, introduced a place preference, which was prevented by chemogenetic inhibition of GPe^PV^→ventral midbrain cells (One-way ANOVA p = 0.0004; pairwise t-tests with multiple comparisons corrections, YFP/saline vs. hM4Di/saline, -1.49 s vs. -25.32 s, p = 1.0; YFP/saline vs. YFP/cocaine, -1.49 s vs. 342.2 s, p = 0.0014; YFP/saline vs. hM4Di/cocaine, -1.49 s vs. 94.6 s, p = 0.80; hM4Di/saline vs. YFP/cocaine, -25.32 s vs. 342.2 s, p = 0.0007; hM4Di/saline vs. hM4Di/cocaine, -25.32 s vs. 94.6 s, p = 0.67; YFP/cocaine vs. hM4Di/cocaine, 342.2 s vs. 94.6 s, p = 0.013). After performing the elevated plus maze and open field tests on these mice, we then re-ran the CPP and sensitization tests, during a period at which the microspheres should no longer be releasing CNO. These same mice that did not previously form a CPP did so during the washout period. Paired t-test, p = 0.0023. (E-F) Repeated cocaine, but not saline injections, caused locomotor sensitization, which was prevented by chemogenetic inhibition of GPe^PV^→ventral midbrain cells (One-way ANOVA p = 0.0090; pairwise t-tests with multiple comparisons corrections, YFP/saline vs. hM4Di/saline, 1.7 m vs. -0.77 m, p = 0.95; YFP/saline vs. YFP/cocaine, 1.7 m vs. 14.2 m, p = 0.050; YFP/saline vs. hM4Di/cocaine, 1.7 m vs. 0.63 m, p = 1.0; hM4Di/saline vs. YFP/cocaine, -0.77 m vs. 14.2 m, p = 0.014; hM4Di/saline vs. hM4Di/cocaine, -0.77 m vs. 0.63 m, p = 0.99; YFP/cocaine vs. hM4Di/cocaine, 14.2 m vs. 0.63 m, p = 0.024). After performing the elevated plus maze and open field tests on these mice, we then re-ran the CPP and sensitization tests, during a period at which the microspheres should no longer be releasing CNO. These same mice that did not previously sensitize to cocaine did so during the washout period (paired t-test, p = 0.047). (G) Inhibition of GPe^PV^→ventral midbrain cells had no significant effect on time spent in the open arms of the elevated plus maze (YFP/saline vs. YFP/cocaine, 30.0% vs. 16.3%, p = 0.002; YFP/cocaine vs. hM4Di/cocaine, 16.3% vs. 13.76%, p = 0.48). (H) Inhibition of GPe^PV^→ventral midbrain cells had no significant effect on the time spent in the middle of an open field (YFP/saline vs. YFP/cocaine, 27.0 s vs. 18.7 s, p = 0.37; YFP/cocaine vs. hM4Di/cocaine, 18.7 s vs. 13.2 s, p = 0.67). (I) Inhibition of GPe^PV^→ventral midbrain cells had no significant effect on basal locomotion, as assessed in the five-minute open field test. (One-way ANOVA p = 0.67). Notably, cocaine was not given to any group acutely during this test. (J) Schematic of electrophysiology experiments to test whether the slow-release CNO microspheres used in this study inhibit hM4Di-expressing cells. For these tests, hM4Di was expressed in SNr cells, and CNO microspheres were injected in the SNr. Saline injections in place of CNO microspheres were used as controls. (K) Sample excitability traces from saline-treated and CNO microsphere-treated conditions, following a 100 pA current injection. (L) hM4Di-expressing cells fire fewer action potentials over a range of current injections in the presence of CNO microspheres than in saline-treated controls (Two-way ANOVA, group effect p = 0.017). n = 4 saline, 6 CNO microspheres. (M) hM4Di-expressing cells showed a higher rheobase in the presence of CNO microspheres than in saline-treated controls (17.5 pA vs. 43.3 pA, p = 0.032). (N) No significant differences in membrane input resistance were observed between groups (346.9 mOhm vs. 327.5 mOhm, p = 0.77). (O) Schematic for experiments assessing whether CNO microsphere-mediated inhibition of GPe^PV^ cell terminals in the ventral midbrain impact cFos expression in the ventral midbrain. GFP was expressed in GPe^PV^ cells in one hemisphere, hM4Di-mCherry was expressed in the other, and one month later, CNO microspheres were injected into the SNr in both hemispheres. Four days later, cFos expression was assessed one hour following a saline i.p. injection. (P) Sample midbrain image showing cFos expression. Images on the R are zoom-ins of the ventral midbrain, with the appropriate treatment (GFP or hM4Di expression in the GPe) indicated. The parabrachial pigmented nucleus (PBP) of the VTA, and SNr, are denoted. (Q) Higher numbers of cFos+ neurons were observed in the SNr in hemispheres where hM4Di was expressed in GPe^PV^ cells (paired t-tests, p = 0.020). (R) Lower numbers of cFos+ neurons were observed in the PBP in hemispheres where hM4Di was expressed in GPe^PV^ cells (paired t-tests, p = 0.019). (S) Experimental schematic of optogenetic activation of GPe^PV^→ventral midbrain projections. Mice expressing YFP rather than ChR2-YFP were used as controls. (T-U) Optogenetic activation of GPe^PV^ cells during a 30-minute pairing session had no effect on CPP (YFP vs. ChR2, 50.0 s vs. 69.3 s, p = 0.79). (V) Optogenetic activation of GPe^PV^ cells had no effect on locomotion over 5 minutes in the open field (YFP vs. ChR2, 9.8 m vs. 10.4 m, p = 0.84). (W) Schematic of intracranial self-stimulation task. Nosepoking into each port had a different outcome. Two seconds of a 0 Hz, 10 Hz, 20 Hz, 30 Hz, or 50 Hz stimulus was given following a successful nosepoke. (X) YFP-expressing mice showed no significant nosepoking activity over a 3-day trial. (Y) ChR2-expressing mice showed no significant nosepoking activity over a 3-day trial.

**Figure S2:**
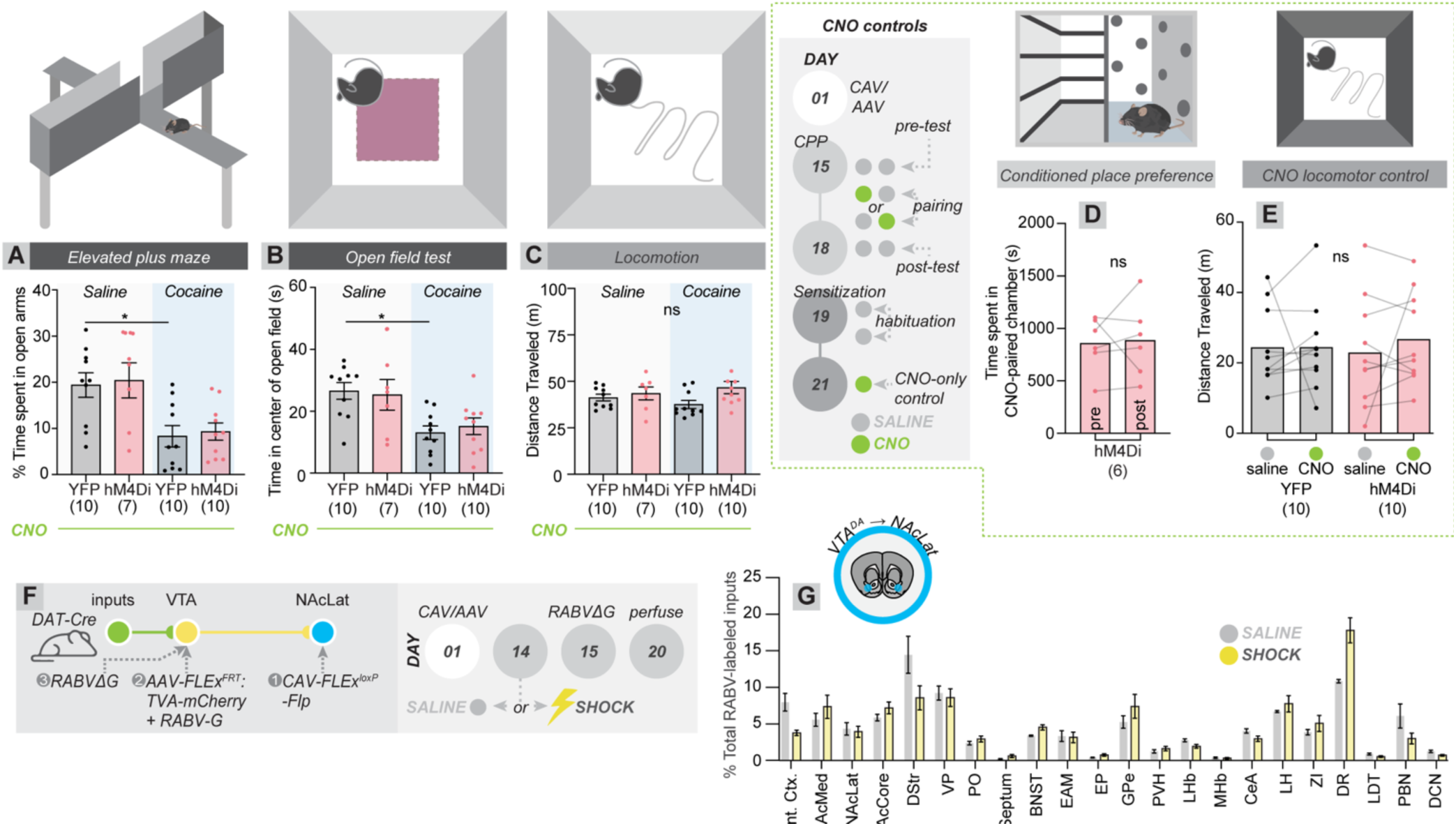
Tests for anxiety-related behavior and controls for chemogenetic modulation of VTA^DA^→NAcLat cells, and cTRIO from VTA^DA^→NAcLat cells following an auditory fear conditioning protocol. (A) Cocaine-treated mice tested on the EPM show reduced time spent in the open arms of the EPM relative to saline-treated controls, and hM4Di-mediated inhibition of VTA^DA^→NAcLat cells had no effect (YFP/saline vs. YFP/cocaine, 19.4% vs. 8.3%, p = 0.017; YFP/cocaine vs. hM4Di/cocaine, 8.3% vs. 9.3%, p = 0.99). (B) Cocaine-treated mice tested in the OFT show reduced time spent in the center of the open field relative to saline-treated controls, and hM4Di-mediated inhibition of VTA^DA^→NAcLat cells had no effect (YFP/saline vs. YFP/cocaine, 26.6 s vs. 13.0 s, p = 0.0024; YFP/cocaine vs. hM4Di/cocaine, 13.0 s vs. 15.2 s, p = 0.55). (C) All treatment groups traveled the same distance in the open field when tested during the forced abstinence period, when cocaine was not acutely given. One-way ANOVA p = 0.11. (D) In animals expressing hM4Di in VTA^DA^→NAcLat cells, a CNO pairing to one chamber of the CPP apparatus (no cocaine) caused neither a place preference nor place aversion (paired t-test, p = 0.83). (E) 5 mg/kg CNO given i.p. had no effect on locomotion in mice expressing YFP or hM4Di in VTA^DA^→NAcLat cells as assessed in a five-minute locomotor test (paired t-tests, YFP p = 0.99; hM4Di p = 0.40). (F) Schematic of cTRIO experiments from VTA^DA^→NAcLat cells in mice performing an auditory fear conditioning task. (G) No overall differences were identified between shocked and unshocked groups (Two-way ANOVA, group effect p >0.99, interaction p < 0.0001), and the level of labeling in the GPe was not significantly different between groups (% labeled inputs in the GPe, 5.2% vs. 6.1%, p = 0.60, n = 4 each).

**Figure S3:**
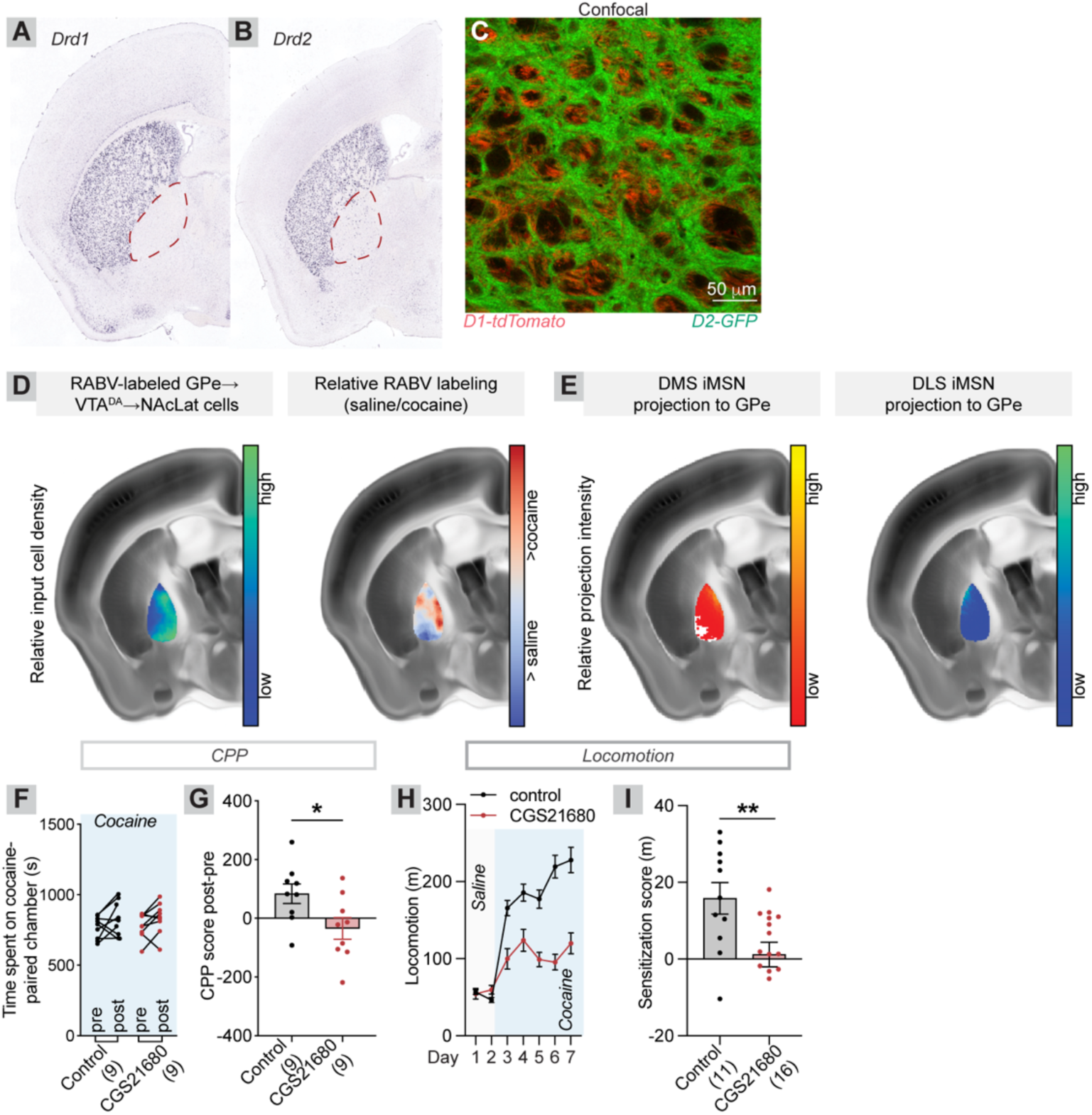
Expression of dopamine receptors in the GPe, anatomical topology of DMS^D2^→GPe→VTA^DA^→NAcLat connections, and effects of pharmacological modulation of A2a receptors. (A) In situ hybridization against *Drd1*. The GPe is circled. Images for panels H and I are borrowed from the Allen Mouse Brain Atlas^71^. (B) In situ hybridization against *Drd2*. (C) Confocal image of the GPe in D1-tdTomato/D2-GFP double transgenic mice. The GPe is heavily innervated by fibers, particularly from D2 cells in the dorsal striatum. (D) Projection topology of GPe→VTA^DA^→NAcLat cells, as defined by the localization of RABV-labeled cells from cTRIO from VTA^DA^→NAcLat cells in saline- and cocaine-treated mice. (E) Projection topology of dorsal striatum D2 neuron projections to GPe, from the DMS and DLS. The DMS has biased projections to the medial GPe, while the DLS has biased projections to the lateral GPe. (F-G) Cocaine CPP in mice treated with saline or 0.25 mg/kg CGS21680 30 minutes before cocaine administration. CGS21680 prevented cocaine CPP (saline vs. CGS21680, 83.7 s vs. -34.1 s, p = 0.030). (H-I) Locomotor sensitization in mice treated with saline or 0.25 mg/kg CGS21680 30 minutes before cocaine administration. CGS21680 prevented locomotor sensitization (saline vs. CGS21680, 15.8 m vs. 1.2 m, p = 0.0084).

**Figure S4:** Additional data for fiber photometry from GPe^PV^ cells and linking photometry signals to behavior. (A) Schematic of experiments. Mice were recorded one day before, during, and after cocaine administration. (B) Histological image of GCaMP expression in GPe^PV^ cells. (C) Sample fiber photometry traces from GPe^PV^ cells. (D) Percent time spent above the activity threshold in the open field 1 day before and 1 day following a single cocaine administration for both GPe^PV^ and GPe^PV^→ventral midbrain cells. GPe^PV^ cells, p = 0.14, n = 8; GPe^PV^→ventral midbrain cells, p = 0.003, n = 9; paired t-tests. Data from these populations are combined in Figure 4H. (E-H) Relationship of maximum or minimum z-scores for different cell populations to the extent of CPP expressed in each mouse. DMS^D2^, r^2^ = 0.55, p = 0.022; GPe^PV^→ventral midbrain, r^2^ = 0.53, p = 0.027; SNr^GABA^, r^2^ = 0.37, p = 0.063; VTA^DA^→NAcLat, r^2^ = 0.44, p = 0.035. DMS^D2^ n = 9, GPe^PV^→ventral midbrain n = 9, SNr^GABA^ n = 10, VTA^DA^→NAcLat n = 10. (I-L) Relationship of maximum or minimum z-scores for different cell populations to the extent of sensitization expressed in each mouse. DMS^D2^, r^2^ = 0.38, p = 0.078; GPe^PV^→ventral midbrain, r^2^ = 0.33, p = 0.11; SNr^GABA^, r^2^ = 0.29, p = 0.11; VTA^DA^→NAcLat, r^2^ = 0.32, p = 0.087. DMS^D2^ n = 9, GPe^PV^→ventral midbrain n = 9, SNr^GABA^ n = 10, VTA^DA^→NAcLat n = 10. (M) Sample fiber photometry traces from GPe^PV^→ventral midbrain cells from Days 1-3. (N-O) PSTH curves from GPe^PV^ (N; n = 8) cells relative to GPe^PV^→ventral midbrain (O; n = 9) cells. PSTH curves are very similar across behaviors, indicating both methods likely targeted the same cell population.

**Figure S5:**
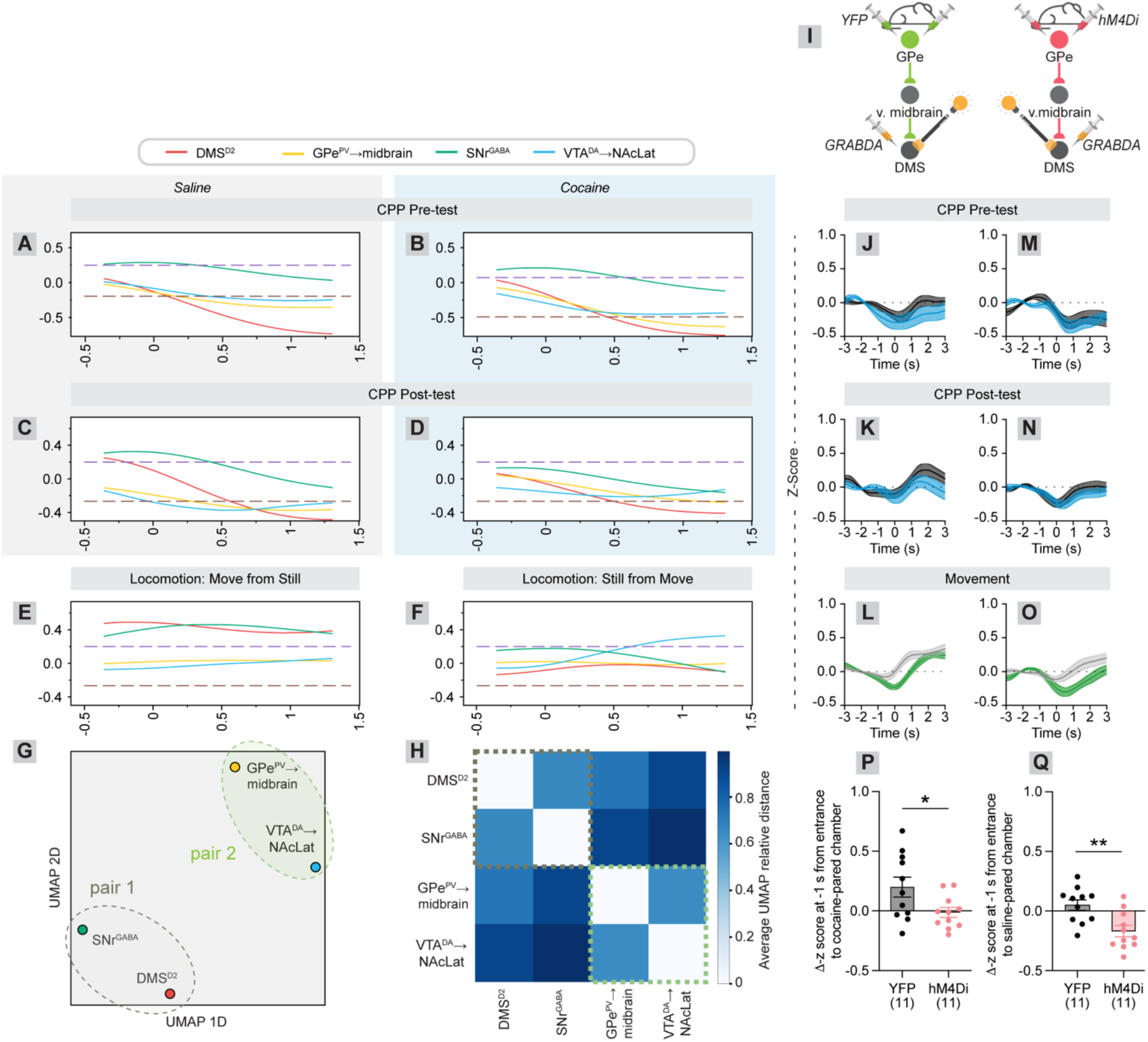
Additional fiber photometry data using GCaMP showing coordinated activity in defined cell pairs, and DA signaling using GRAB_DA2m_ in the DMS. (A-F) Alignment of traces for each cell population during CPP pretest (A-B), posttest (C-D), and locomotion (E-F). (G) To quantitatively determine which conditions had the most similar PSTH curves across the CPP pretest, CPP posttest, and locomotion assessment (open field), the PSTH curves for each test were stitched into one table by condition and UMAP was used to visualize them in two dimensions. Analysis of PSTH curves with UMAP was restricted to -0.5 to 1.5 s surrounding each event. (H) Since individual UMAP embeddings are stochastic depending on initial seeding conditions, we examined the average relative Euclidian distance between the same data points in 20 UMAP embeddings of the same data and plotted the averaged results as a correlogram. (I) Schematics of experiments where either YFP or hM4Di was expressed bilaterally in GPe^PV^ cells, and GRAB_DA2m_ was expressed unilaterally in the DMS (as in Figure 4S-a). (J-O) PSTH curves for mice expressing YFP (J-L) or hM4Di (M-O) in GPe^PV^ cells, and GRAB_DA2m_ in the DMS. (P) Change in z-score from CPP pre-test to CPP post-test at -1 s from entrances into the cocaine-paired chamber, indicating a change in the anticipatory DA signal of chamber crossings. YFP vs. hM4Di, 0.20 vs. -0.01, p = 0.03. (Q) Change in z-score from CPP pre-test to CPP post-test at -1 s from entrances into the saline-paired chamber, indicating a change in the anticipatory DA signal of chamber crossings. YFP vs. hM4Di, 0.05 vs. -0.17, p = 0.003. Panels T and U together indicate that GPe^PV^ cell inhibition during cocaine administration altered how cocaine induces changes DA signaling in the DMS during chamber crossings in the CPP task.

**Figure S6:**
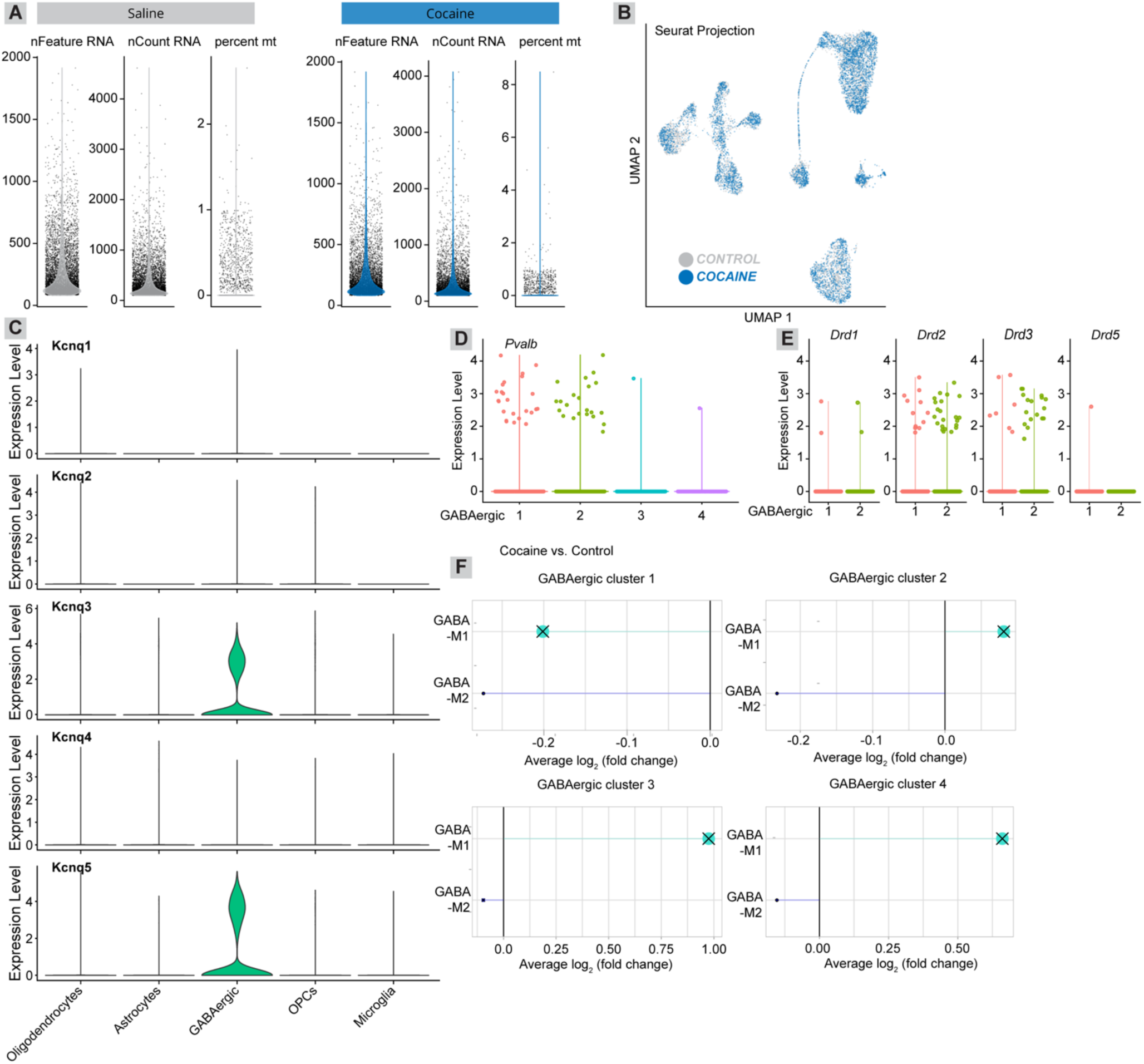
QC metrics, gene expression, and hdWGCNA plots for GPe snRNA-seq data. (A) QC metrics from left to right for saline and cocaine-treated samples: violin plots showing distributions of the number of unique genes detected (nFeature RNA), the number of molecules sequenced (nCount RNA), and the percent of reads that correspond to mitochondrial (mt) genes in each nucleus after QC. (B) UMAP visualization of snRNA-seq data colored by treatment: cocaine or saline. (C) Violin plots showing the normalized expression level of *Kcnq* genes, split by cell type. (D) Violin plot showing normalized expression level of *Pvalb*, split by GABAergic neuron subtype. (E) Violin plot showing normalized expression level of dopamine receptors *Drd1-5*, shown for both clusters of GABAergic clusters that express *Pvalb*. (F) Differential module eigengene (DME) results for each GABAergic gene module detected by hdWGCNA in each GABAergic subcluster. DME analysis was done to compare cocaine-treated and saline-treated samples (positive fold change indicates higher module expression in cocaine). If differences were not statistically significant, this is denoted with an x symbol.

**Figure S7:**
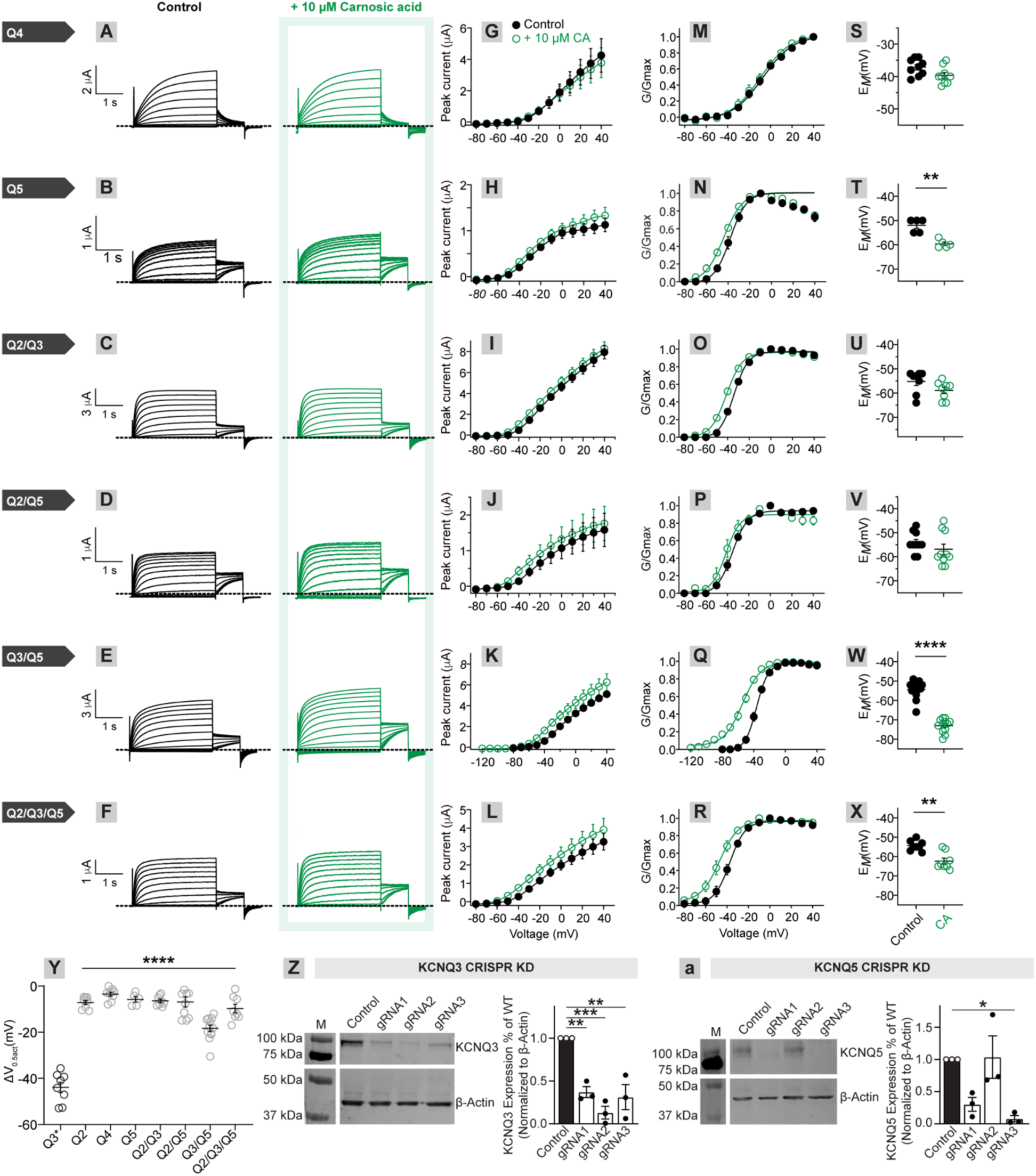
Effects of carnosic acid on various KCNQ isoforms. (A-F) Mean traces showing KCNQ currents as indicated, +/- carnosic acid (voltage protocol inset), expressed in Xenopus oocytes and measured by two-electrode voltage clamp; *n* = 5-12 per group for all panels. (G-L) Mean normalized tail current for traces as in A-F. (M-R) Conductance (G/Gmax) for each subunit combination. (S-X) Mean unclamped resting membrane potential (*E*_M_) for cells as in A-F. (Y) Voltages for the half-maximum activation (Δ1V_0.5act_) for each channel/combination tested. One-way ANOVA, p < 0.0001. (Z-a) Western blot analysis of CRISPR-based knockout of *Kcnq3* and *Kcnq5*. Three unique gRNAs were assessed for each gene. Controls are neurons infected with an AAV expressing Cas9 without a gRNA. AAV preparations were prepared for each gRNA, and these were used to infect primary hippocampal preparations. Western blots were then carried out to measure protein expression of KCNQ3 (Z) and KCNQ5 (a). A significant reduction in protein expression was observed for each gRNA to *Kcnq3* (One-way ANOVA p = 0.0006; pairwise t-tests with multiple comparisons corrections, control vs. gRNA1 1.0 vs. 0.37, p = 0.0045; control vs. gRNA2 1.0 vs. 0.13, p = 0.0005; control vs. gRNA 1.0 vs. 0.31, p = 0.0026). For KCNQ5, a statistically significant reduction in protein expression was only observed for a single gRNA (One-way ANOVA p = 0.0096; pairwise t-tests with multiple comparisons corrections, control vs. gRNA1 1.0 vs. 0.30, p = 0.087; control vs. gRNA2 1.0 vs. 1.04, p > 0.99; control vs. gRNA3 1.0 vs. 0.079, p = 0.024). Based on these results, gRNA2 for *Kcnq3* and gRNA3 for *Kcnq5* were used for CRISPR experiments in Figure 6.

## Acknowledgements

We would like to thank Allison White and Ben Menarchek for assistance with cocaine self-administration experiments. This work utilized the infrastructure for high-performance and high-throughput computing, research data storage and analysis, and scientific software tool integration built, operated, and updated by the Research Cyberinfrastructure Center (RCIC) at the University of California, Irvine (UCI). The RCIC provides cluster-based systems, application software, and scalable storage to directly support the UCI research community. https://rcic.uci.edu. This work was made possible, in part, through access to the following: the Genomics Research and Technology Hub (formerly Genomics High-Throughput Facility) Shared Resource of the Cancer Center Support Grant (P30CA-062203), the Single Cell Analysis Core shared resource of Complexity, Cooperation and Community in Cancer (U54CA217378), the Genomics-Bioinformatics Core of the Skin Biology Resource Based Center @ UCI (P30AR075047) at the University of California, Irvine and NIH shared instrumentation grants 1S10RR025496-01, 1S10OD010794-01, and 1S10OD021718-01. This work was funded by NIH DP2-AG067666, R00-DA041445, R01-DA054374, R01-DA056599, R01-NS130044, TRDRP T31KT1437 and T31IP1426, One Mind OM-5596678, Alzheimer’s Association AARG-NTF-20-685694, New Vision Research CCAD2020-002, Brightfocus A2022031S, Brain and Behavior Research Foundation (NARSAD 26845), and ADPA APDA-5589562 to KTB, T31DT1729 and F30-DA056215 to MH, T32 GM136624 to KB and PD, and NSF GRFP DGE-1839285 to PD.

## STAR+METHODS

Detailed methods include the following:

- KEY RESOURCES TABLE
- RESOURCE AVAILABILITY

- Lead contact
- Materials availability
- Data and code availability
- EXPERIMENTAL MODEL AND SUBJECT DETAILS

- Animals
- METHOD DETAILS

- Drug administration
- Brain region abbreviations
- Transsynaptic tracing/cTRIO
- Immunohistochemistry
- Axonal arborization mapping from VTA^DA^ cells
- Slice electrophysiology
- TEVC
- Fiber photometry
- Peri-stimulus time histograms (PSTHs)
- Behavioral assays

- CPP
- Locomotion and locomotor sensitization
- OFT/EPM
- Flupentixol administration
- Carnosic acid administration
- Intracranial self-stimulation
- IVSA
- Auditory fear conditioning
- CRISPR
- Single nucleus RNA sequencing
- QUANTIFICATION AND STATISTICAL ANALYSIS

### KEY RESOURCES TABLE

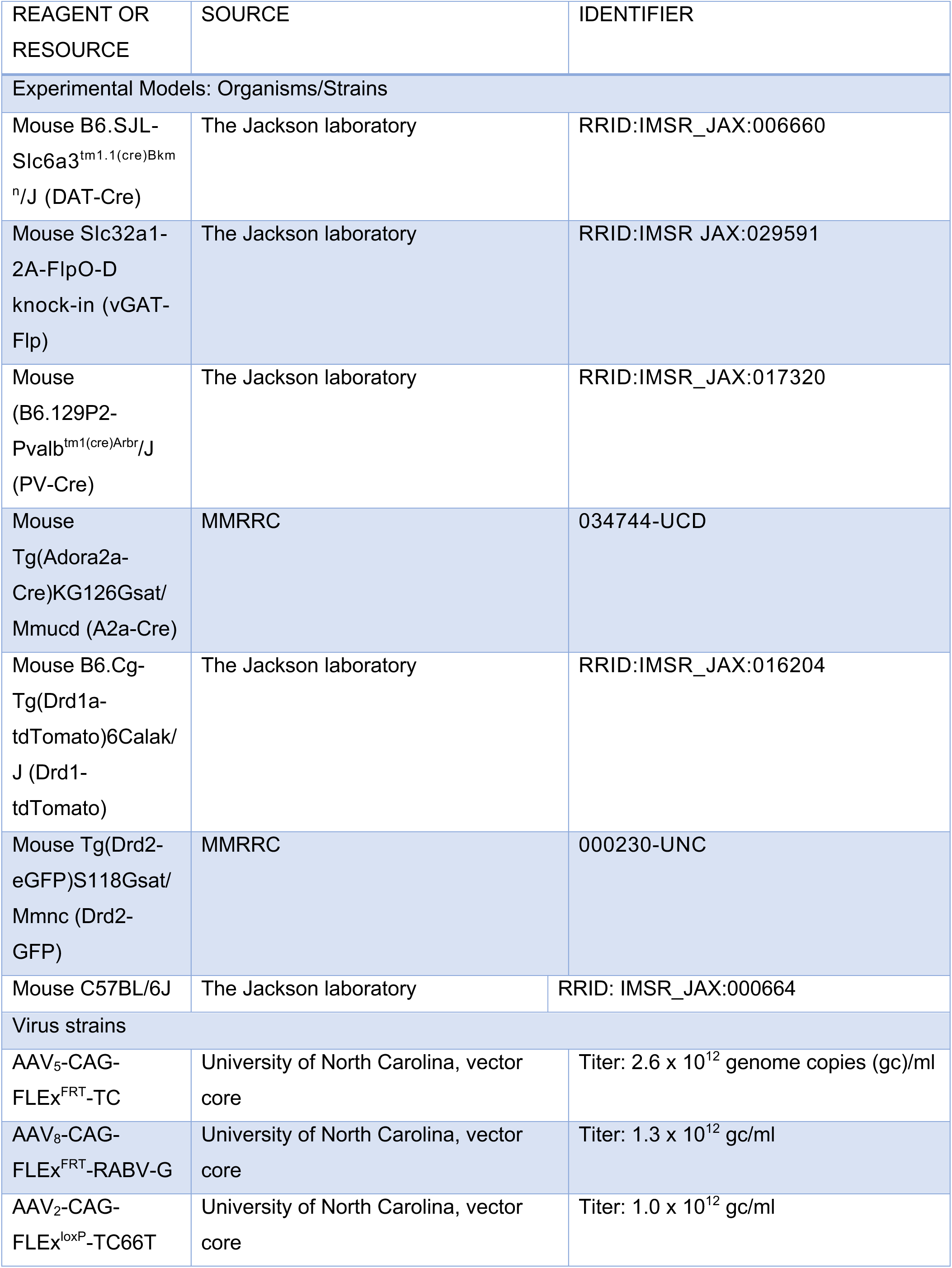
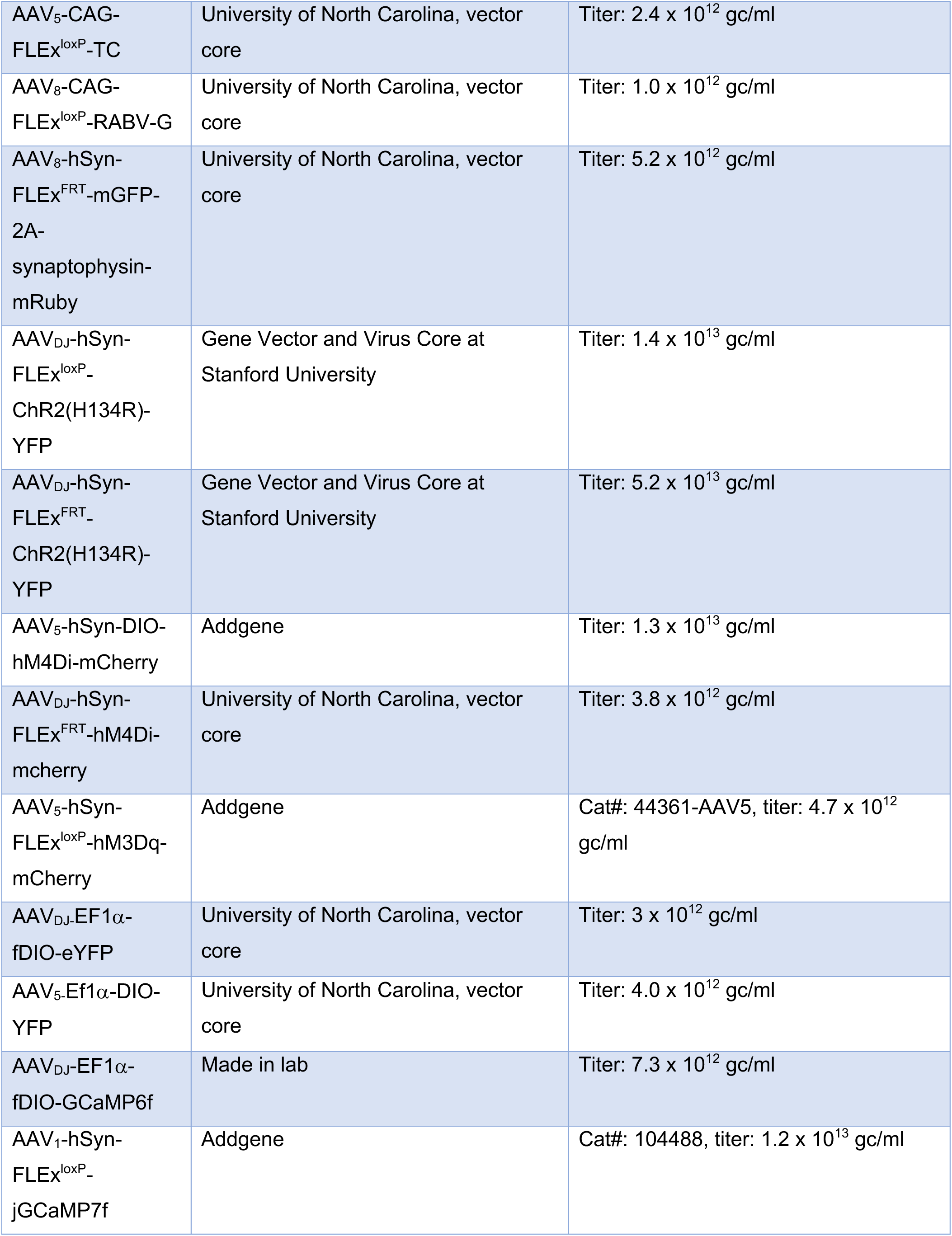
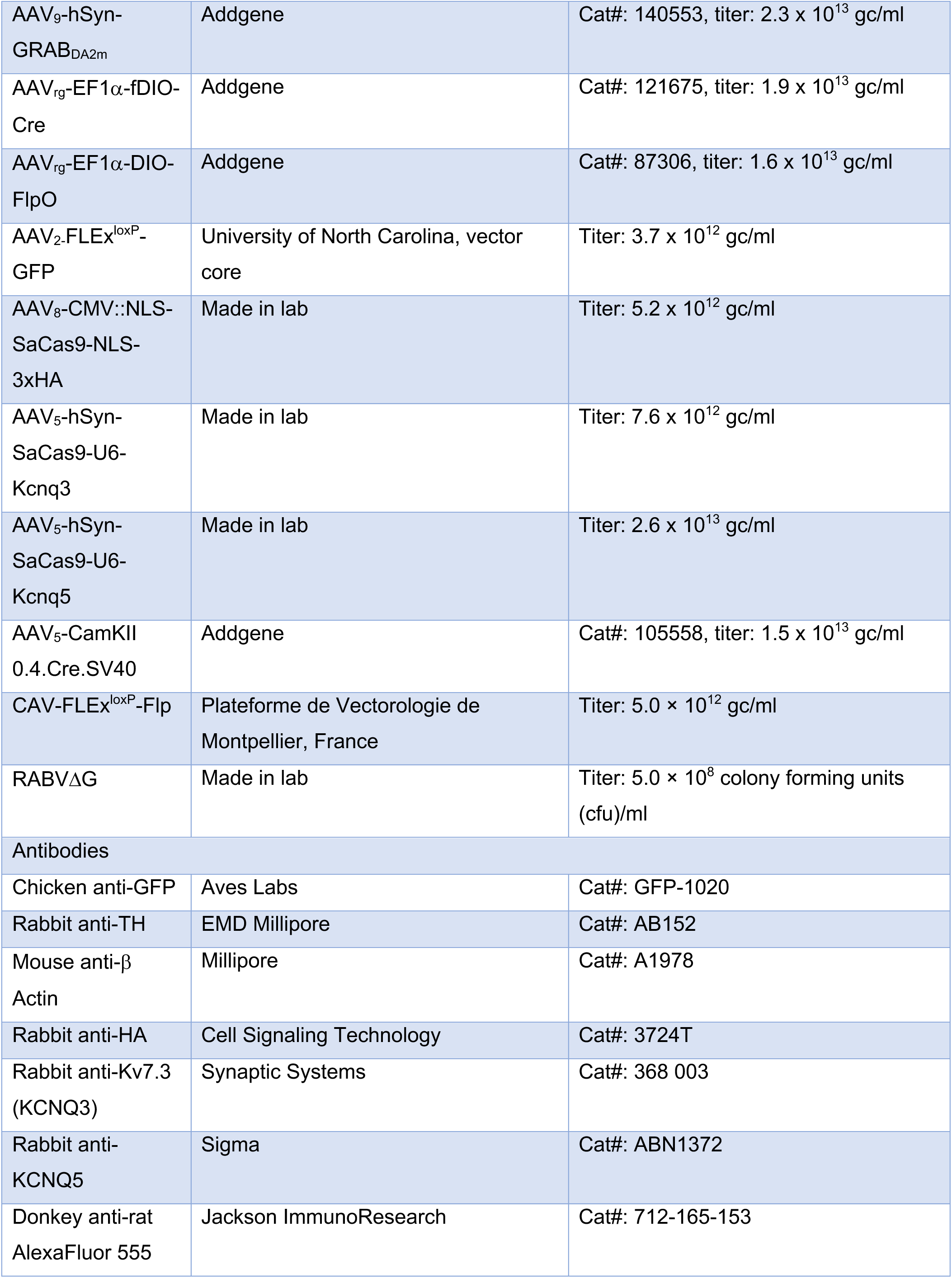
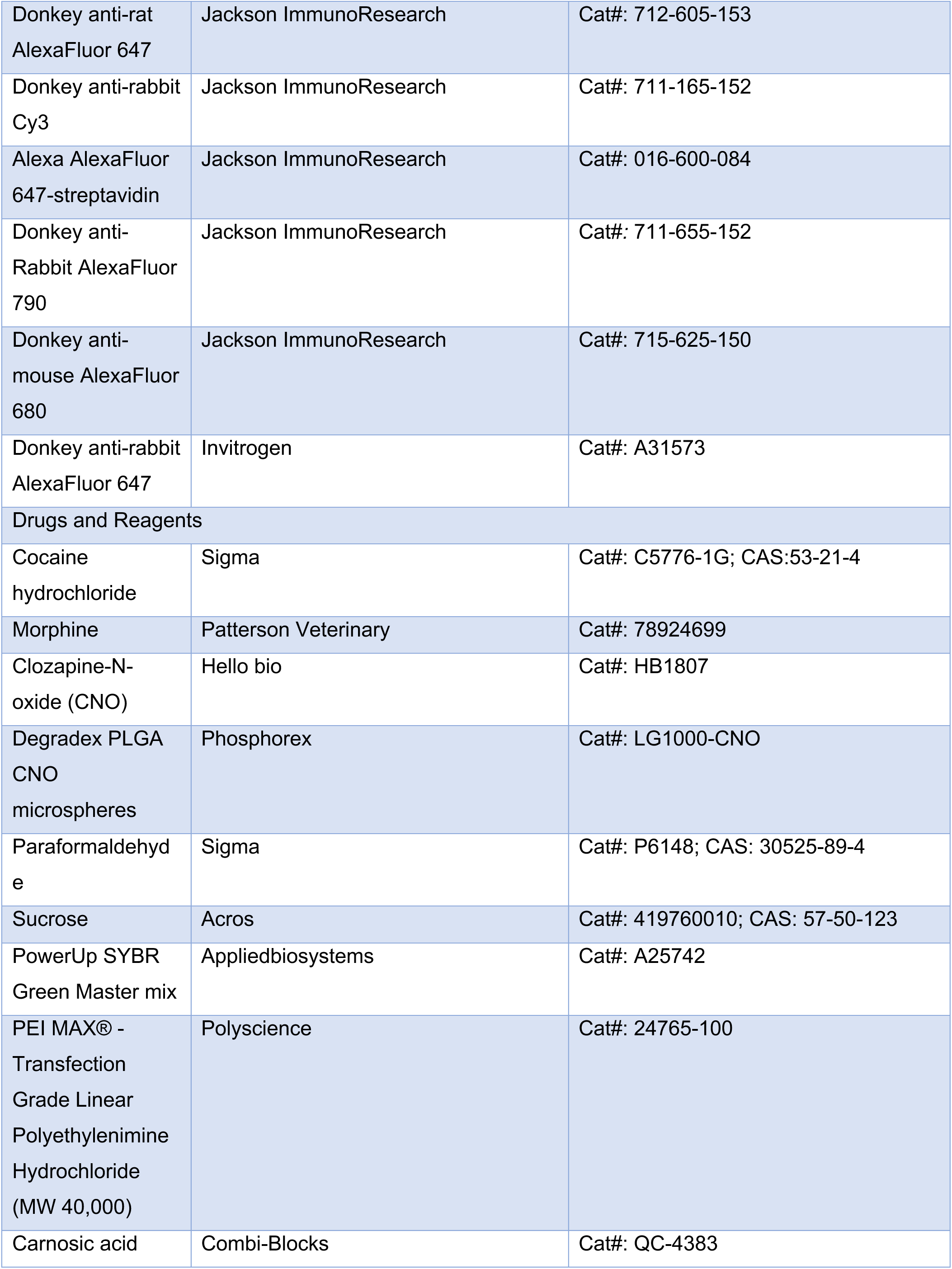
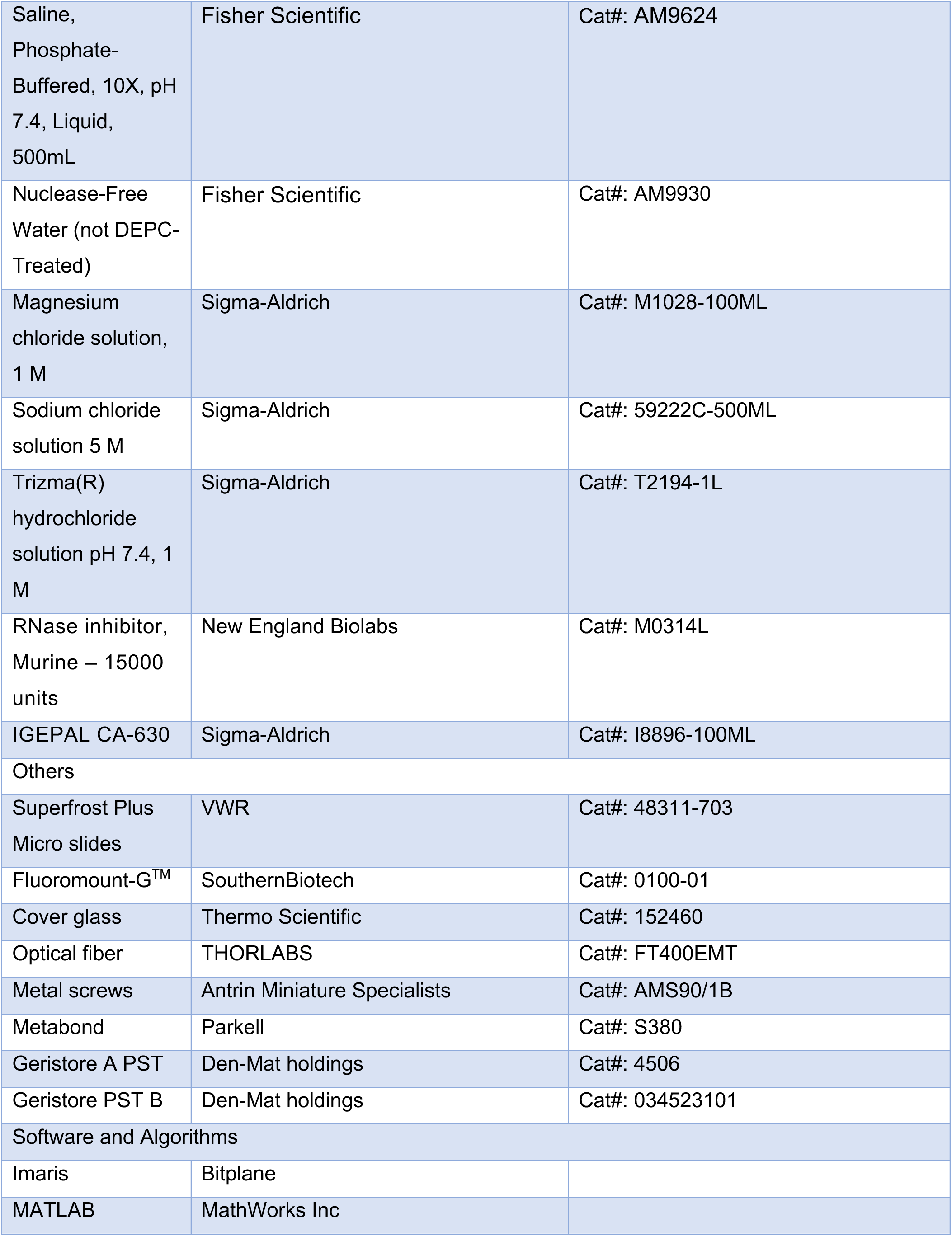
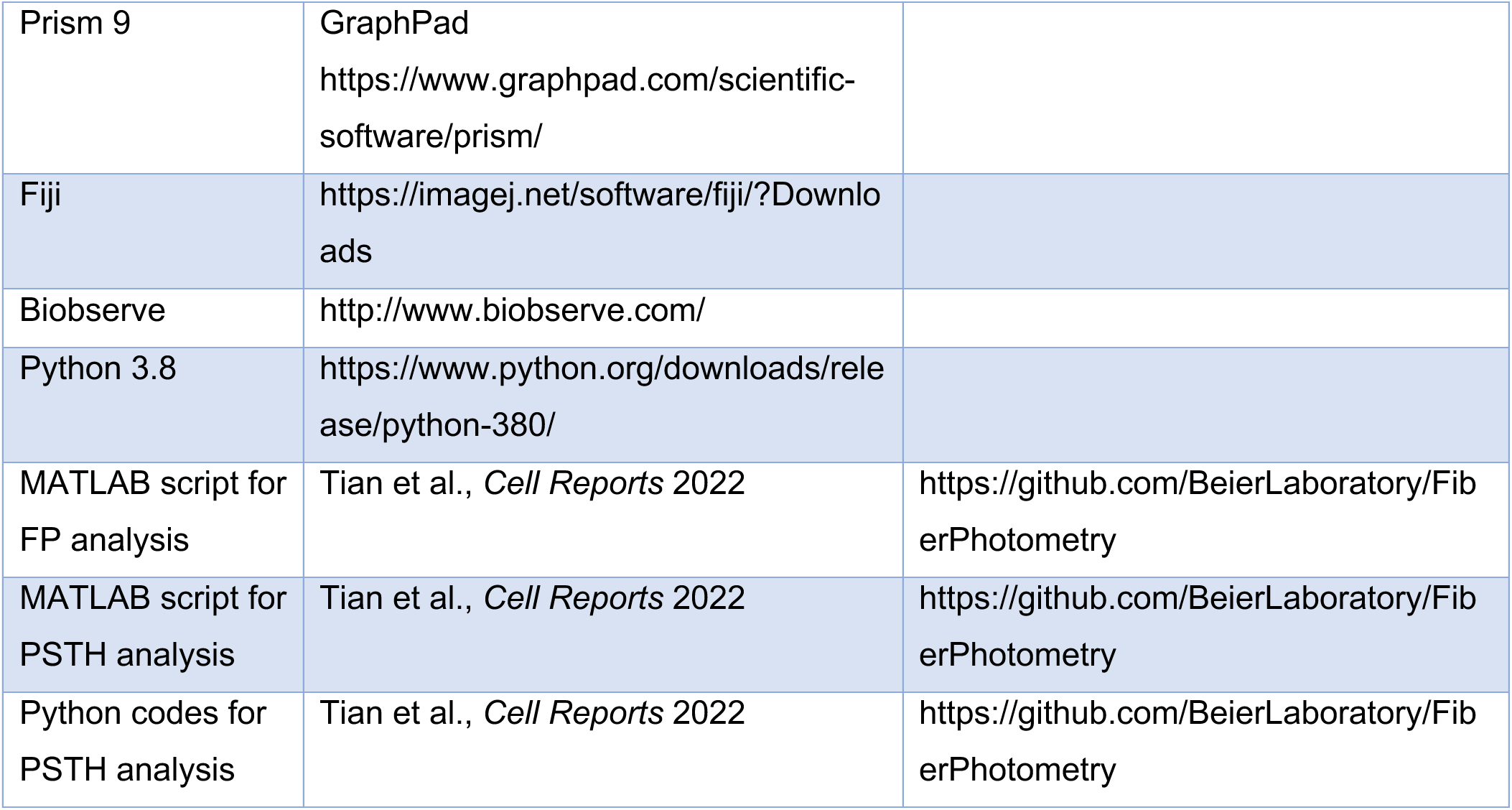

### RESOURCE AVAILABILITY

#### Lead contact

Further information and requests for resources and data should be directed to and will be fulfilled by the lead contact, Kevin Beier.

#### Materials availability

*RABVΔG* made in lab in this study will be distributed upon reasonable request.

#### Data and code availability

- All original code has been deposited at GitHub and is publicly available as of the date of publication. The DOI is listed in the Key Resources Table.
- Raw snRNA-seq counts data and metadata will be uploaded to the Gene Expression Omnibus upon publication.
- Any additional information required to reanalyze the data reported in this paper is available upon reasonable request.

### EXPERIMENTAL MODEL AND SUBJECT DETAILS

#### Animals

Mice were housed on a 12-hour light–dark cycle with food and water *ad libitum*. Males and females from a C57BL/6J background were used for all experiments in approximately equal proportions, except for IVSA experiments, where all males were used. This was done to streamline experiments, and focus on comparisons to results in the literature, which have predominately performed experiments in male mice. All surgeries were done under isoflurane anesthesia. All procedures complied with the animal care standards set forth by the National Institute of Health and were approved by the University of California, Irvine’s Institutional Animal Care and Use Committee (IACUC).

### METHOD DETAILS

#### Drug administration

Cocaine was administered at a dose of 15 mg/kg, CNO at 5 mg/kg, and carnosic acid at one of two concentrations: intraperitoneally at 830 μg//kg and 7.5 mg/kg, and intracranially at 10 μM.

#### Brain region abbreviations

Brain region abbreviations throughout this manuscript are as follows:

Ant. Ctx: Anterior cortex
BNST: Bed nucleus of the stria terminalis
CeA: Central amygdala
DCN: Deep cerebellar nuclei
DStr: Dorsal striatum
DMS: Dorsomedial striatum
DLS: Dorsolateral striatum
DR: Dorsal raphe
EAM: Extended amygdalar area
GPe: Globus pallidus external segment
LDT: Laterodorsal tegmental area
LH: Lateral hypothalamus
LHb: Lateral habenula
MHb: Medial habenula
mPFC: Medial prefrontal cortex
NAc: Nucleus accumbens
NAcCore: Nucleus accumbens core
NAcLat: Nucleus accumbens lateral shell
NAcMed: Nucleus accumbens medial shell
PO: Preoptic area
PVH: Paraventricular nucleus of the hypothalamus
PBN: Parabrachial nucleus
SNc: Substantia nigra pars compacta
SNr: Substantia nigra pars reticulata
STN: Subthalamic nucleus
VP: Ventral pallidum
VTA: Ventral tegmental area
ZI: Zona incerta

#### Transsynaptic tracing/cTRIO

cTRIO experiments were performed as previously described^21^, except that a single injection of cocaine (15 mg/kg) or saline was administered one day prior to RABV injection^72^. We injected 500 nL of CAV-FLEx^loxP^-Flp unilaterally into the NAcLat, NAcMed, DLS, amygdala, or mPFC, and during the same surgery, also injected 500 nL of a 1:1 volume mix of AAV_5_-FLEx^FRT^-TC and AAV_8_-FLEx^FRT^-RABV-G into the VTA, or SNc in the case of CAV injection into the DLS. After 13 days, a single injection of cocaine or saline was given i.p. A G-deleted, GFP-expressing, EnvA-pseudotyped RABV was injected into the VTA or SNc the following day. Animals were sacrificed five days following RABV injection. For flupentixol injections, right after EnvA-pseudotyped RABV was injected into the VTA, 250 nL of 40 mg/mL flupentixol was injected bilaterally into the DMS or GPe. After 5 minutes following animals waking up from isoflurane surgeries, cocaine was administered i.p.

For mapping inputs to GPe^PV^ cells in saline- and cocaine-treated mice, 100 nL of a 1:1 volume mix of AAV_2_-FLEx^loxP^-TC66T and AAV_8_-FLEx^FRT^-RABV-G was injected into the GPe. After 13 days, a single injection of cocaine or saline was given i.p. A G-deleted, GFP-expressing, EnvA-pseudotyped RABV was injected into the GPe the following day. Animals were sacrificed five days following RABV injection.

To identify input labeling changes that occur following an aversive foot shock, in a separate cohort of animals, 13 days following CAV/AAV injection animals were placed into an auditory fear conditioning chamber, as described in the behavior section below. RABV was injected 1 day later, and animals sacrificed five days subsequent, as described above.

Coordinates used for viral injections were (relative to bregma, midline, or dorsal brain surface and in mm):

- Amygdala: AP −1.43, ML 2.5, DV −4.5;
- NAcMed: AP +1.55, ML 0.7, DV −4.0;
- NAcLat: AP +1.45, ML 1.75, DV −4.0;
- DMS: AP +0.75, ML 1.5, DV −2.8
- DLS: AP +0.25, ML 2.5, DV −3.4;
- GPe: AP −0.35, ML 1.75, DV −3.5;
- mPFC: two injections, one at AP +2.15, ML 0.27, DV –2.1 and another at AP +2.15, ML0.27, DV –1.6;
- SNc: AP −3.2, ML 1.2, DV −4.2.
- SNr: AP −3.2, ML 1.3, DV −4.5
- VTA: AP −3.2, ML 0.4, DV −4.2;

#### Immunohistochemistry

The primary antibody chicken anti-GFP (Aves Labs) was used at 1:1,000, rabbit anti-TH (Millipore) at 1:1,000, rat anti-cFos at 1:5,000, rabbit anti-HA at 1:1000, and rat anti-mCherry (ThermoFisher) at 1:2,000. All secondary antibodies were used at a concentration of 1:500.

The fixed brains were dehydrated using 30% sucrose in 1x PBS and were sliced into 60 µm sections using a cryostat. These brain sections were then incubated in a blocking solution [3% NDS in PBST (1xPBS with 0.3% Triton X-100)] at RT for 1 hr, followed by immersion in a primary antibody solution (1% NDS and primary antibody in PBST) at 4°C for 72 hrs. After washing with PBST three times, the sections were incubated in a secondary antibody solution (1% NDS and secondary antibody in PBST) at RT for 2 hrs. The brain sections were then mounted on Superfrost Plus^TM^ microscope slides (Fisher, cat # 1255015). Following drying, the brain sections were incubated in 1xPBS containing DAPI at a dilution of 1:1000 (ThermoFisher, cat# D1306) for 10 min. After the liquid was removed, Fluoromount-G (SouthernBiotech, cat# 0100-01) was applied to the sections and cover glass (Thermo Scientific, cat# 152460) was used to cover the brain sections.

#### Axonal arborization mapping from VTA^DA^ cells

Axonal tracing experiments were performed as previously described^21^. We injected 500 nL of AAV_DJ_-hSyn-FLEx^FRT^-mGFP-2A-Synaptophysin-mRuby unilaterally into the VTA, and CAV-FLEx^loxP^-Flp into the NAcMed, NAcLat, DLS, amygdala, or mPFC. After 2 months, animals were sacrificed, brains were cut with a thickness of 60 μm and immunolabeled for GFP to enhance signal. Sections were imaged on an Olympus IX83 microscope using a 4x objective. The data were quantified as previously described^21^.

#### Electrophysiology

##### Slice electrophysiology

Mice were deeply anaesthetized with isoflurane, and coronal midbrain or GPe slices (300 μm) were prepared after intracardial perfusion with ice-cold artificial cerebrospinal fluid (ACSF) containing (in mM): 230 sucrose, 10 glucose, 25 NaHCO_3_, 2.5 KCl, 1.2 NaH_2_PO_4_, 0.5 CaCl_2_, 7 MgCl_2_, and oxygenated with 95% O_2_/5% CO_2_. After 60 min of recovery, slices were transferred to a recording chamber and perfused continuously at 2–4 ml/min with oxygenated ACSF containing (in mM): 120 NaCl, 10 glucose, 25 NaHCO_3_, 2.5 KCl, 1.2 NaH_2_PO_4_, 2.5 CaCl_2_, 2.3 MgCl_2_ at 30 °C. Patch pipettes (3.8–4.4 MΩ) were pulled from borosilicate glass (G150TF-4; Warner Instruments) and filled with internal solution containing (in mM): 131 CsCH_3_SO_3_, 10 HEPES, 10 glucose, 2 CaCl_2_, 10 EGTA, 5 MgATP, 0.4 NaGTP, 10 phosphocreatine, 5 QX314, and 0.2% biocytin (pH 7.3, 292 mOsm) for voltage clamp recordings. Spontaneous EPSCs and IPSCs were captured continuously. For both spontaneous and evoked recordings in GPe^PV^ cells, EPSCs were recorded at -70 mV, and IPSCs at 0 mV.

For analysis of VTA^DA^→NAcLat cell action potential latency in current clamp, the internal solution contained (in mM): 130 KMeSO_3_, 9 KCl, 0.1 EGTA, 10 HEPES, 4 Mg-ATP, 0.4 Na-GTP, 7.5 Na_2_-phosphocreatine, adjusted to pH 7.3 with KOH, 290–295 mOsm. For VTA^DA^→NAcLat cell recordings, ChR2 was stimulated by flashing 473-nm light (2 ms pulses, 0.1 Hz, 10 mW) through the light path of the microscope using an ultrahigh-powered light-emitting diode (LED) powered by an LED driver (Thorlabs) under computer control. The light intensity of the LED was not changed during the experiments. A dual lamp house adaptor (Olympus) was used to switch between the fluorescence lamp and LED light source. Sweeps were alternated where ChR2 was activated at 20 Hz for 500 ms, throughout the duration of the current step.

To assess neuronal intrinsic excitability, we performed *in vitro* whole-cell patch clamp recordings in acute brain slices under the current clamp mode. Coronal acute brain slices were prepared in ice-cold cutting solution (in mM: 135 NMDG, 10 D-glucose, 4 MgCl_2_, 0.5 CaCl_2_, 1 KCl, 1.2 KH_2_PO_4_, 20 HEPES, and sucrose to adjust the osmolality to 305–310 mmol/kg; pH 7.35; bubbled with 95%O_2_/5% CO_2_). Slices were transferred to a submerged recovery chamber filled with sucrose-based artificial CSF (sACSF; in mM: 55 sucrose, 85 NaCl, 25 D-glucose, 2.5 KCl, 1.25 NaH_2_PO_4_, 0.5 CaCl_2_, 4 MgCl_2_, 26 NaHCO_3_; 300– 305 mmol/kg; bubbled with 95%O_2_/5% CO_2_). The recovery chamber stayed in a 37°C water bath for 30 min before being held at room temperature. For whole-cell patch clamp recordings, brain slices were placed in a submerged recording chamber filled with circulating ACSF (in mM: 126 NaCl, 10 D-glucose, 2 MgCl_2_, 2 CaCl2, 2.5 KCl, 1.25 NaH_2_PO_4_, 1.5 sodium pyruvate, 1 L-glutamine; 295–300 mmol/kg; bubbled with 95%O_2_/5% CO_2_) at room temperature. Fluorescently labeled neurons in the GPe or SNr were patched with fine-tip glass pipettes (4-6 M) filled with K-methanesulfonate (K-met) based internal solution (in mM: 130 K-met, 10 HEPES, 5 KCl, 5 Na-phosphocreatine, 2 MgCl_2_, 2 Mg-ATP, 0.6 EGTA, and 0.3 Na-GTP; pH 7.25; 285-300 mOsm). The patch pipettes for GPe^PV^ cells in the KCNQ3/5 KO mice were filled with biocytin for post-hoc identification.

Whole-cell recordings were achieved under voltage-clamp at -70 mV. Cells were then switched to current clamp with no holding current. Resting membrane potential was measured when cells were stabilized. A series of other intrinsic excitability measurements were done while cells are held at -70 mV: 1) Current-spike curves, determined by injecting a series of current steps (0-500 pA at 50 pA increments, 500 ms); 2) Rheobase, calculated as the current needed to generate the first action potential, by injecting a series of 500 ms current steps at 5 pA increments; and 3) Input resistance, calculated as Rin = V / I, at I = -50 pA.

Clampfit 11.2 (Molecular Devices Inc., CA) was used for processing slice electrophysiology data. Action potential measurements were made using the “Threshold Search” mode under “Event Detection”. Detected events also went through a visual inspection to exclude any outliers or noise. For all analyses, normality was first determined using the Shapiro-Wilk test. An unpaired t-test was used for normally distributed data and a Mann–Whitney test was used for data that was not normally distributed. For comparisons between the current-spike curves, two-way repeated measures ANOVA (RM ANOVA) was used to test for statistical differences between conditions.

For all recordings, input resistance and access resistance were monitored continuously throughout each experiment; experiments were terminated if these changed by > 20%. Labeled neurons were visualized with a 40x water-immersion objective on an upright fluorescent microscope (BX51WI; Olympus) equipped with infrared differential interference contrast video microscopy and epifluorescence (Olympus).

#### Validation of CNO microspheres

CNO microspheres were synthesized to enable slow release of CNO after a single infusion. Degradex PLGA CNO microspheres were custom ordered from Phosphorex. Beads of target mean diameter 1 μm were dissolved in 0.5% trehalose at a concentration of 5 mg micro-sphere per ml. The estimated CNO loading efficiency was 5%, the estimated burst release was 50%, and the estimated release time was 7 days. The target concentration of CNO release at a steady state was 100 pg/h^19,73^.

##### Acute slice preparations

To assess the effect of CNO microspheres on neuronal excitability, we performed whole-cell patch clamp recordings in SNr cells expressing hM4Di following injections of either CNO microspheres or saline. Briefly, 500 nL of a 1:1 volume mix of AAV_5_-CamKII 0.4.cre.SV40 and AAV_5_-hSyn-DIO-hM4Di-mCherry was injected unilaterally into the SNr (in mm, AP: -3.2; ML: 1.3; DV: -4.5) of wild-type mice. Three to four weeks later, 500 nL of either saline or CNO microspheres was injected in the same location. Approximately 3 days following microsphere injection, acute brain slices were prepared, and mCherry-labeled cells expressing hM4Di in the SNr were patched. Neuronal intrinsic excitability of recorded SNr cells in the saline and CNO groups was quantified with the same method described above (see the method section on slice electrophysiology).

##### In vivo

500 nL of AAV_5_-hSyn-DIO-hM4Di-mCherry was injected into the GPe on the right hemisphere and AAV_2_-FLEx^loxP^-GFP into the GPe on the left hemisphere of PV-cre mice. One month later, 500 nL of CNO microspheres were infused into SNr bilaterally. Four days later, cocaine or saline was administered i.p. One hour later, mice were subjected to transcardial perfusion with 1xPBS followed by 4% formaldehyde in 1xPBS. Following extraction, the brains were post-fixed in 4% formaldehyde for 24 hrs and sliced into 60 μm sections. The brain sections went through the immunohistochemistry procedure as descried above with the rat anti-cFos antibody and the brain sections consisting of SNr and VTA were then mounted. Images of the midbrain were captured using an Olympus IX83 microscope, and manual cell counting was performed on the acquired images using FIJI.

##### Channel subunit cRNA preparation, *Xenopus laevis* oocyte injection, and Two-Electrode Voltage Clamp (TEVC)

cRNA transcripts encoding KCNQ 2, 3, 4 and 5 channels were generated *in vitro* and expressed in stage V and VI *Xenopus laevis* oocytes as previously described^39^. TEVC was performed at room temperature and oocytes expressing KCNQ channels recorded 2-5 days post-injection in 4 mM extracellular KCl bath solution with recording electrodes of 1-2 MΩ resistance. Conductance-voltage curves were generated as previously described^39^. Data was analyzed using Clampfit (Molecular Devices).

##### Fiber photometry

Fiber photometry experiments were performed as previously described^18^. To measure activity in GPe^PV^ cells, 500 nL of AAV_1_-hSyn-FLEx^loxP^-jGCaMP7f was injected into the GPe. A 400 μm diameter, 0.39NA optical fiber was implanted at the same location. For GPe^PV^→ventral midbrain neurons, 500 nL of AAV_rg_-DIO-FLPo was injected into the ventral midbrain, 500 nL of AAV_DJ_-EF1α-fDIO-GCaMP6f was injected in the GPe, and fibers implanted over the GPe of PV-Cre mice. For VTA^DA^→NAcLat terminal recordings, 500 nL of AAV_1_-hSyn-FLEx^loxP^-jGCaMP7f was injected into the VTA, and fibers were implanted over the NAcLat of DAT-Cre mice. For DMS^D2^ cells, AAV_1_-hSyn-FLEx^loxP^-jGCaMP7f was injected into the DMS, and a fiber was implanted over the DMS of A2a-Cre mice. For SNr^GABA^ cells, 500 nL of AAV_DJ_-EF1α-fDIO-GCaMP6f was injected into the SNr, and a fiber was implanted over the SNr of vGAT-Flp mice. For GRAB_DA2m_ recordings, AAV_9_-hSyn-GRAB_DA2m_ was injected into the DMS, and a fiber implanted over the DMS unilaterally while either AAV_5_-hSyn-DIO-hM4Di-mCherry or AAV_5_-DIO-YFP was injected bilaterally into the GPe of PV-Cre mice.

Implants were placed approximately 0.2 mm above the site of viral injection and were secured to the skull with metal screws (Antrin Miniature Specialists), Metabond (Parkell), and Geristore dental epoxy (DenMat). Mice were allowed to recover for at least 4 weeks before experiments began.

Fiber photometry recordings were made using previously described equipment^74^. Mice were run through an injection/behavior timeline shown in Figure 4B. In brief, 465 nm and 405 nm excitation light were controlled via a RZ5P real-time processor (Tucker Davis Technologies) using Synapse software and were used to stimulate Ca^2+^-dependent and isosbestic emission, respectively. All optical signals were bandpass filtered with a Fluorescence Mini Cube FMC4 (Doric) and were measured with a femtowatt photoreceiver (Newport). Signal processing was performed with MATLAB (MathWorks Inc.). Signals were first motion corrected by subtracting the least squares best fit of the control trace to the calcium signal. Data points containing large motion artifacts were then manually removed. To assess neural activity, we quantified the time spent above threshold, which was set at 2.91 times the median absolute deviation (MAD) of each day’s recording, a value that equates to the 95% confidence interval for Gaussian data^75^.

Area under curve (AUC) of fiber photometry traces were computed by integrating across the GCaMP signals. The MATLAB implementation of the trapezoidal rule (trapz) was used to integrate the normalized GCaMP signal, which had been subtracted by its minimum value. AUCs for hM4Di and YFP were collected for different days of cocaine administration and compared with a paired t-test.

##### Peri-stimulus time histograms (PSTHs)

PSTHs were generated using timestamps corresponding to certain behavioral events. The video recorded by Synapse software was rerun using Biobserve, which returns a frame-by-frame record of coordinates corresponding to the nose, center of body, and the base of the mouse’s tail. After defining the borders of each behavior arena within Biobserve, a custom Python script was used to generate a list of timestamps corresponding to selected behavioral events. These events are as follows: for both the CPP pre-test and post-test, the moment the mouse enters either the saline-paired or cocaine-paired chamber; for locomotion, the moment the mouse initiated or ceased movement. The coordinate corresponding to the mouse’s body center was used for all calculations. For locomotion, motion was calculated by taking the distance the center body coordinate moved from frame to frame. The velocity threshold for still was lower than 12.5 cm/s. The state of still was defined as the duration of still lasting for over 20 s. A second MATLAB script aligned each behavioral timestamp with the raw photometry trace and resampled the Biobserve coordinates to match the sampling rate of the fiber photometry data. The PSTH curve was then generated by charting the mean Z-score during the three-second pre- and post-event time intervals centered around each behavioral event.

To quantitatively determine which conditions had the most similar PSTH curves across all tests, the PSTH curves for each test were stitched into one table by condition and Uniform Manifold Approximation and Projection (UMAP) was used to visualize them in two dimensions. Analysis of PSTH curves with UMAP was restricted to -0.5 to 1.5 s surrounding each event. This 4x2010 table was reduced to two dimensions with UMAP and conditions with similar PSTH profiles were identified (Figure S5G). In addition, since individual UMAP embeddings are stochastic depending on initial seeding conditions, we examined the average relative Euclidian distance between the same data points in 20 UMAP embeddings of the same data and plotted the averaged results as a correlogram (Figure S5H).

#### Behavioral assays

**CPP:** To test for drug-induced CPP, animals were first tested in a single drug pairing, two-chamber CPP test. Each chamber was given different wall contexts. On the first day, animals were initially placed into the right chamber, and allowed to freely explore both chambers for thirty minutes (pre-test). On the second day, animals were saline-conditioned to the chamber in which they spent more time during the pre-test, and on the third day, cocaine-conditioned (15 mg/kg) to the chamber in which they spent less time during the pre-test. On the fourth day, animals were again initially placed into the right chamber, and allowed to explore freely (post-test). In tests where hM4Di was used along with YFP-expressing controls, 5 mg/kg CNO was injected thirty minutes before the beginning of both the cocaine and saline pairings. CPP scores were computed as the time spent in the drug paired chamber posttest-pretest. Each session was 30 minutes, except for in optogenetics experiments, which were 15 minutes. Mice were subject to exclusion if the time spent on one side during the CPP pretest was less than 600 seconds (except for in optogenetic experiments, as the test was shorter).

##### Locomotion and locomotor sensitization

To test sensitization, animals were habituated to open field boxes equipped with motion tracking for two days (receiving saline injections before each session). Boxes contained polka dotted contexts on the walls for the duration of the sensitization testing. Animals were then injected with cocaine immediately before entry into the open-field boxes, for five consecutive days, for 30 minutes each. For assessing potential CNO-induced changes in locomotion, an extra day was introduced following the two days of habituation, but prior to cocaine injection. Mice were excluded if data from >1 day of the task were missing.

For behavioral experiments where CNO microspheres were injected (e.g., Figure S1A-I, Figure 2E-J), a modified behavioral protocol was used for CPP and locomotor sensitization, as shown in Figure S1B. This was done because the slow-release CNO microspheres release CNO for approximately one week^14,18,19^; therefore, all cocaine injections were administered within one week following injection of the microspheres. Briefly, the CPP pretest and two habituation days in the open field were initiated one month following viral injection. Following these three days, CNO microspheres were injected into the ventral midbrain (Figure S1, S3) or DMS or GPe (Figure 2). After allowing one day for recovery, the saline pairing in the CPP task was performed in the morning to one chamber, and the cocaine pairing to the other chamber in the afternoon. The following morning, the CPP post-test was performed. In the afternoon of the same day, cocaine was administered, and the animals were placed into the open field, where the first day of locomotor sensitization testing was conducted. Sensitization experiments then continued each day for a total of five days. In Figure S1, following a ten-day abstinence period, animals were tested for the time spent in the open arms of the elevated plus maze and in the center of an open field. Following these tests, the CPP task was then repeated, as was sensitization, to assess whether these behaviors could be elicited by cocaine in the absence of CNO, as the microspheres were depleted of CNO by this time.

##### OFT/EPM

In some cases, after waiting ten days following the final drug injection, we then tested the animals in the open field test and elevated plus maze for anxiety-like behaviors. For the OFT the time spent in the center square (1/3 of the total area of the arena) during a five-minute test period was calculated. The following day, animals were tested in the EPM. Animals were placed at the end of one of the open arms, facing outwards, at the beginning of the trial. The percentage of time spent in the two open arms during the five-minute testing period was quantified. OFT experiments were performed at an illuminance level ∼300 lux, and EPM ∼25 lux.

For fiber photometry experiments, behavioral assays were performed exactly as described above. For chemogenetic inhibition experiments to test the necessity of defined cell populations in behavioral adaptation, to inhibit projection-defined subsets of ventral midbrain DA cells (Figure 1, S2), we used a viral-genetic intersectional method. We injected 500 nL of CAV-FLEx^loxP^-Flp bilaterally into the NAcMed, NAcLat, DLS, or amygdala, or 2 μL into the mPFC to target DA neurons projecting to each of these sites. During the same surgery, 500 nL of AAV_5_-hSyn-FLEx^FRT^-hM4Di (or YFP) was injected bilaterally into the ventral midbrain. Animals were allowed 2 weeks to recover. 5 mg/kg CNO was injected i.p. 30 min before each injection of 15 mg/kg cocaine (or saline during CPP pairing and day 3 of locomotor habituation to test for CNO effects on basal locomotor activity).

For chemogenetic activation experiments, mice expressing hM3Dq or YFP in DMS^D2^ cells were injected with CNO 30 minutes before testing CPP and locomotor behavior following cocaine administration.

#### Effects of flupentixol on cocaine CPP and locomotor sensitization

Cannulas were implanted in the GPe or the DMS. To prepare mice for intracranial implantations, we anesthetized mice with isoflurane and used a stereotaxic frame to secure the animals and ensure precise placement of the bilateral guide cannulas. To prepare the skull for implantation of the cannulas, a 1.4-mm-diameter drill bit was used to create a craniotomy (AP: -0.35 mm, ML: ±1.75 mm), and the stereotaxic manipulator was used to lower the sterile surgical-grade steel cannulas with their attached base plate (P1 Technologies) to the proper DV coordinates from the brain surface. For the GPe, we used the coordinates AP: -0.35 mm, ML: ±1.75 mm, DV: 3.5 mm. For the DMS, we used two separate coordinates, 10 mice for each: AP: +0.35 mm, ML: ±1.75 mm, DV: -3.5 mm and AP: +0.75 mm, ML: ±1.5 mm, DV: - 2.8 mm. The bilateral guide cannulas were secured to the skull with metal screws (Antrin Miniature Specialists), Metabond (Parkell), and dental epoxy (DenMat). Following proper placement and secure attachment, dummy cannulas (P1 Technologies) were inserted to seal the open ports that allow direct access to the brain. Immediately following surgical implantation, flunixin (Vetameg, 2.5 mg/kg of body weight) and antibiotic (Baytril, 5 mg/kg of body weight) were injected i.p. Implanted mice were then singly housed and allowed to recover for 10 days prior to any infusions.

For infusions, on experimental days mice were briefly anesthetized using isoflurane (3-4% in O_2_) to induce the animals and anesthesia was maintained using the lowest dose of isoflurane (1-1.5%) to sustain immobility during infusions. Dummy cannulas were removed and using a micro-infusion injector (WPI, Nanofil) with a syringe (Hamilton syringe 600 series, Hamilton Company) connected directly to tubing (C312VT/PKG .027X.045 vinyl tube, P1 Technologies) which was connected to the single internal cannulas, 250 μL of 40 mg/mL of flupentixol in sterile 0.9% saline, or 0.9% sterile saline, was infused directly into both of the cannulas simultaneously at a rate of 1.6 nL/sec. After five minutes following completion of infusion, bilateral cannulas were carefully removed. We then removed the animals from the stereotaxic frame and allowed the mice to recover for thirty minutes before beginning behavioral testing. All custom-made cannula parts were obtained from P1 Technologies.

In the CPP task, experimental mice received an infusion of flupentixol, while control mice received an infusion of 0.9% saline. Experimental and control mice were then i.p. injected with 0.9% saline in the first pairing, and 15 mg/kg cocaine in the second pairing, each 30 minutes following cannula infusions. For locomotor testing, following habituation days, the implanted mice then underwent five consecutive days of cannula infusions of flupentixol or saline 30 minutes prior to i.p. injected cocaine to assess their locomotor behavior each day.

#### Carnosic acid administration prior to CPP and sensitization

To test the effects of carnosic acid on CPP and sensitization, two methods were used. First, carnosic acid (2.5 mg/kg, dissolved in 3% DMSO in 0.9% saline) was given i.p. 30 minutes prior to cocaine administration. Animals were then tested for CPP and locomotor sensitization as detailed elsewhere. The other method was delivering carnosic acid directly into the GPe via intracranial infusion. These were performed in the same was as for flupentixol infusion experiments, except that for each infusion, either 500 nL of carnosic acid (10 μM carnosic acid in 0.9% sterile saline and 3% DMSO) or vehicle (0.9% sterile saline and 3% DMSO) was infused.

#### Intracranial self-stimulation (ICSS)

Standard stereotaxic procedures were used to infuse virus, and optical fibers were implanted over the GPe under isoflurane anesthesia as described elsewhere. AAV_5_-Ef1α-DIO-YFP or AAV_DJ_-hSyn-FLEx^loxP^- ChR2(H134R)-YFP were infused into the GPe at a rate of 100 nL/min and the injection needle was left in place for 5 minutes after each infusion before it was slowly removed. Following surgery, mice were returned to group housing after being allowed to recover on a heating pad. During the same surgery, mice were implanted with optical fibers (made in-house with 0.39 NA, 200 μm diameter optical fiber, Thorlabs) targeted just dorsal to the GPe. Implants were secured to the skull with metal screws and Geristore dental epoxy. Mice were allowed to recover for >10d before behavioral procedures commenced.

Experimental sessions were conducted in operant conditioning chambers (24 cm W x 20 cm D x 18 cm H, Med Associates Inc.) contained within sound-attenuating cubicles. The left side of the chamber was fitted with 5 nosepoke ports, each with an LED light at the rear. A video camera (Med Associates Inc.) was positioned at the top of the sound-attenuating cubicle that provided a top-down view of the entire conditioning chamber. Prior to behavioral sessions, mice were gently attached to patch cables made in-house with optical fiber (0.39 NA, 200 μm diameter, Thorlabs) via a ceramic split sleeve (Precision Fiber Products). The patch cables were also connected to bilateral rotary joints (Doric Lenses), which permitted free rotation while transmitting blue light from an upstream 473 nm blue laser (Laserglow). Optical stimulation was controlled by a computer running Med PC IV software (Med Associates Inc.), which also recorded responses at all nosepoke ports and initiated and terminated video recording.

Prior to the first behavioral session, mice were familiarized with cereal treats (Fruit Loops, Kellogg) in their home cage. On the first training day, all nosepokes were baited with crushed cereal treats to facilitate initial investigation. The start of the session was indicated to the mouse by the illumination of a white house light. Session length was 60 min, during which time mice were free to respond at any nosepoke port. 4 ports were designated “active” ports, and a response at these ports produced 2 s of optical stimulation at a particular frequency (10, 20, 30, or 50 Hz); the LED at the back of the corresponding port was concurrently illuminated to provide a visual cue signaling the presence of optical stimulation. Responses made within the 2 s stimulation period were recorded but had no consequence. Responses at a 5th “inactive” nosepoke port were recorded but did not result in either optical stimulation or cue light presentation. Testing occurred once per day for 3 days. Peak light output during photostimulation was estimated to be ∼2.75 mW at the tip of the implanted fiber (∼22 mW/mm^2^), and ∼5 mW/mm^2^ at the targeted tissue 200 µm from the fiber tip. The power density estimates were based on the light transmission calculator at www.optogenetics.org/calc.

#### Intravenous self-administration (IVSA)

##### Animals

All mouse studies were conducted in accordance with the National Institutes of Health Guidelines for Animal Care and Use and with approved animal protocols from the University of California Irvine Institutional Animal Care and Use Committee. Prior to self-administration testing, mice were group-housed, 5 animals per cage. Mice were maintained in a temperature (21±1°C) and humidity (55±10%) controlled room with a 12-h light:12-h dark cycle (light on between 19:00 and 07:00h). Following jugular vein catheter implantation, mice were single-housed and kept on the same conditions as above. At the beginning of the IVSA experiments, animals were 10–12 weeks old and 21–30 g body weight. All IVSA behavioral experiments were conducted during the dark time period and the time of self-administration was held constant for individual animals throughout the duration of the study to control for the influence of circadian effects. Tap water and standard laboratory chow were supplied ad libitum, except during testing. All experimental procedures and animal husbandry were conducted according to standard ethical guidelines (Association for the Assessment and Accreditation of Laboratory Animal Care International (AAALACi) Guidelines on the Care and Use of Laboratory Animals) and approved by the University Laboratory Animal Resources (ULAR) at UCI.

#### Jugular Catheterization Surgery

Indwelling catheters (Instech Laboratories, Inc., Plymouth Meeting, PA) were surgically implanted into the right jugular vein of adult male C57BL/6J mice. Briefly, mice were anesthetized with an i.p. injection of a freshly prepared ketamine/xylazine (12/1 mg/mL, 10 mL/kg) solution. Polyurethane catheters (Part# C20PU-MJV1301; Instech, Plymouth Meeting, PA) were cut 3.5 cm from the collar away from the tip to remove excess catheter tubing and secured to a vascular access button (Part# VABM1B/25; Instech) prior to surgical implantation. The catheters with a collar 1.3 cm from the round tip were inserted into the right jugular vein and threaded under the skin of the shoulder to the dorsum-mounted vascular access button. Vannas micro-dissecting spring scissors (Part# RS-5610; Roboz Surgical Instrument Co., Inc., Gaithersburg, MD) were used to carefully make a micro-incision of the right jugular vein for catheter insertion. The attentive post-operative care was performed by the surgeon for 7 consecutive days post-surgery. To prevent possible infection, on post-surgery days 1-3, all mice received a once-daily subcutaneous injection of the antibiotic Amikacin (10 mg/kg). To ease the animals’ perceived discomfort during recovery by reducing pain, fever, and inflammation, all animals received a once-daily subcutaneous injection of the NSAID Ketoprofen (1 mg/kg) on post-surgery days 1-7. To keep the catheters non-obstructed and prevent infection for the duration of the study, the catheters were flushed once to twice daily with 50 μL of a mixture of 30 U/ml heparin and 1 mg/mL gentamicin solution.

#### Jugular Catheter Patency Testing

All mice underwent patency testing on post-surgery day 7 to confirm that the catheters were free-flowing and properly placed. Patency and optimal placement of the jugular catheters were determined by obvious loss of muscle tone and rapid sedation within 3 seconds from an infusion of 20-30 μL of a 15 mg/mL ketamine solution in sterile saline directly into the injection port of the back-mounted vascular access button which if properly placed and patent, should have fed directly into the jugular vein of the animals and caused sedation. If sedation was not witnessed in under 3 seconds or resistance was met, the animals were considered non-patent. Additional patency tests were performed when catheter patency appeared compromised and at the completion of the study to verify catheter patency. Mice that did not pass the patency tests at any point during the experiment were excluded from the study.

#### Drug Treatment

Doses were calculated based on the weight of cocaine and carnosic acid salts and were not adjusted for fractional base weight. For IVSA studies, carnosic acid (830 μg/kg or 7.5 mg/kg) was i.p. injected 30 minutes prior to the animals being individually placed in the operant chambers. The volume of the i.p. injections was based on individual animal weight. Cocaine (0.5 mg/kg/infusion) was self-administered intravenously in a volume of 14 μL over a 2 s period to animals with verified patent indwelling jugular catheters according to the assigned acquisition protocol and their drug-paired active lever pressing. The drug solutions were formulated to deliver the appropriate dose of cocaine to each mouse based on the groups’ average body weight. For example, for a group of mice with a mean weight of 30 g requiring a 0.5 mg/kg/infusion cocaine dose, a 1.07 mg/mL cocaine infusion solution would be prepared based on the following calculation: 0.5 mg/kg x 0.030 kg/mouse x each infusion/0.014 mL. The weight of each animal was rounded to the nearest gram, and infusion solutions were prepared accordingly.

#### Self-Administration

Following a 7–10 day post-surgical recovery period, mice that passed the patency test were i.p. injected with vehicle, 830 μg/kg carnosic acid, or 7.5 mg/kg carnosic acid, and then trained to IV self-administer cocaine (0.5 mg/kg/infusion) paired with a flashing cue light above the active lever, lever insertions, and retractions and the sound of the syringe pumps actuated by active lever pressing in operant chambers (Med Associates, Inc., Fairfax, VT). The active, reward lever was distinguished by a solid cue light above the active lever. Cocaine rewards were delivered by infusions from a single-speed syringe pump (Med Associates) in a volume of 14 μL over a 2 s period. Each infusion was followed by a 40 s time-out period during which no additional reinforcements were provided. This time-out was implemented to prevent the dangerous health consequences of excessive cocaine consumption in a short duration of time. For the first 20 s of the assigned time-out period, the cue light flashed at a frequency of 1 Hz (i.e., 1 s on and 1 s off) and the levers remained accessible, but additional lever presses during the time-out period had no programmed consequences. For the remaining 20 s (of the 40 s total of the time-out period), the levers were retracted and remained inaccessible. Throughout the acquisition trials, 1 hr daily sessions were used to train the mice to discriminate the active drug-delivering lever from the inactive non-drug-delivering lever, and the fixed lever response to reinforcement ratio (i.e., fixed ratio, FR) was FR1. During the acquisition trials, mouse presses of the active lever (during the time-in period) resulted in an infusion of cocaine while inactive lever presses were recorded but had no programmed consequences. Animals that did not show consistent lever pressing and at least 75% preference for the active lever over 3 consecutive sessions were tested for patency. Animals with confirmed non-patent catheters were removed from the study.

#### Auditory fear conditioning

Mice were first habituated to the auditory fear conditioning chambers. Mice were individually placed in the chamber (Coulbourn Instruments) located in the center of a sound attenuating cubicle. The conditioning chamber was cleaned with 10% ethanol to provide a background odor. A ventilation fan provided a background noise at ∼55 dB. After a 2 min exploration period, three 2 kHz, 85 dB tones were played, for 30s each, with a 90s interval between them. Following these initial tones, three subsequent tones were pained with a 1s, 0.75 mA foot shock. The foot shocks co-terminated with the tone. The mice remained in the chamber for another 60 s before being returned to the home cages. RABV injections were then performed the following day.

#### Knockdown of KCNQ3 and KCNQ5 using CRISPR to examine carnosic acid specificity Generation of Kcnq3 and Kcnq5 sgRNAs

Short guide RNA sequences for saCas9 CRISPR-mediated knockdown of endogenous KCNQ3 and KCNQ5 were designed using CRISPOR^76^. We designed three candidate guides per target and validated each functionally using Western blot of endogenous KCNQ3 and KCNQ5 in mouse primary hippocampal cultures. The KCNQ3 gRNA targets exon 2 (NM_152923.3): Upper: 5’- caccgTTGCTTTGAGGATCTGGGCTG-3’ and Lower: 5’- aaacCAGCCCAGATCCTCAAAGCAAc-3’. The KCNQ5 gRNA targets exon 4 (NM_001160139.1): Upper: 5’- caccgTGACGTGGCGAAAATATTACC-3’ and Lower: 5’- aaacGGTAATATTTTCGCCACGTCAc-3’. The guides were annealed and inserted into a custom AAV plasmid, AAV hSyn saCas9-U6-sgRNA (gift from Dr. Matthew Kennedy, CU Anschutz). Briefly, this plasmid was generated from a parental AAV hSyn FLEX-saCas9-U6-sgRNA^77^. The saCas9 and U6-sgRNA were separately PCR amplified from the parental plasmid. A 5’ Kozak sequence (gccacc) and a 3’ 2xHA tag were added to saCas9. The modified saCas9 and U6-sgRNA were added to pAAV hSyn via Gibson Assembly. pAAV hSyn saCas9-U6-sgRNA was digested with BsaI and the annealed guides were inserted.

#### Western blot validation of KCNQ3 and KCNQ5 sgRNAs

Crude AAV preparations of Kcnq3 and Kcnq5 sgRNAs inserted in saCas9-U6-sgRNA were prepared. Briefly, HEK293T cells were triple transfected with individual sgRNA-containing pAAV saCas9-U6 plasmids along with pHelper and pAAV-DJ. Three days post-transfection, cells were gently harvested with PBS supplemented with 10mM EDTA, pelleted, and resuspended in AAV freezing media (150 mM NaCl, 20 mM Tris, pH 8.0, 2mM MgCl_2_). Cells were lysed by freeze-thaw 3x and treated with benzonase (50U/mL) for 30 minutes at 37C. Lysates were centrifuged for 15 minutes at 15,000 x g at 4C. Centrifuged lysates were stored at -80 C until use.

Mouse primary hippocampal neurons were generated as previously described^78^. On day *in vitro* 4 (DIV 4), neurons were infected with 5 μL of crude AAV lysate per well. Neuron lysates were harvested on DIV 14-16 and run on a 7.5% SDS-PAGE gel and transferred to nitrocellulose. Nitrocellulose membranes were exposed to the following antibodies: primary: anti-KCNQ3 (1:1000) or anti-KCNQ5 (1:5,000) and anti-β-Actin (1:20,000). The following secondary antibodies were used: Donkey anti-mouse AlexaFluor 680 (1:10,000) and Donkey anti-Rabbit AlexaFluor 790 (1:10,000). Blots were visualized on an Azure Biosystems Western blot imaging system.

#### Purified AAV production, purification, and titration for *in vivo* tests

All the procedures were carried out following Addgene’s AAV Production in HEK293T Cells, and AAV titration by qPCR using SYBR Green technology, with minor modifications. Briefly, pHelper, pAAV2-5 and either pAAV-hSyn-SaCas9-U6-KCNQ3-kozak or pAAV-hSyn-SaCas9-U6-KCNQ5-kozak plasmids were transfected into HEK293T cells using PEIMAX. The molar ratio of pHelper, pAAV2-5, and transfer plasmid was maintained at 1:1:1, while the overall plasmid to PEIMAX ratio was 1:3. After 72 hrs post-transfection, cell lysate and medium underwent precipitation with PEG, followed by purification through an iodixanol gradient. The titer of the purified AAVs was quantified by SYBR qPCR using primers specific to the ITRs (5’-GGAACCCCTAGTGATGGAGTT-3’ and 5’-CGGCCTCAGTGAGCGA-3’).

#### Viral infusion

For brain slice electrophysiology, to localize the PV-positive cells, rAAV_2_-FLEx^loxP^-GFP was co-injected with AAV_5_-hSyn-SaCas9-U6-KCNQ3-kozak and AAV_5_-hSyn-SaCas9-U6-KCNQ5-kozak into the GPe of PV-Cre mice. Acute slice preparations were made 14 days later, as described in the electrophysiology section.

For behavioral tests, 600 nL of a 1:1 volume mix of AAV_5_-hSyn-SaCas9-U6-KCNQ3 and AAV_5_-hSyn-SaCas9-U6-KCNQ5 was injected into the GPe bilaterally. For the no gRNA control, 300 nL of AAV_8_-CMV::NLS-SaCas9-NLS-3xHA was infused into GPe bilaterally. After 21 days post-AAV injection, the mice underwent cocaine CPP and locomotor tests, preceded by an i.p. injection of carnosic acid 30 min before cocaine administration.

#### Single-nucleus RNA-seq

Six 2-month-old mice (3 males and 3 females) were injected i.p. with either cocaine (15 mg/kg) or saline 24 hrs before nuclei isolation. To prepare nuclei, mice were anaesthetized with isoflurane, intracardially perfused with ice-cold RNase free 1x PBS, and then brains were quickly removed. Coronal brain sections (300 μm) were prepared using a vibratome (Leica, VT1200) in ice-cold artificial cerebrospinal fluid (ACSF, as detailed above, and oxygenated with 95% O_2_/5% CO_2_). The GPe tissues were dissected in ice-cold ACSF under a stereomicroscope and were combined by treatment into two samples, cocaine or saline. Tissues were homogenized using a 2 mL Dounce homogenizer (10 passes of A pestle, and 20 passes of B pestle) in 1 mL of lysis buffer (10 mM Tris-HCl, 10 mM NaCl, 3 mM MgCl_2_, 1 mM DTT, 0.1 % Igepal, and 1U/μL RNase inhibitor). Lysates were transferred to a 15 mL tube with an additional 1 mL lysis buffer, kept on ice for 4 min, and centrifuged at 500 x g for 5 min at 4°C. The pellets with nuclei were washed with 1 mL of nuclei wash buffer (NWR, 1x PBS, 1% BSA and 1U/μL RNase inhibitor) before being FACS sorted with DAPI using a BD FACSAria Fusion cell sorter. cDNAs were prepared using the 10x Chromium Single Cell 3′ v3 platform and libraries were sequenced using the Illumina NovaSeq 6000 S4 platform at the Genomics Research and Technology Hub (GRTH) at the University of California, Irvine.

#### Single-nucleus RNA-seq data processing and analysis

##### Single-nucleus RNA-seq alignment and QC

Alignment of reads in the snRNA-seq libraries to the mouse reference transcriptome (mm10, v2020-A) was done using Cellranger (v6.0.1 10X Genomics). The resulting matrices of counts per gene per cell were further processed using Seurat (v4.1.1^79^). Nuclei with the following characteristics were discarded as they were likely debris, damaged, doublets, or outliers: a percentage of reads aligned to mitochondrial genes of over 10, counts of less than 250 or over 4,000, or detected genes of less than 15 or over 2,000. Samples from cocaine-treated and saline-treated mice were collected and processed simultaneously, so batch effects were neither expected nor apparent in the two libraries.

##### Single-nucleus RNA-seq data integration, dimensionality reduction, and clustering

To integrate the two datasets, cocaine-treated and saline-treated, sctransform based normalization was used to normalize, scale, and find variable features within each dataset^80^. Mitochondrial mapping percentage, a potential confounder, was regressed out, and default sctransform parameters were used. 3,000 features were selected as integration features, or features that varied the most across both datasets, and the two datasets were integrated using the IntegrateData command in Seurat, with anchor features being set to our list of 3,000 genes, the normalize method specified as sctransform, and otherwise default parameters. After integration, the standard Seurat workflow for dimensionality reduction, visualization, and clustering was run using the following commands: RunPCA, RunUMAP, FindNeighbors, and FindClusters. For RunUMAP and FindNeighbors the first 20 principal components were used, and for RunUMAP the number of nearest neighbors parameter was set to 20. The resolution parameter for FindClusters was set to 0.3.

##### Annotation of cell types in single-nucleus RNA-seq data

Clusters were annotated as cell types using known marker genes for neurons and glial cells. Oligodendrocyte markers used were Mobp and Mog, OPC markers were Cspg4 and Pdgfra, astrocyte markers were Aqp4 and Gja1, microglia markers were Csf1r and Tmem119, and GABAergic neuron markers were Gad1 and Gad2^27,81–83^. Marker genes for known GABAergic neuron subpopulations in the GPe (Pvalb, Lhx6, Etv1, Npas1, Foxp2, and Meis2) were also included in Figure 5K to help visualize distinctions between the 4 GABAergic subclusters. Marker genes for other cell types including endothelial cells and glutamatergic neurons were also considered during annotation, but these genes were either completely absent from the dataset or expressed very sparsely and weakly, indicating that these cell types were not present in significant amounts.

##### Differential gene expression analysis

Before analysis, raw, filtered counts were normalized using the NormalizeData command in Seurat. Differential gene expression was performed on each cell type cluster using MAST implemented in Seurat. Genes were considered significantly upregulated or downregulated if the adjusted p-value (Bonferroni correction) was less than 0.05, and log_2_FC was either over 0.1 or less than -0.1. Results including statistics are reported in text for genes of interest and in full in Supplementary Table 1.

##### Module detection and analysis using hdWGCNA

hdWGCNA^29^ was performed on the GABAergic cluster of the snRNA-seq data. First, metacells were constructed to combat sparsity in the counts matrix. The hdWGCNA function MetacellsByGroups was applied with metacells being constructed by cell type and by treatment condition (cocaine or saline). Maximum number of shared cells between two metacells was set to 10, and otherwise default parameters were applied. The optimal soft power threshold was determined to be 8 by the hdWGCNA function TestSoftPowers. The standard hdWGCNA workflow for snRNA-seq data was then run for the GABAergic cluster using the following hdWGCNA commands: ConstructNetwork, ModuleEigengenes, ModuleConnectivity, and ModuleExprScore. Two modules were found within the network constructed by hdWGCNA and this network was visualized with the top 3 hub genes labeled using UMAP. This was implemented using 2 hdWGCNA functions: RunModuleUMAP and ModuleUMAPPlot. Module Eigengenes, which are the first principal component of each module, were calculated and we then proceeded to differential module eigengene analysis to test if there were differences in module expression in the cocaine versus saline condition. We did this for the GABAergic cluster as a whole, and for the 4 GABAergic subclusters individually. FindDMEs, a hdWGCNA function, was implemented with the Mann-Whitney U test as a comparison between the cocaine versus saline condition of metacells in each cluster or subcluster. Results (adjusted P value and log_2_FC) were plotted as lollipop plots. Next, gene ontology enrichment analysis of genes in each module was performed to see what biological processes, molecular functions, or cellular components might be associated with each module. This analysis was done, and results visualized using EnrichR (v3.1^84^) and hdWGCNA functions RunEnrichr, GetEnrichrTable, and EnrichrBarPlot sequentially. Enrichment analysis was done using the 2021 versions of the Gene Ontology databases.

#### #Supplemental tables

DGE analysis for each cluster, subcluster, and the dataset as a whole is included in Supplementary Table 1.

Genes expressed in GABAergic neurons and their assigned hdWGCNA module and corresponding kME values is included in Supplementary Table 2.

### QUANTIFICATION AND STATISTICAL ANALYSIS

Statistics for all experiments but sequencing experiments were calculated using GraphPad Prism 9 software. Statistical significance between direct comparisons was assessed by unpaired or paired t-tests. When multiple conditions were compared, one- or two-way ANOVAs were first performed, as appropriate, then t-tests were then performed for each individual comparison. Multiple comparisons corrections were performed when multiple such t-tests were being performed, and significance was assessed using the Holm-Bonferroni method. In conditions where multiple comparisons were performed and the results were still considered significant, asterisks were presented corresponding to the original p-values. Paired t-tests were used in cases with repeat measurements. Dot plots presented throughout the manuscript include a bar representing the mean value for each group. Error bars represent s.e.m. throughout. For all figures, ns P > 0.05, * P ≤ 0.05, ** P ≤ 0.01, *** P ≤ 0.001, **** P ≤ 0.0001.

## References Cited

1. Bromberg-Martin, E.S., Matsumoto, M., and Hikosaka, O. (2010). Dopamine in Motivational Control: Rewarding, Aversive, and Alerting. Neuron 68, 815–834. 10.1016/j.neuron.2010.11.022.

2. Schultz, W. (2007). Multiple Dopamine Functions at Different Time Courses. Annu Rev Neurosci 30, 259–288. 10.1146/annurev.neuro.28.061604.135722.

3. Ungless, M.A., Argilli, E., and Bonci, A. (2010). Effects of stress and aversion on dopamine neurons: Implications for addiction. Preprint, 10.1016/j.neubiorev.2010.04.006 10.1016/j.neubiorev.2010.04.006.

4. Wise, R.A. (2004). Dopamine, learning and motivation. Nat Rev Neurosci 5, 483–494. 10.1038/nrn1406.

5. Kalivas, P.W., and Volkow, N.D. (2005). The neural basis of addiction: a pathology of motivation and choice. Am J Psychiatry 162, 1403–1413. 10.1176/appi.ajp.162.8.1403.

6. Hyman, S.E., Malenka, R.C., and Nestler, E.J. (2006). Neural mechanisms of addiction: the role of reward-related learning and memory. Annu Rev Neurosci 29, 565–598. 10.1146/annurev.neuro.29.051605.113009.

7. Wolf, M.E. (2010). The Bermuda Triangle of cocaine-induced neuroadaptations. Trends Neurosci 33, 391–398. 10.1016/j.tins.2010.06.003.

8. Everitt, B.J., and Robbins, T.W. (2005). Neural systems of reinforcement for drug addiction: From actions to habits to compulsion. Preprint, 10.1038/nn1579 10.1038/nn1579.

9. La Manno, G., Gyllborg, D., Codeluppi, S., Nishimura, K., Salto, C., Zeisel, A., Borm, L.E., Stott, S.R.W., Toledo, E.M., Villaescusa, J.C., et al. (2016). Molecular Diversity of Midbrain Development in Mouse, Human, and Stem Cells. Cell 167, 566–580.e19. 10.1016/j.cell.2016.09.027.

10. Phillips, R.A., Tuscher, J.J., Black, S.L., Andraka, E., Fitzgerald, N.D., Ianov, L., and Day, J.J. (2022). An atlas of transcriptionally defined cell populations in the rat ventral tegmental area. Cell Rep 39. 10.1016/j.celrep.2022.110616.

11. Saunders, A., Macosko, E.Z., Wysoker, A., Goldman, M., Krienen, F.M., de Rivera, H., Bien, E., Baum, M., Bortolin, L., Wang, S., et al. (2018). Molecular Diversity and Specializations among the Cells of the Adult Mouse Brain. Cell 174, 1015–1030.e16. 10.1016/j.cell.2018.07.028.

12. Poulin, J.F., Gaertner, Z., Moreno-Ramos, O.A., and Awatramani, R. (2020). Classification of Midbrain Dopamine Neurons Using Single-Cell Gene Expression Profiling Approaches. Preprint at Elsevier Ltd, 10.1016/j.tins.2020.01.004 10.1016/j.tins.2020.01.004.

13. Poulin, J.F., Zou, J., Drouin-Ouellet, J., Kim, K.Y.A., Cicchetti, F., and Awatramani, R.B. (2014). Defining midbrain dopaminergic neuron diversity by single-cell gene expression profiling. Cell Rep 9, 930–943. 10.1016/j.celrep.2014.10.008.

14. Tian, G., Hui, M., Macchia, D., Derdeyn, P., Rogers, A., Hubbard, E., Liu, C., Vasquez, J.J., Taniguchi, L., Bartas, K., et al. (2022). An extended Amygdala-midbrain circuit controlling cocaine withdrawal-induced anxiety and reinstatement. Cell Rep 39, 1–16. 10.1016/j.celrep.2022.110775.

15. Nestler, E.J. (2005). Is there a common molecular pathway for addiction? Nat Neurosci 8, 1445– 1449. 10.1038/nn1578.

16. Lüscher, C., and Malenka, R.C. (2011). Drug-Evoked Synaptic Plasticity in Addiction: From Molecular Changes to Circuit Remodeling. Neuron 69, 650–663. 10.1016/j.neuron.2011.01.017.

17. Koob, G.F., and Volkow, N.D. (2010). Neurocircuitry of addiction. Neuropsychopharmacology 35, 217–238. 10.1038/npp.2009.110.

18. Beier, K.T., Kim, C.K., Hoerbelt, P., Hung, L.W., Heifets, B.D., Deloach, K.E., Mosca, T.J., Neuner, S., Deisseroth, K., Luo, L., et al. (2017). Rabies screen reveals GPe control of cocaine-triggered plasticity. Nature 549, 345–350. 10.1038/nature23888.

19. Stachniak, T.J., Ghosh, A., and Sternson, S.M. (2014). Chemogenetic Synaptic Silencing of Neural Circuits Localizes a Hypothalamus→Midbrain Pathway for Feeding Behavior. Neuron 82, 797–808. 10.1016/j.neuron.2014.04.008.

20. Schwarz, L.A., Miyamichi, K., Gao, X.J., Beier, K.T., Weissbourd, B., DeLoach, K.E., Ren, J., Ibanes, S., Malenka, R.C., Kremer, E.J., et al. (2015). Viral-genetic tracing of the input–output organization of a central noradrenaline circuit. Nature 524, 88–92. 10.1038/nature14600.

21. Beier, K.T., Steinberg, E.E., Deloach, K.E., Xie, S., Miyamichi, K., Schwarz, L., Gao, X.J., Kremer, E.J., Malenka, R.C., and Luo, L. (2015). Circuit Architecture of VTA Dopamine Neurons Revealed by Systematic Input-Output Mapping. Cell 162, 622–634. 10.1016/j.cell.2015.07.015.

22. Beier, K.T., Gao, X.J., Xie, S., DeLoach, K.E., Malenka, R.C., and Luo, L. (2019). Topological Organization of Ventral Tegmental Area Connectivity Revealed by Viral-Genetic Dissection of Input-Output Relations. Cell Rep 26, 159–167.e6. 10.1016/j.celrep.2018.12.040.

23. Kita, H. (2007). Globus pallidus external segment. Prog Brain Res 160, 111–133. 10.1016/S0079-6123(06)60007-1.

24. Oh, S.W., Harris, J.A., Ng, L., Winslow, B., Cain, N., Mihalas, S., Wang, Q., Lau, C., Kuan, L., Henry, A.M., et al. (2014). A mesoscale connectome of the mouse brain. Nature 508, 207–214. 10.1038/nature13186.

25. Yücel, M., Oldenhof, E., Ahmed, S.H., Belin, D., Billieux, J., Bowden-Jones, H., Carter, A., Chamberlain, S.R., Clark, L., Connor, J., et al. (2019). A transdiagnostic dimensional approach towards a neuropsychological assessment for addiction: an international Delphi consensus study. Preprint at Blackwell Publishing Ltd, 10.1111/add.14424 10.1111/add.14424.

26. McInnes, L., Healy, J., and Melville, J. (2018). UMAP: Uniform Manifold Approximation and Projection for Dimension Reduction. ArXiv 1802.*03426*, 1–51.

27. Hegeman, D.J., Hong, E.S., Hernández, V.M., and Chan, C.S. (2016). The external globus pallidus: Progress and perspectives. Preprint at Blackwell Publishing Ltd, 10.1111/ejn.13196 10.1111/ejn.13196.

28. Schroeder, B.C., Hechenberger, M., Weinreich, F., Kubisch, C., and Jentsch, T.J. (2000). KCNQ5, a Novel Potassium Channel Broadly Expressed in Brain, Mediates M-type Currents. Journal of Biological Chemistry 275, 24089–24095. 10.1074/jbc.M003245200.

29. Morabito, S., Reese, F., Rahimzadeh, N., Miyoshi, E., and Swarup, V. High dimensional co-expression networks enable discovery of transcriptomic drivers in complex biological systems. bioRxiv. 10.1101/2022.09.22.509094.

30. Langfelder, P., and Horvath, S. (2008). WGCNA: An R package for weighted correlation network analysis. BMC Bioinformatics 9. 10.1186/1471-2105-9-559.

31. Morabito, S., Miyoshi, E., Michael, N., Shahin, S., Martini, A.C., Head, E., Silva, J., Leavy, K., Perez-Rosendahl, M., and Swarup, V. (2021). Single-nucleus chromatin accessibility and transcriptomic characterization of Alzheimer’s disease. Nat Genet 53, 1143–1155. 10.1038/s41588-021-00894-z.

32. Tzingounis, A. V., Heidenreich, M., Kharkovets, T., Spitzmaul, G., Jensen, H.S., Nicoll, R.A., and Jentsch, T.J. (2010). The KCNQ5 potassium channel mediates a component of the afterhyperpolarization current in mouse hippocampus. Proc Natl Acad Sci U S A 107, 10232– 10237. 10.1073/pnas.1004644107.

33. Main, M.J., Cryan, J.E., Dupere, J.R.B., Cox, B., Clare, J.J., and Burbidge, S.A. (2000). Modulation of KCNQ2/3 Potassium Channels by the Novel Anticonvulsant Retigabine. Mol Pharmacol 58, 253–262.

34. Kalappa, B.I., Soh, H., Duignan, K.M., Furuya, T., Edwards, S., Tzingounis, A. V., and Tzounopoulos, T. (2015). Potent KCNQ2/3-specific channel activator suppresses in vivo epileptic activity and prevents the development of tinnitus. Journal of Neuroscience 35, 8829–8842. 10.1523/JNEUROSCI.5176-14.2015.

35. Li, S., Choi, V., and Tzounopoulos, T. (2013). Pathogenic plasticity of Kv7.2/3 channel activity is essential for the induction of tinnitus. Proc Natl Acad Sci U S A 110, 9980–9985. 10.1073/pnas.1302770110.

36. Singh, N.A., Westenskow, P., Charlier, C., Pappas, C., Leslie, J., Dillon, J., Anderson, V.E., Sanguinetti, M.C., and Leppert, M.F. (2003). KCNQ2 and KCNQ3 potassium channel genes in benign familial neonatal convulsions: Expansion of the functional and mutation spectrum. Brain 126, 2726–2737. 10.1093/brain/awg286.

37. Singh NA, Charlier, C., Stauffer, D., DuPont, B., Leach, R., Melis, R., Ronen, G., Bjerre, I., Quattlebaum, T., Murphy, J., et al. (1998). A novel potassium channel gene, KCNQ2, is mutated in an inherited epilepsy of newborns. Nat Genet 18, 25–29.

38. Wang, H.S., Pan, Z., Shi, W., Brown, B.S., Wymore, R.S., Cohen, I.S., Dixon, J.E., and McKinnon, D. (1998). KCNQ2 and KCNQ3 potassium channel subunits: Molecular correlates of the M-channel. Science (1979) 282, 1890–1893. 10.1126/science.282.5395.1890.

39. Manville, R.W., Hogenkamp, D., and Abbott, G.W. (2023). Ancient medicinal plant rosemary contains a highly efficacious and isoform-selective KCNQ potassium channel opener. Commun Biol 6. 10.1038/s42003-023-05021-8.

40. Roberts, A.J., Casal, L., Huitron-Resendiz, S., Thompson, T., and Tarantino, L.M. (2018). Intravenous cocaine self-administration in a panel of inbred mouse strains differing in acute locomotor sensitivity to cocaine. Psychopharmacology (Berl) 235, 1179–1189. 10.1007/s00213-018-4834-7.

41. Berridge, K.C., and Robinson, T.E. (1998). What is the role of dopamine in reward: Hedonic impact, reward learning, or incentive salience? Brain Res Rev 28, 309–369. 10.1016/S0165-0173(98)00019-8.

42. Robinson, T.E., and Berridge, K.C. (2000). The psychology and neurobiology of addiction: An incentive-sensitization view. Addiction 95. 10.1080/09652140050111681.

43. Robinson, T.E., and Berridge, K.C. (1993). The neural basis of drug craving: An incentive-sensitization theory of addiction. Brain Res Rev 18, 247–291. 10.1016/0165-0173(93)90013-P.

44. Everitt, B.J., and Robbins, T.W. (2013). From the ventral to the dorsal striatum: Devolving views of their roles in drug addiction. Neurosci Biobehav Rev 37, 1946–1954. 10.1016/j.neubiorev.2013.02.010.

45. Everitt, B.J., and Robbins, T.W. (2005). Neural systems of reinforcement for drug addiction: From actions to habits to compulsion. Nat Neurosci 8, 1481–1489. 10.1038/nn1579.

46. Ito, R., Robbins, T.W., and Everitt, B.J. (2004). Differential control over cocaine-seeking behavior by nucleus accumbens core and shell. Nat Neurosci 7, 389–397. 10.1038/nn1217.

47. Robledo, P., Maldonado-Lopez, R., and Koob, G.F. (1992). Role of Dopamine Receptors in the Nucleus Accumbens in the Rewarding Properties of Cocaine. Ann N Y Acad Sci 654, 509–512. 10.1111/j.1749-6632.1992.tb26015.x.

48. Salmani, B.Y., Lahti, L., Gillberg, L., Jacobsen, J.K., Mantas, I., and Perlmann, T. Transcriptomic atlas of midbrain dopamine neurons uncovers differential vulnerability in a Parkinsonism lesion model. 10.1101/2023.06.05.543445.

49. Peciña, S., and Berridge, K.C. (2005). Hedonic hot spot in nucleus accumbens shell: Where do μ-Opioids cause increased hedonic impact of sweetness? Journal of Neuroscience 25, 11777–11786. 10.1523/JNEUROSCI.2329-05.2005.

50. Al-Hasani, R., McCall, J.G., Shin, G., Gomez, A.M., Schmitz, G.P., Bernardi, J.M., Pyo, C.O., Park, S. Il, Marcinkiewcz, C.M., Crowley, N.A., et al. (2015). Distinct Subpopulations of Nucleus Accumbens Dynorphin Neurons Drive Aversion and Reward. Neuron 87, 1063–1077. 10.1016/j.neuron.2015.08.019.

51. de Jong, J.W., Afjei, S.A., Pollak Dorocic, I., Peck, J.R., Liu, C., Kim, C.K., Tian, L., Deisseroth, K., and Lammel, S. (2019). A Neural Circuit Mechanism for Encoding Aversive Stimuli in the Mesolimbic Dopamine System. Neuron 101, 133–151.e7. 10.1016/j.neuron.2018.11.005.

52. Chen, G., Lai, S., Bao, G., Ke, J., Meng, X., Lu, S., Wu, X., Xu, H., Wu, F., Xu, Y., et al. (2023). Distinct reward processing by subregions of the nucleus accumbens. Cell Rep 42. 10.1016/j.celrep.2023.112069.

53. Yang, H., de Jong, J.W., Tak, Y.E., Peck, J., Bateup, H.S., and Lammel, S. (2018). Nucleus Accumbens Subnuclei Regulate Motivated Behavior via Direct Inhibition and Disinhibition of VTA Dopamine Subpopulations. Neuron 97, 434–449.e4. 10.1016/j.neuron.2017.12.022.

54. Nestler, E.J. (2013). Cellular basis of memory for addiction. Dialogues Clin Neurosci 15, 431– 443.

55. Nestler, E.J. (2014). Epigenetic mechanisms of drug addiction. Neuropharmacology 76, 259–268. 10.1016/j.neuropharm.2013.04.004.

56. Maze, I., and Nestler, E.J. (2011). The epigenetic landscape of addiction. Ann N Y Acad Sci 1216, 99–113. 10.1111/j.1749-6632.2010.05893.x.

57. Nestler, E.J., and Lüscher, C. (2019). The Molecular Basis of Drug Addiction: Linking Epigenetic to Synaptic and Circuit Mechanisms. Neuron 102, 48–59. 10.1016/j.neuron.2019.01.016.

58. Cabana-Domínguez, J., Shivalikanjli, A., Fernàndez-Castillo, N., and Cormand, B. (2019). Genome-wide association meta-analysis of cocaine dependence: Shared genetics with comorbid conditions. Prog Neuropsychopharmacol Biol Psychiatry 94. 10.1016/j.pnpbp.2019.109667.

59. Fernàndez-Castillo, N., Cabana-Domínguez, J., Soriano, J., Sànchez-Mora, C., Roncero, C., Grau-López, L., Ros-Cucurull, E., Daigre, C., Van Donkelaar, M.M.J., Franke, B., et al. (2015). Transcriptomic and genetic studies identify NFAT5 as a candidate gene for cocaine dependence. Transl Psychiatry 5. 10.1038/tp.2015.158.

60. Zhou, Z., Yuan, Q., Mash, D.C., and Goldman, D. (2011). Substance-specific and shared transcription and epigenetic changes in the human hippocampus chronically exposed to cocaine and alcohol. Proc Natl Acad Sci U S A 108, 6626–6631. 10.1073/pnas.1018514108.

61. Teng, L., Fan, L., Peng, Y., He, X., Chen, H., Duan, H., Yang, F., Lin, D., Lin, Z., Li, H., et al. (2019). Carnosic Acid Mitigates Early Brain Injury After Subarachnoid Hemorrhage: Possible Involvement of the SIRT1/p66shc Signaling Pathway. Front Neurosci 13. 10.3389/fnins.2019.00026.

62. Satoh, T., Trudler, D., Oh, C.K., and Lipton, S.A. (2022). Potential Therapeutic Use of the Rosemary Diterpene Carnosic Acid for Alzheimer’s Disease, Parkinson’s Disease, and Long-COVID through NRF2 Activation to Counteract the NLRP3 Inflammasome. Preprint at MDPI, 10.3390/antiox11010124 10.3390/antiox11010124.

63. He, X., Zhang, M., Li, S.T., Li, X., Huang, Q., Zhang, K., Zheng, X., Xu, X.T., Zhao, D.G., and Ma, Y.Y. (2022). Alteration of gut microbiota in high-fat diet-induced obese mice using carnosic acid from rosemary. Food Sci Nutr 10, 2325–2332. 10.1002/fsn3.2841.

64. Liu, Y., Zhang, Y., Hu, M., Li, Y.H., and Cao, X.H. (2019). Carnosic acid alleviates brain injury through NF-κB-regulated inflammation and Caspase-3-associated apoptosis in high fat-induced mouse models. Mol Med Rep 20, 495–504. 10.3892/mmr.2019.10299.

65. Mirza, F.J., Zahid, S., and Holsinger, R.M.D. (2023). Neuroprotective Effects of Carnosic Acid: Insight into Its Mechanisms of Action. Preprint at MDPI, 10.3390/molecules28052306 10.3390/molecules28052306.

66. Barni, M. V., Carlini, M.J., Cafferata, E.G., Puricelli, L., and Moreno, S. (2012). Carnosic acid inhibits the proliferation and migration capacity of human colorectal cancer cells. Oncol Rep 27, 1041–1048. 10.3892/or.2012.1630.

67. Shin, H.B., Choi, M.S., Ryu, B., Lee, N.R., Kim, H.I., Choi, H.E., Chang, J., Lee, K.T., Jang, D.S., and Inn, K.S. (2013). Antiviral activity of carnosic acid against respiratory syncytial virus. Virol J 10. 10.1186/1743-422X-10-303.

68. Greenhill, C. (2011). Liver: Carnosic acid could be a new treatment option for patients with NAFLD or the metabolic syndrome. Preprint, 10.1038/nrgastro.2011.9 10.1038/nrgastro.2011.9.

69. Azhar, M., Zeng, G., Ahmed, A., Dar Farooq, A., Choudhary, M.I., De-Jiang, J., and Liu, X. (2021). Carnosic acid ameliorates depressive-like symptoms along with the modulation of FGF9 in the hippocampus of middle carotid artery occlusion-induced Sprague Dawley rats. Phytotherapy Research 35, 384–391. 10.1002/ptr.6810.

70. Soh, H., Springer, K., Doci, K., Balsbaugh, J.L., and Tzingounis, A. V (2022). KCNQ2 and KCNQ5 form heteromeric channels independent of KCNQ3. 10.1073/pnas.

71. Lein, E.S., Hawrylycz, M.J., Ao, N., Ayres, M., Bensinger, A., Bernard, A., Boe, A.F., Boguski, M.S., Brockway, K.S., Byrnes, E.J., et al. (2007). Genome-wide atlas of gene expression in the adult mouse brain. Nature 445, 168–176. 10.1038/nature05453.

72. Tian, G., Hui, M., Macchia, D., Derdeyn, P., Rogers, A., Hubbard, E., Liu, C., Vasquez, J.J., Taniguchi, L., Bartas, K., et al. (2022). An extended Amygdal-midbrain circuit controlling cocaine withdrawal-induced anxiety and reinstatement. Cell Rep 39, 1–16. 10.1016/j.celrep.2022.110775.

73. Park, J.S., Rhau, B., Hermann, A., McNally, K.A., Zhou, C., Gong, D., Weiner, O.D., Conklin, B.R., Onuffer, J., and Lim, W.A. (2014). Synthetic control of mammalian-cell motility by engineering chemotaxis to an orthogonal bioinert chemical signal. Proc Natl Acad Sci U S A 111, 5896–5901. 10.1073/pnas.1402087111.

74. Lerner, T.N., Shilyansky, C., Davidson, T.J., Evans, K.E., Beier, K.T., Zalocusky, K.A., Crow, A.K., Malenka, R.C., Luo, L., Tomer, R., et al. (2015). Intact-Brain Analyses Reveal Distinct Information Carried by SNc Dopamine Subcircuits. Cell 162, 635–647. 10.1016/j.cell.2015.07.014.

75. Gunaydin, L.A., Grosenick, L., Finkelstein, J.C., Kauvar, I. V., Fenno, L.E., Adhikari, A., Lammel, S., Mirzabekov, J.J., Airan, R.D., Zalocusky, K.A., et al. (2014). Natural neural projection dynamics underlying social behavior. Cell 157, 1535–1551. 10.1016/j.cell.2014.05.017.

76. Concordet, J.P., and Haeussler, M. (2018). CRISPOR: Intuitive guide selection for CRISPR/Cas9 genome editing experiments and screens. Nucleic Acids Res 46, W242–W245. 10.1093/nar/gky354.

77. Hunker, A.C., Soden, M.E., Krayushkina, D., Heymann, G., Awatramani, R., and Zweifel, L.S. (2020). Conditional Single Vector CRISPR/SaCas9 Viruses for Efficient Mutagenesis in the Adult Mouse Nervous System. Cell Rep 30, 4303–4316.e6. 10.1016/j.celrep.2020.02.092.

78. Lloyd, B.A., Han, Y., Roth, R., Zhang, B., and Aoto, J. (2023). Neurexin-3 subsynaptic densities are spatially distinct from Neurexin-1 and essential for excitatory synapse nanoscale organization in the hippocampus. Nat Commun 14. 10.1038/s41467-023-40419-2.

79. Hao, Y., Hao, S., Andersen-Nissen, E., Mauck, W.M., Zheng, S., Butler, A., Lee, M.J., Wilk, A.J., Darby, C., Zager, M., et al. (2021). Integrated analysis of multimodal single-cell data. Cell 184, 3573–3587.e29. 10.1016/j.cell.2021.04.048.

80. Hafemeister, C., and Satija, R. (2019). Normalization and variance stabilization of single-cell RNA-seq data using regularized negative binomial regression. Genome Biol 20. 10.1186/s13059-019-1874-1.

81. Bennett, M.L., Bennett, F.C., Liddelow, S.A., Ajami, B., Zamanian, J.L., Fernhoff, N.B., Mulinyawe, S.B., Bohlen, C.J., Adil, A., Tucker, A., et al. (2016). New tools for studying microglia in the mouse and human CNS. Proc Natl Acad Sci U S A 113, E1738–E1746. 10.1073/pnas.1525528113.

82. Keirstead HS, L.J.B.W. (1998). Response of the oligodendrocyte progenitor cell population (defined by NG2 labelling) to demyelination of the adult spinal cord. Glia 22, 161–170.

83. McKenzie, A.T., Wang, M., Hauberg, M.E., Fullard, J.F., Kozlenkov, A., Keenan, A., Hurd, Y.L., Dracheva, S., Casaccia, P., Roussos, P., et al. (2018). Brain Cell Type Specific Gene Expression and Co-expression Network Architectures. Sci Rep 8. 10.1038/s41598-018-27293-5.

84. Kuleshov, M. V., Jones, M.R., Rouillard, A.D., Fernandez, N.F., Duan, Q., Wang, Z., Koplev, S., Jenkins, S.L., Jagodnik, K.M., Lachmann, A., et al. (2016). Enrichr: a comprehensive gene set enrichment analysis web server 2016 update. Nucleic Acids Res 44, W90–W97. 10.1093/nar/gkw377.

